# Understanding Abundances and Behaviors of Shorebirds in Coastal Louisiana

**DOI:** 10.1101/2025.11.15.688530

**Authors:** T.J. Zenzal, Amanda N. Anderson, Nicholas M. Enwright, Hana R. Thurman, Wyatt C. Cheney, Delaina LeBlanc, Robert C. Dobbs, Brock Geary, J. Hardin Waddle

**Author notes:** Virginia Tech Department of Geosciences, Blacksburg, VA. Louisiana Department of Wildlife and Fisheries, Lafayette, LA. Department of Pathobiology, Wildlife Futures Program, University of Pennsylvania School of Veterinary Medicine, Kennett Square, PA.

## Abstract

Barrier islands provide resources and ecological services that are integral to economic and environmental interests, such as protection of coastal infrastructure and provision of wildlife habitat. Over time, barrier islands may become eroded and experience land loss, which can require management actions to restore island integrity. Barrier island restoration can create or modify habitats, which can impact the organisms depending on them. Our objective was to understand how the abundance and behaviors of a suite of shorebird species responded to restoration and habitat factors at two restored sites in coastal Louisiana (USA). For five focal species, we used abundance from the breeding and non-breeding seasons as well as breeding, foraging, and maintenance behaviors as response variables in boosted regression tree models to determine the importance of various geospatial and remotely sensed predictor variables related to restoration. Across sites and species, remotely sensed variables, particularly a brightness index, tended to be more important than restoration phases as predictors of bird abundance and behavior. Our results suggest that sediment composition, moisture, and vegetative cover are related to shorebird coastal habitat selection, although the direction and strength of relationships differ among these variables and our focal species. Tying these remote sensing metrics to restoration design and management actions can help land managers better understand factors that attract and benefit birds. Additional research can advance understanding in how remote sensing can be used to monitor the availability of functional habitats for shorebirds.

## Introduction

Coastal habitats are some of the most heavily impacted landscapes in the world, and the degradation and loss of coastal habitats are a primary threat to shorebird survival and reproductive success (Atkinson, 2003; Gibson et al., 2018; Iglecia & Winn, 2021; LeDee et al., 2008). Shoreline modifications, human disturbance, and loss of foraging areas are accelerating dramatic habitat and population changes for shorebirds, which can be exacerbated by sea-level rise and climate change (Atkinson, 2003; Brush et al., 2019; Gibson et al., 2018; Smith et al., 2023). As an example, ∼68% of North American shorebird species are in decline, leading to a population reduction of ∼37% since 1970 (Rosenberg et al., 2019). Shorebirds are especially vulnerable to habitat loss because they typically return to sites with specific habitat features and conditions that provide abundant and predictable resources to meet their survival requirements (Brusati et al., 2001; Brush et al., 2019; LeDee et al., 2008; Withers, 2002).

Consequently, shorebird population declines can be severe when high-quality habitat is lost or degraded (Brush et al., 2019; LeDee et al., 2008; Smith et al., 2023).

The Gulf of America (Gulf of Mexico; *hereafter* Gulf) contains highly important habitat for 39 shorebird species during at least some portion of their annual life cycle, and Louisiana (USA) in particular hosts substantial portions of regional and global coastal bird populations (Remsen et al., 2019; Withers, 2002). Nearshore habitats, such as barrier islands and headlands, can play an important role in sustaining robust populations of shorebirds because they provide the required highly dynamic and diverse network of functional habitat types required by shorebirds (Brush et al., 2019; Drake et al., 2001; Withers, 2002). Generally, shorebirds on the Gulf coast use early successional, open landscapes with minimal vegetation for foraging, roosting, and breeding. Shorebird ecology is closely regulated by adequate food resources and roosting habitats. For example, shorebirds typically select safe roost sites with abundant food resources and low commuting costs among functional habitat types (Brush et al., 2019; Iglecia & Winn, 2021; Noel & Chandler, 2008). Food resources likely influence foraging distribution and site fidelity, and extensive intertidal zones with shallow slopes, saturated substrates, and sediment influxes support high concentrations of invertebrates required by shorebirds (Brush et al., 2019; Iglecia & Winn, 2021; McIntyre & Heath, 2011; Schulz & Leberg, 2019). Additionally, roost sites located above the high tide line with open views and wind breaks are critical for rest, predator avoidance, digestion, and other maintenance behaviors (Brush et al., 2019; Farrell et al., 2016).

Shorebird survival is maximized when high-quality foraging areas like intertidal flats and unvegetated mudflats are near roost sites like beach-backshore and sandflats, minimizing daily energy expenditure (Brush et al., 2019; Farrell et al., 2016; Iglecia & Winn, 2021). In addition to complex habitat requirements, shorebird populations are limited by adult mortality as well as generally low fecundity and productivity in the breeding season, which contributes to shorebirds being especially susceptible to population declines (Iglecia & Winn, 2021; Rosenberg et al., 2019; Smith et al., 2023).

Whereas barrier islands primarily serve to protect the coastline, they also provide important habitat across a shorebird’s life cycle. Wilson’s Plover (*Anarhynchus wilsonia*) and American Oystercatcher (*Haematopus palliatus*), for example, select nest sites within early successional habitats that include unvegetated washover fans, backshore zones of beaches, and low elevation dunes with sandy to shell substrate, typically above the high tide line (American Oystercatcher Working Group et al., 2020; DeRose-Wilson et al., 2013; Zdravkovic et al., 2023). Additionally, proximity to cover and food resources is an important predictor of reproductive success and nest site selection as breeding birds require access to these resources during the nesting or young-rearing stages (Bergstrom, 1988; DeRose-Wilson et al., 2013; Lauro & Burger, 1989; Ray, 2011; Schulte & Simons, 2014). Being able to remotely assess the functional habitats needed by shorebirds can provide a powerful and cost-effective management tool without potentially disturbing populations with traditional ground-based avian surveys.

Coastal restoration and habitat management are tools that land managers can use to mitigate the effects of direct habitat loss and changing environmental conditions on shorebird dynamics. Although restoration can increase the total amount of land area, coastal engineering may alter coastal processes that create and maintain intertidal as well as early successional habitats — directly affecting habitat available to shorebirds.

For example, natural processes that form complex geomorphology can be absent and affect features like shoreline structure, sediment characteristics, invertebrate assemblages, and microhabitats (Atkinson, 2003; Farrell et al., 2016; Iglecia & Winn, 2021; McIntyre & Heath, 2011). Of course, some of these impacts may be temporary, and longevity of restoration impacts on habitats is not well understood. Despite uncertainty related to barrier island restoration, this management is ongoing along the northern Gulf coast as state and federal agencies work to develop natural protections for life and property from severe tropical cyclone impacts.

As engineered coastal habitats become increasingly widespread, there is a growing need to dovetail protecting human life and property with designing projects that maximize benefits for target wildlife populations. Engineering projects aim to achieve stable states with predictable results, but project outcomes often differ from initial expectations. Project success related to targeted wildlife populations can be measured in terms of providing natural processes that maintain a diversity of habitats, food resources, and bird species (Atkinson, 2003; Iglecia & Winn, 2021; Withers, 2002).

Considering that Louisiana’s coastal habitats are important for sustaining shorebird populations, including Piping Plover (*Charadrius melodus*) and Red Knot (*Calidris canutus*) recovery, baseline monitoring that incorporates response metrics related to breeding, roosting, and foraging habitat can be used to evaluate restoration performance and reduce uncertainty when planning for restoration that benefits people and shorebirds. Our objective was to use machine learning to determine how abundance and behavior of focal shorebird species (Red Knot, Piping Plover, Snowy Plover [*Anarhynchus nivosus*], Wilson’s Plover, and American Oystercatcher) are related to coastal restoration and associated habitat features.

## Methods

### Study Sites

To address our objective, we worked at two locations along the northern Gulf in coastal Louisiana, USA that were slated for restoration by the Coastal Protection and Restoration Authority of Louisiana. One site is the Caminada Headland (29.139°N, - 90.134°W; Figure 1), which is located within Lafourche and Jefferson parishes, to the south and east of Port Fourchon, Louisiana. The other site is Whiskey Island (29.046°N, -90.840°W; Figure 2), which is south of Cocodrie, Louisiana in Terrebonne Parish and one of four barrier islands that comprises the Isle Dernieres Barrier Islands Refuge (IDBIR).

**Figure 1.**
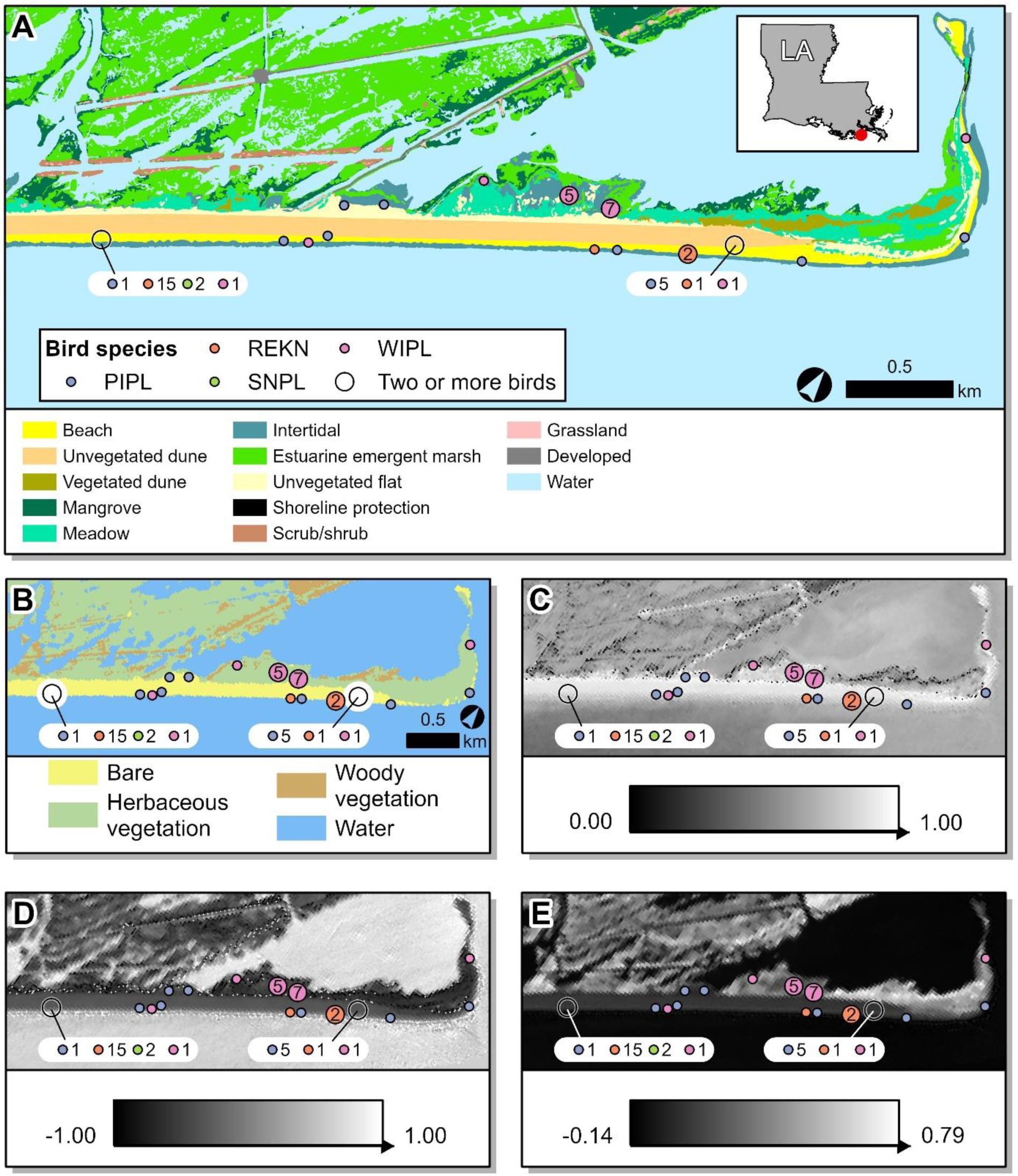
Examples from the avian survey of Caminada Headland, Louisiana, USA on September 6, 2017 shown with closest habitat maps and spectral index values. Area shown includes a subset of the headland near Caminada Pass. Bird species are differentiated by color, with larger circles representing multiple birds that were clustered at or near the same location (≤118 m apart). For clusters where three or more different species were present, a breakout is included to show the number of individuals by species. Bird species include Piping Plover (PIPL), Red Knot (REKN), Snowy Plover (SNPL), and Wilson’s Plover (WIPL). Unless specified, all maps and spectral indices were developed from Landsat 8 satellite imagery captured on September 8, 2017. a) Bird points and detailed habitat map developed using National Agriculture Imagery Program (NAIP) 1-m orthoimagery from September 8–12, 2017, b) Bird points and multitemporal, satellite-based habitat map, c) Bird points and Brightness Index (BI), d) Bird points and Modified Normalized Difference Water Index (MNDWI), e) Bird points and Modified Soil Adjusted Vegetation Index (MSAVI).

**Figure 2.**
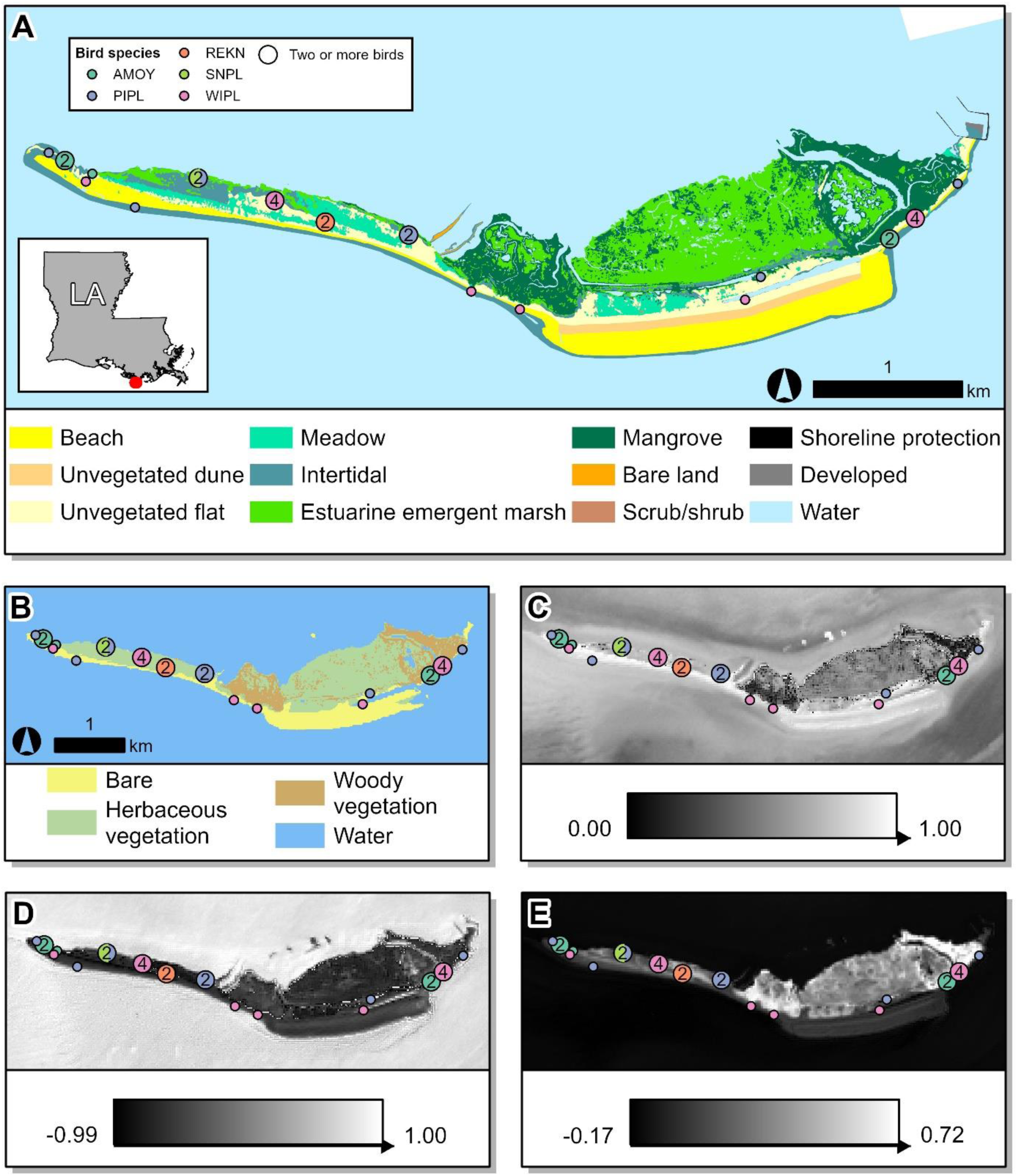
Example locations from the avian survey of Whiskey Island, Louisiana, USA on September 7, 2017 shown with closest habitat maps and spectral index values. Bird species are differentiated by color, with larger circles representing multiple birds that were clustered at or near the same location (≤22 m apart). Bird species include Piping Plover (PIPL), Red Knot (REKN), Snowy Plover (SNPL), and Wilson’s Plover (WIPL). Unless specified, all maps and indices were developed from Landsat 8 satellite imagery captured on September 8, 2017. a) Bird points and detailed habitat map developed using National Agriculture Imagery Program (NAIP) 1-m orthoimagery from September 10–11, 2017, b) Bird points and multitemporal, satellite-based habitat map, c) Bird points and Brightness Index (BI), d) Bird points and Modified Normalized Difference Water Index (MNDWI), e) Bird points and Modified Soil Adjusted Vegetation Index (MSAVI).

Caminada Headland extends ∼21 km and consists of beach and intertidal landcovers that span from Belle Pass eastward to Caminada Pass on Elmers Island (Figure 1). Caminada Headland consists of the following habitat types: 1) intertidal and beach habitats facing the Gulf, 2) unvegetated dune and unvegetated flat habitats making up the interior of the headland, and 3) estuarine emergent marsh, black mangrove (*Avicennia germinans*), intertidal, meadow, and scrub/shrub habitats along the bay side of the headland (Table S1). Over the last 100 years (1880s–2015), the shoreline of the headland has eroded by an average of 12 m/yr (Byrnes et al., 2018). There have been two restoration projects, which occurred in 2015 and 2016, on Caminada Headland (refer to LeBlanc et al., [2023] for more details). The goal of Caminada Headland’s restoration was to sustain coastal habitat, recreate littoral sand transport, and diminish storm impacts on infrastructure (Folse & Lee, 2016). These restoration projects established ∼4.29 km^2^ of beach and dune habitat by dredging ∼8.8 million m^3^ of sediment from offshore sources (Coastal Protection and Restoration Authority of Louisiana, 2018).

Whiskey Island (Figure 2) is composed of: 1) beach and intertidal habitats facing the Gulf, 2) intertidal, mangrove, and estuarine emergent marsh habitats facing Caillou Bay to the north, and 3) spits of beach, intertidal, and unvegetated flat habitats at the island’s east and west ends. Within the interior portion of the island there are nontidal habitats, which include vegetated dune, meadow, and unvegetated flats (refer to Enwright et al., [2020] for habitat definitions). Throughout the island, inundation or flooding from tides, rain, or other events create ephemeral pools or tidal inlets within low-lying areas as well as areas where mangrove or estuarine emergent marsh habitats transition to other habitat types. Over the span of ∼100 years (1855–2015), the shoreline has eroded by an annual average of approximately 16 m/yr (Byrnes et al., 2018). In 2018, Whiskey Island underwent its most recent restoration project (refer to LeBlanc et al. [2023] for more details). Restoration of Whiskey Island involved dredging ∼8 million m^3^ of sediment, which was sourced from offshore areas, to produce 3.86 km^2^ of beach, dune, and marsh habitats (Coastal Protection and Restoration Authority of Louisiana, 2018). After sediment deposition, vegetation plantings were implemented to stabilize the mainland shore and mitigate erosion by attenuating wave and tidal energy (Coastal Protection and Restoration Authority of Louisiana, 2018). Following restoration at Whiskey Island, land area expanded from 3.03 km^2^ in 2015 to 4.78 km^2^ in 2019.

### Field Surveys

While field survey methods are outlined in LeBlanc and colleagues (2023) and associated data are presented in Zenzal and colleagues (2023), a brief summary is provided here, for reference. Avian ecologists from the Barataria-Terrebonne National Estuary Program conducted bird surveys along the Caminada Headland at least semimonthly from January 2013 to June 2019. Surveys adhered to the International Shorebird Survey (ISS) protocol and were conducted ± 2 days of the designated census dates (U.S. Fish and Wildlife Service, 2015). Survey gaps during the sampling period were due to logistical limitations, including adverse weather and personnel restraints. All suitable habitats (e.g., beach, unvegetated flat, intertidal, meadow) were surveyed to locate focal species that included Piping Plover, Red Knot, Snowy Plover, and Wilson’s Plover. Surveys for focal species occurred from west Belle Pass eastward to Caminada Pass (∼22.5 km). The area was split into 4–5 sections, with each section surveyed on foot by 1–4 observers in a single day. For each focal species, observers documented: 1) number of individuals, 2) leg-band combinations and alphanumeric flag codes, 3) geographic coordinates of the observer, 4) geographic subarea within the entire survey area (West Belle Pass, West Beach, East Beach, Bayou, and Elmers Island), 5) location (Gulf, bay, or interior), and 6) behavior (e.g., foraging, roosting, breeding). Given that the geographic coordinates reflect the position of an observer and not a bird, we used a Euclidean allocation analysis to project each bird’s position in space. The projected location of a bird was calculated by moving the geographic coordinates of the observer to the nearest pixel of habitat type, based on Euclidean distance, that matched a bird’s documented location (Gulf, bay, or interior).

At Whiskey Island, avian ecologists from the U.S. Geological Survey (USGS) conducted bi-weekly surveys to monitor the non-breeding avian community (August–February) and weekly surveys to monitor the avian breeding community (March–July). During the sampling period (2012–2020), some surveys were missed due to challenges with data collection at a remote site (e.g., weather, personnel availability), funding lapses, or the COVID-19 pandemic. Surveys were completed within a single day by a pair of observers, each independently assessing the focal species’ habitat by foot.

These focal species included Piping Plover, Snowy Plover, Wilson’s Plover, American Oystercatcher, and Red Knot. For each individual of a focal species observed, surveyors recorded: 1) number of individuals, 2) leg-band combinations and alphanumeric flag codes, 3) geographic coordinates of the bird, 4) location (e.g., Gulf, bay, interior), 5) behavior (e.g., foraging, roosting, breeding), and 6) tidal stage (i.e., high, low, rising, falling).

We grouped the location of a focal species observations into Gulf, bay, or interior. The Gulf-facing portion of the island is located on the southern side, where the beach zone spans the area between the low and high-water lines. Gulf areas have little to no vegetation and can experience high-energy shorelines, wave action, and storm surge. Bay portions of the island are on the north side where habitats, such as intertidal flats, marshes, and mangroves are abutting estuarine open water with typically low wave energy and are exposed to changing water levels and storm surge. Interior portions of the island are beyond the high-water line, which includes vegetated or unvegetated dunes, unvegetated barrier flats, and shrubby areas.

For the analyses, we grouped specific behaviors into three overarching behavioral categories: foraging, maintenance, and breeding. Birds were categorized as foraging if they were walking while pecking at substrate, devouring prey, or transporting prey. Birds were categorized as engaging in maintenance behaviors when they were observed preening, roosting, or loafing. Preening is defined as birds conducting feather maintenance. Roosting was noted when a bird had their beak tucked behind their head. Loafing is defined as a bird in standing or sitting posture, apparently idle and not related to another behavior. Similar to Zenzal and colleagues (2025), birds were categorized as breeding when displaying at least one of the following: 1) courtship or paired, 2) territoriality (e.g., territorial postures, vocalizations, aggression), 3) defense (e.g., alarm-calling/agitation, mobbing, broken-wing display), 4) nest construction (i.e., scraping), 5) evidence of eggs or young, and 6) adult carrying food or brooding young. We noted the behavior first observed unless it was apparent that due to the presence of the surveyor a bird became vigilant or defensive, which required the surveyor to move past the bird, allow a short acclimation period, and then document the first non-vigilant or defensive behavior. In cases where defensive or vigilance persisted past the acclimation period (∼5 mins), we marked that individual’s behavior as “defense.”

### Geospatial Analyses

We used a variety of geospatial predictor variables to understand how remotely sensed habitat metrics may be related to avian abundance and behavior. We derived geospatial variables from: 1) 1-m orthoimagery-based habitat maps, 2) inundation classes and distance to water from orthoimagery-based maps, 3) satellite-based habitat maps developed from Landsat 8 surface reflectance imagery (USGS, 2021) and Sentinel-2 surface reflectance imagery (ESA, 2015), and 4) spectral indices from these satellite imagery. The spectral indices included the brightness index (BI; a spectral index used to indicate brightness of soil and substrate; range: 0–1; Escadafal et al., 1989), modified soil adjusted vegetation index (MSAVI; a spectral index used to indicate vegetation greenness; range: -1–1; Qi et al., 1994), and modified normalized difference water index (MNDWI; a spectral index used to indicate wetness and water bodies; range: -1–1; Xu, 2006). BI provides information on vegetative cover and moisture levels with dense vegetation and wet substrate having a lower value. MSAVI provides information on the presence and amount of green vegetation. MNDWI provides information on substrate moisture and presence of water. Products were developed using methodology outlined in Enwright and coauthors (2020) or Thurman and coauthors (2023), with supplemental information providing additional detail.

### Statistical Analyses

To meet our objective of understanding how habitat covariates are associated with avian abundance and behaviors (i.e., foraging, breeding, and maintenance) for each focal species (i.e., American Oystercatcher [Whiskey Island only], Piping Plover, Red Knot, Snowy Plover, and Wilson’s Plover), we employed a machine learning approach known as boosted regression trees (BRT). Machine learning approaches provide an advantage over traditional analytical tests by simultaneously: 1) selecting relatively important predictor variables from a potentially large dataset, 2) identifying functional forms of associations between response and predictor variables, and 3) incorporating a high number of interactions among predictor variables (Elith et al., 2008). Additionally, BRT results are relatively straightforward to interpret because the relative influence of predictor variables sum to 100% and illustrate stronger effects on the response variables when values are higher (Elith et al., 2008).

BRTs can handle correlated predictor variables when generating accurate predictions, but including them may cause issues when interpreting models as the relative influence of two highly correlated variables would likely be split, suggesting a weaker effect of each (Elith et al., 2008). Therefore, we tested for correlated variables using a Spearman rank-order correlation coefficient analysis (numeric variables) or Cramer’s V (categorical variables) and removed what we believed to be the less biologically meaningful variable from pairs that appeared to be highly correlated (Spearman’s: rho > |0.85|; Cramer’s: V > 0.85). After accounting for highly correlated variables, we included the following predictor variables into each model: BI (Figure 1 and 2), restoration phase (i.e., pre, during, or post-restoration), MNDWI (Figure 1 and 2), MSAVI (Figure 1 and 2), orthoimagery-based detailed habitat classes (Table S1; Figure 1 and 2), tidal stage (Whiskey Island only), inundation class (Table S2), location (Gulf, bay, or interior), site (Bayou, East Beach, Elmers Island, West Belle Pass, West Beach; Caminada Headland only), distance to nearest water (m), and satellite-based habitat classes (bare, herbaceous, water, woody; Figure 1 and 2). Our species-specific response variables included daily abundance of birds as well as the presence or absence of breeding, foraging, or maintenance behaviors for each individual bird observation. We were only able to analyze breeding behavior for American Oystercatcher (Whiskey Island only) and Wilson’s Plover since they were the only two focal species breeding at our study sites. We followed the methods and script provided in Elith and coauthors (2008) to run all the models, which involved using a tree complexity of 3 and adjusting the learning rate (range: 0.00004–0.5) to produce between 1,000 and 10,000 trees. Models of avian abundance used a Poisson distribution and models of behavior used a Bernoulli distribution. Due to issues with model singularity or convergence in some instances, we adjusted the bag fraction (range: 0.5–0.9) for models to run correctly. We interpreted predictor variables with >10% relative influence given that by random chance alone any variable could have a 10% relative influence (100% relative influence divided by 10 predictor variables = 10% relative influence/predictor variable; Müller et al., 2013; Schofield et al., 2018). All analyses were performed in the R statistical language (R Core Team, 2021) using packages “dismo” (Hijmans et al., 2021), “dplyr” (Wickham et al., 2022), “data.table” (Dowle & Srinivasan, 2021), “gbm” (Greenwell et al., 2020), “raster” (Hijmans, 2022), “sp” (Bivand et al., 2013; Pebesma & Bivand, 2005), “rcompanion” (Mangiafico, 2025), and “corrplot” (Wei & Simko, 2021).

## Results

### Caminada Headland

At Caminada Headland, we identified predictor variables that were important across all species in the various models. Distance to water was important in the non-breeding abundance, foraging, and maintenance models. We also found BI, MNDWI, and MSAVI to be important across models analyzing breeding abundance as well as all behavioral models (breeding, foraging, and maintenance). We did not find restoration phase to be an important predictor variable in any analysis at Caminada Headland.

#### Piping Plover

We were able to model non-breeding abundances as well as foraging and maintenance behaviors of Piping Plover at Caminada Headland. We found meaningful associations between non-breeding abundance and distance to water (21.1%), orthoimagery-based detailed habitat classes (19.8%), location (16.3%), and MNDWI (15.5%; CV: 0.46 ± 0.03 [this and all following presented as mean ± standard error]; Figure 3A). In the behavioral analysis, we found foraging behaviors to be meaningfully associated with location (29.8%), BI (17.7%), MSAVI (14.3%), distance to water (12.9%), and MNDWI (12.8%; CV: 0.88 ± 0.01; Figure 3B). The analysis of maintenance activities found meaningful associations with location (31.1%), BI (18.3%), MSAVI (14.0%), distance to water (12.6%), and MNDWI (12.1; CV: 0.88 ± 0.00; Figure 3C). Further information on the direction of relationships is described in the Supplemental Information along with full model outputs (Figures S1–S3).

**Figure 3.**
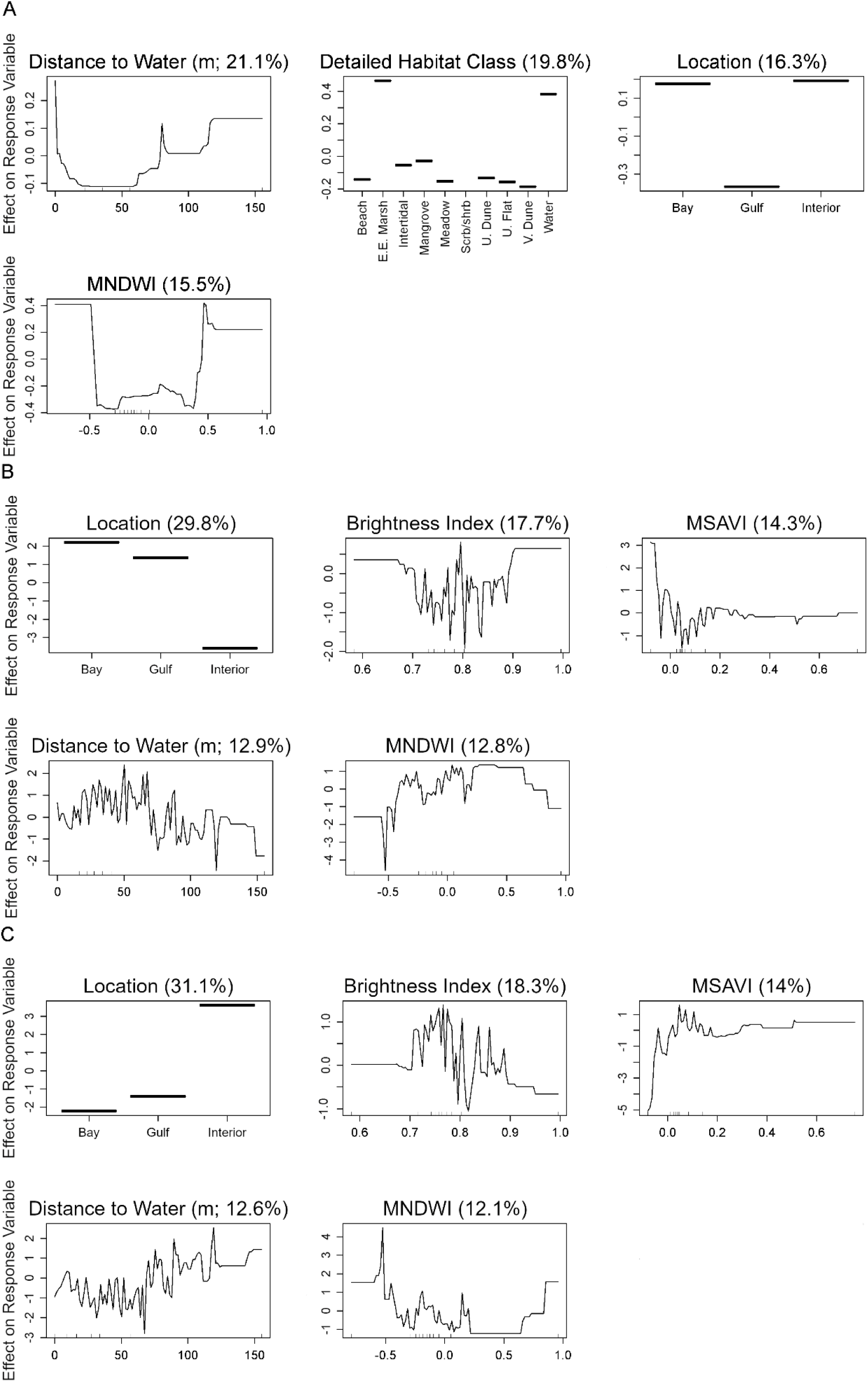
Partial dependence plots of variables related to Piping Plover (A) abundance, (B) foraging behaviors, and (C) maintenance behaviors during the non-breeding season at Caminada Headland, Louisiana, USA from a boosted regression tree model. Y-axes are centered to have a zero mean over the data distribution. The relative influence (percent) of each predictor variable is shown in parentheses. Rug plots along the X-axis of each continuous variable plot illustrates the distribution of the data. Only variables with greater than 10% relative influence are shown here (refer to Supplementary Information for full model output). Abbreviated labels on the X-axis of Detailed Habitat Class include Estuarine Emergent Marsh (E.E. Marsh), Scrub/Shrub (Scrb/Shrb), Unvegetated Dune (U. Dune), Unvegetated Flat (U. Flat), and Vegetated Dune (V. Dune). Abbreviated labels on the X-axis of Inundation Class include Intertidal Unvegetated (Intertidal Unveg.) and Intertidal Vegetated (Intertidal Veg.). Abbreviated labels on the X-axis of Site include East Beach (E. Beach), West Belle Pass (W. Belle Pass), and West Beach (W. Beach). MNDWI indicates Modified Normalized Difference Water Index and MSAVI indicates Modified Soil Adjusted Vegetation Index.

#### Red Knot

We were able to model non-breeding abundances as well as foraging and maintenance behaviors of Red Knot at the Caminada Headland. When analyzing abundances during the non-breeding season, we found only two variables to have a meaningful association with the response: distance to water (60.4%) and BI (29.0%; CV: 0.28 ± 0.06; Figure 4A). In our behavioral analysis, we found foraging to be affiliated with distance to water (23.5%), BI (17.5%), MSAVI (14.7%), MNDWI (12.4%), and location (12.2%; CV: 0.96 ± 0.00; Figure 4B). Investigating maintenance activities in our analysis, we found meaningful associations between the response variable and distance to water (26.4%), BI (16.8%), MSAVI (14.7%), MNDWI (12.0%), and location (11.9%; CV: 0.96 ± 0.01; Figure 4C). Further information on the direction of relationships is described in the Supplemental Information along with full model outputs (Figures S4–S6).

**Figure 4.**
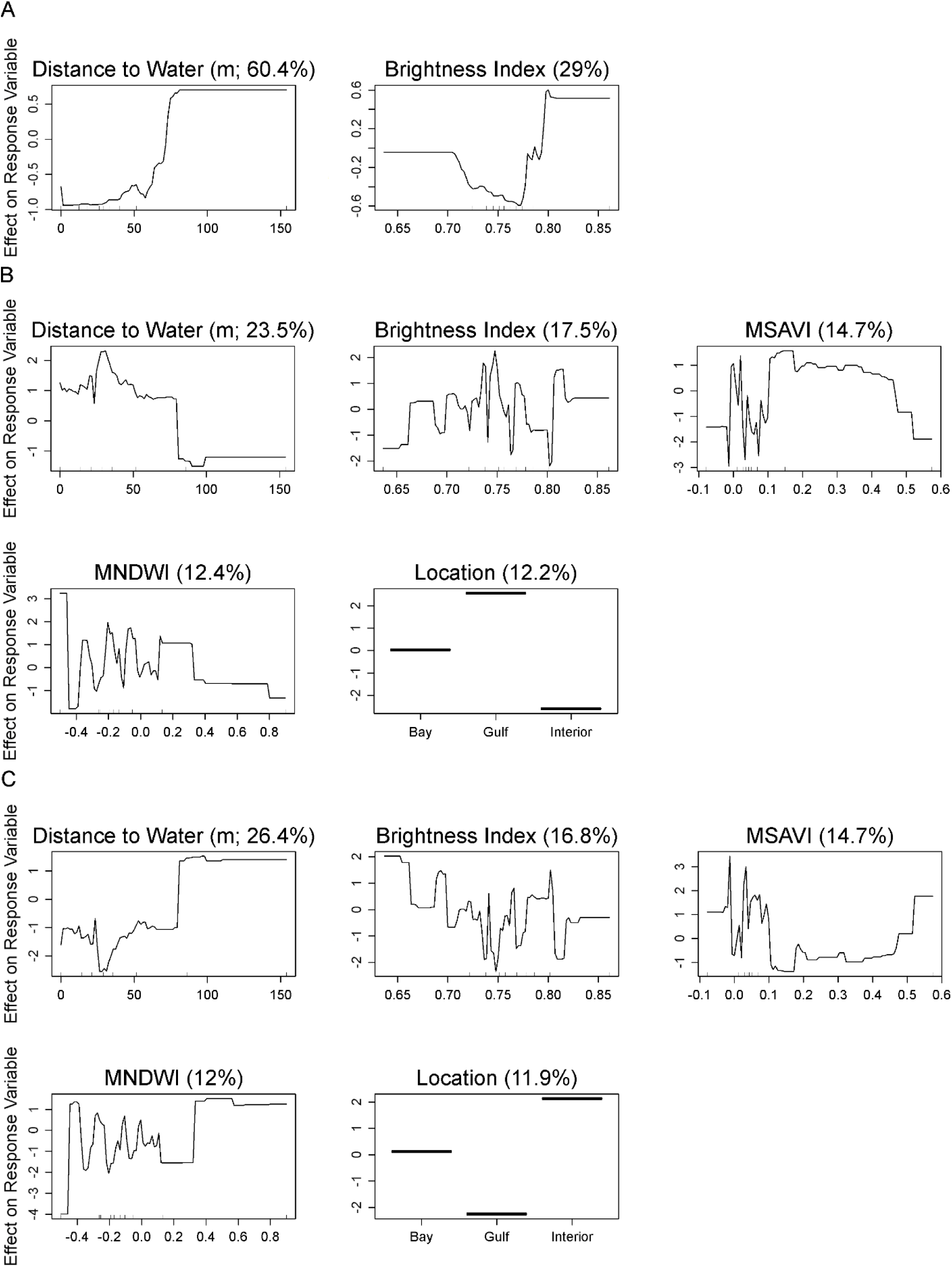
Partial dependence plots of variables related to Red Knot (A) abundance, (B) foraging behaviors, and (C) maintenance behaviors during the non-breeding season on Caminada Headland, Louisiana, USA from a boosted regression tree model. Y-axes are centered to have a zero mean over the data distribution. The relative influence (percent) of each predictor variable is shown in parentheses. Rug plots along the X-axis of each continuous variable plot illustrates the distribution of the data. Only variables with greater than 10% relative influence are shown here (refer to Supplementary Information for full model output). Abbreviated labels on the X-axis of Detailed Habitat Class include Estuarine Emergent Marsh (E.E. Marsh), Scrub/Shrub (Scrb/Shrb), Unvegetated Dune (U. Dune), Unvegetated Flat (U. Flat), and Vegetated Dune (V. Dune). Abbreviated labels on the X-axis of Inundation Class include Intertidal Unvegetated (Intertidal Unveg.) and Intertidal Vegetated (Intertidal Veg.). Abbreviated labels on the X-axis of Site include East Beach (E. Beach), West Belle Pass (W. Belle Pass), and West Beach (W. Beach). MNDWI indicates Modified Normalized Difference Water Index and MSAVI indicates Modified Soil Adjusted Vegetation Index.

#### Snowy Plover

We were able to model non-breeding abundances as well as foraging and maintenance behaviors of Snowy Plover at Caminada Headland. When analyzing non-breeding abundance, we found abundance to be meaningfully related to MNDWI (29.8%), MSAVI (24.3%), distance to water (12.3%), orthoimagery-based detailed habitat maps (11.0%), and BI (11.0%; CV: 0.28 ± 0.08; Figure 5A). For the behavioral analysis, we found foraging to be meaningfully associated with BI (20.0%), location (17.3%), distance to water (17.1%), MSAVI (14.3%), and MNDWI (12.5%; CV: 0.56 ± 0.3; Figure 5B). The analysis of maintenance activities found BI (22.8%), distance to water (19.1%), MSAVI (12.7%), MNDWI (12.4%), and location (11.8%) to have a strong association with the response (CV: 0.54 ± 0.03; Figure 5C). Further information on the direction of relationships is described in the Supplemental Information along with full model outputs (Figures S7–S9).

**Figure 5.**
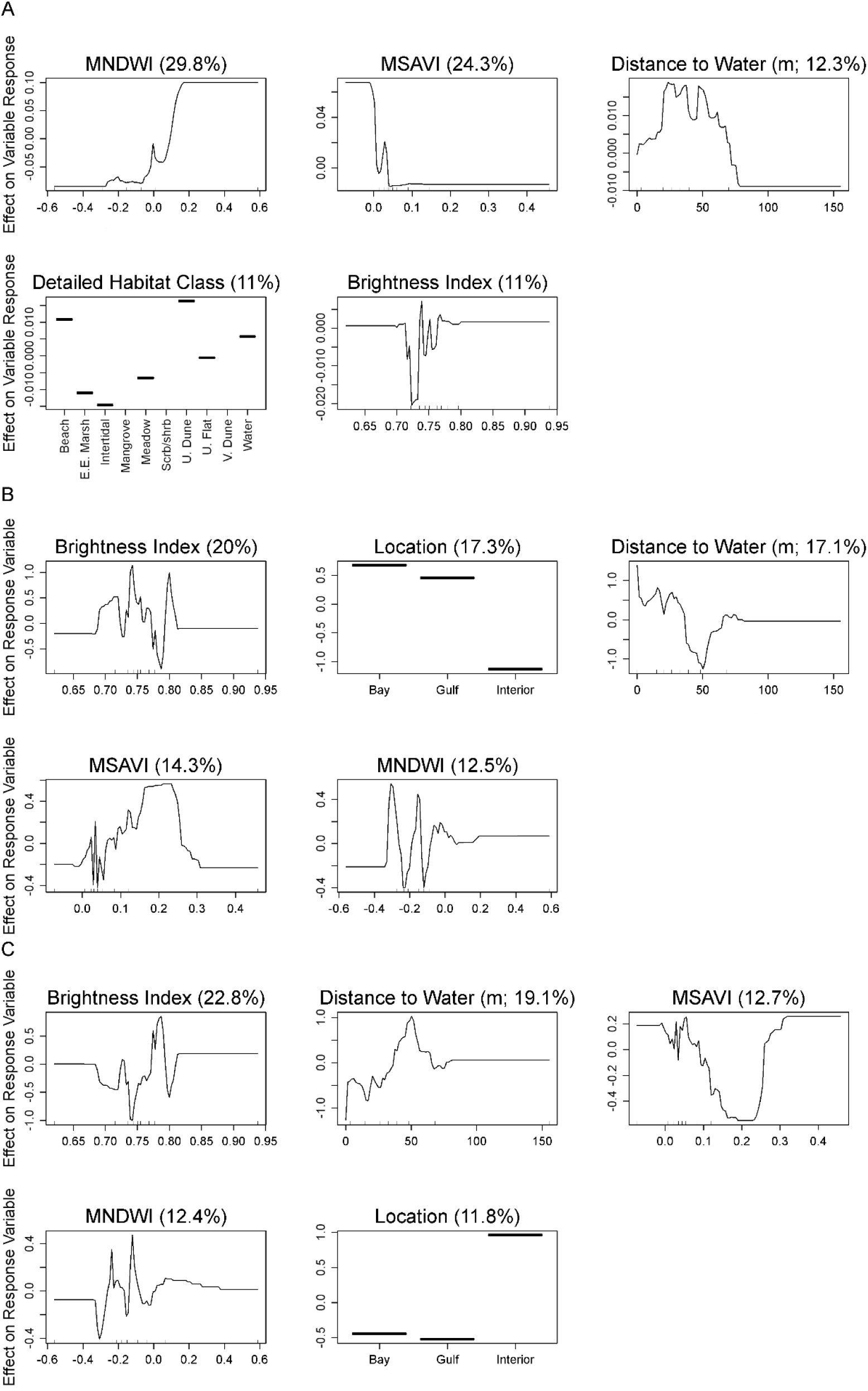
Partial dependence plots of variables related to Snowy Plover (A) abundance, (B) foraging behaviors, and (C) maintenance behaviors during the non-breeding seasons on Caminada Headland, Louisiana, USA from a boosted regression tree model. Y-axes are centered to have a zero mean over the data distribution. The relative influence (percent) of each predictor variable is shown in parentheses. Rug plots along the X-axis of each continuous variable plot illustrates the distribution of the data. Only variables with greater than 10% relative influence are shown here (refer to Supplementary Information for full model output). Abbreviated labels on the X-axis of Detailed Habitat Class include Estuarine Emergent Marsh (E.E. Marsh), Scrub/Shrub (Scrb/Shrb), Unvegetated Dune (U. Dune), Unvegetated Flat (U. Flat), and Vegetated Dune (V. Dune). Abbreviated labels on the X-axis of Inundation Class include Intertidal Unvegetated (Intertidal Unveg.) and Intertidal Vegetated (Intertidal Veg.). Abbreviated labels on the X-axis of Site include East Beach (E. Beach), West Belle Pass (W. Belle Pass), and West Beach (W. Beach). MNDWI indicates Modified Normalized Difference Water Index and MSAVI indicates Modified Soil Adjusted Vegetation Index.

#### Wilson’s Plover

We were able to model breeding and non-breeding abundances as well as breeding, foraging, and maintenance behaviors of Wilson’s Plover at Caminada Headland. During the breeding season, we found abundance to be strongly associated with BI (32.8%), MNDWI (18.8%), MSAVI (16.5%), distance to water (15.3%), and orthoimagery-based detailed habitat classes (11.6%; CV: 0.30 ± 0.03; Figure 6A). During the non-breeding season, MSAVI (30.4%), distance to water (24.8%), BI (14.5%), and MNDWI (12.8%) were meaningfully affiliated with abundance (CV: 0.46 ± 0.04; Figure 6B). Analyzing the occurrence of breeding behaviors found MSAVI (26.1%), location (23.9%), site (13.5%), BI (13.1%), and MNDWI (12.0%) to be meaningfully associated with the response (CV: 0.88 ± 0.02; Figure 6C). Taking foraging behaviors into consideration, we found foraging to be strongly linked to MNDWI (19.0%), MSAVI (18.9%), BI (16.7%), location (15.4%), and distance to water (14.6%; CV: 0.80 ± 0.01; Figure 6D). When taking into consideration maintenance activities, we found MSAVI (21.2%), location (18.6%), MNDWI (17.4%), BI (16.2%), and distance to water (14.2%) to have meaningful associations (CV: 0.82 ± 0.01; Figure 6E). Further information on the direction of relationships is described in the Supplemental Information along with full model outputs (Figures S10–S14).

**Figure 6.**
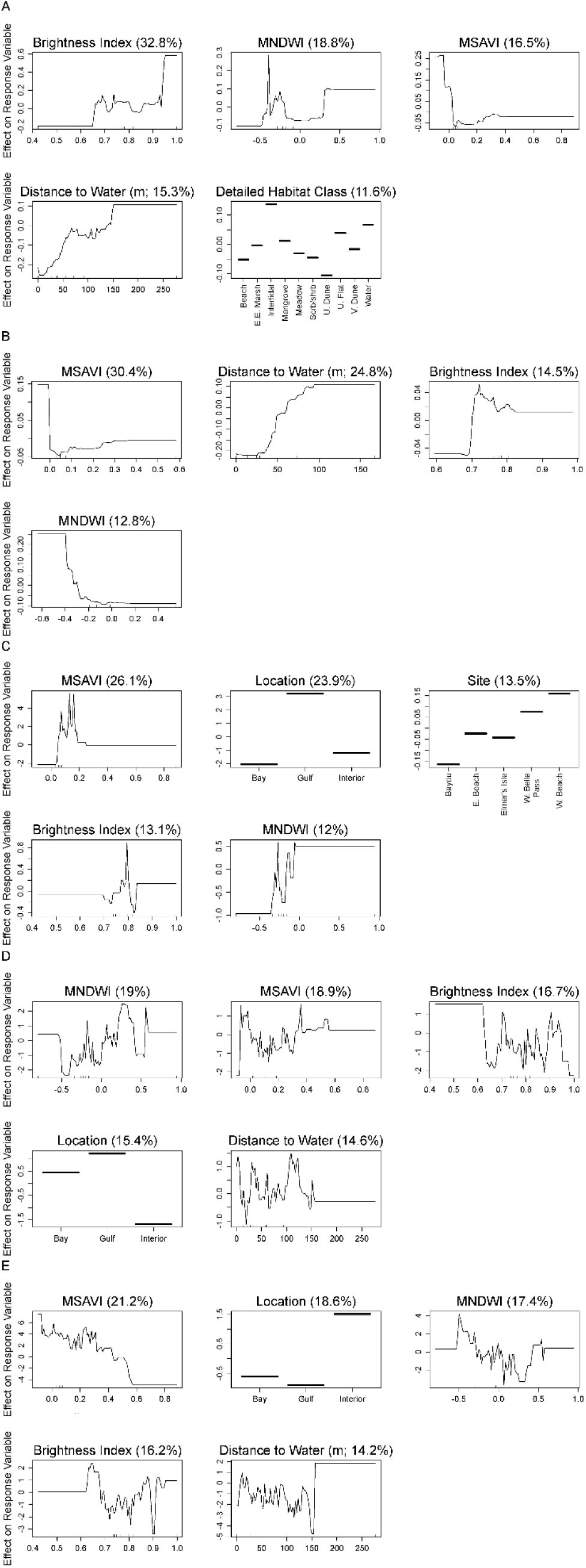
Partial dependence plots of variables related to Wilson’s Plover (A) breeding abundance, (B) non-breeding abundance, (C) breeding behaviors, (D) foraging behaviors, and (E) maintenance behaviors from Caminada Headland, Louisiana, USA from a boosted regression tree model. Y-axes are centered to have a zero mean over the data distribution. The relative influence (percent) of each predictor variable is shown in parentheses. Rug plots along the X-axis of each continuous variable plot illustrates the distribution of the data. Only variables with greater than 10% relative influence are shown here (refer to Supplementary Information for full model output). Abbreviated labels on the X-axis of Detailed Habitat Class include Estuarine Emergent Marsh (E.E. Marsh), Scrub/Shrub (Scrb/Shrb), Unvegetated Dune (U. Dune), Unvegetated Flat (U. Flat), and Vegetated Dune (V. Dune). Abbreviated labels on the X-axis of Inundation Class include Intertidal Unvegetated (Intertidal Unveg.) and Intertidal Vegetated (Intertidal Veg.). Abbreviated labels on the X-axis of Site include East Beach (E. Beach), West Belle Pass (W. Belle Pass), and West Beach (W. Beach). MNDWI indicates Modified Normalized Difference Water Index and MSAVI indicates Modified Soil Adjusted Vegetation Index.

### Whiskey Island

At Whiskey Island, we identified predictor variables that were important across all species in the various models. BI was found to be important for non-breeding abundance, breeding abundance, and maintenance behavior models. MNDWI was found to be important in the breeding abundance and foraging behavior models. We found MSAVI and restoration phase to be important in the non-breeding abundance and breeding abundance models, respectively. We did not identify any common important predictor variables for species where we investigated breeding behaviors.

#### American Oystercatcher

We were able to model breeding and non-breeding abundances as well as breeding, foraging, and maintenance behaviors of American Oystercatcher at Whiskey Island (Figure 7). For breeding abundance, we found meaningful associations with restoration phase (26.6%), MSAVI (15.2%), MNDWI (13.0%), and BI (11.2%; CV: 0.22 ± 0.05; Figure 7A). For the non-breeding abundance model, MNDWI (27.2%), inundation class (24.1%), MSAVI (18.2%), and BI (16.2%) were found to have the most meaningful associations with non-breeding abundance of oystercatchers (CV: 0.30 ± 0.09; Figure 7B). In the breeding behavior analysis, inundation class (80.8%) was the only variable to be considered important as all others had relative influences <5% (CV: 0.86 ± 0.02; Figure 7C). In the foraging analysis, we found foraging to be associated with BI (36.0%), MSAVI (21.5%), and MNDWI (12.8%; CV: 0.76 ± 0.02; Figure 7D). In the maintenance activities analysis, we found an association between the response variable and inundation class (67.3%) and BI (11.2%) to be the only meaningful relationships; all other variables had relative influences <7% (CV: 0.83 ± 0.02; Figure 7E). Further information on the direction of relationships is described in the Supplemental Information along with full model outputs (Figures S15–S19).

**Figure 7.**
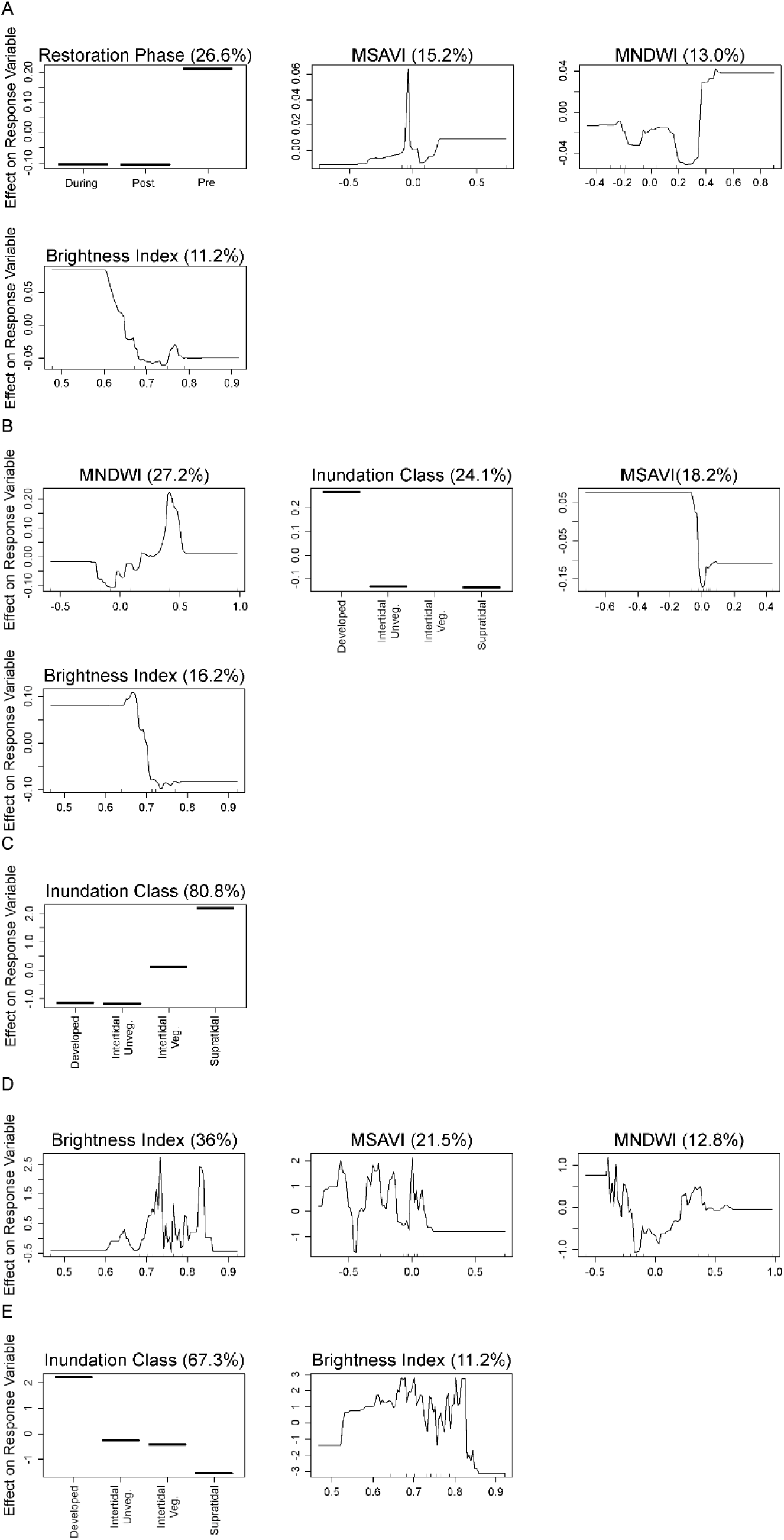
Partial dependence plots of variables related to American Oystercatcher (A) breeding abundance, (B) non-breeding abundance, (C) breeding behaviors, (D) foraging behaviors, and (E) maintenance behaviors from Whiskey Island, Louisiana, USA from a boosted regression tree model. Y-axes are centered to have a zero mean over the data distribution. The relative influence (percent) of each predictor variable is shown in parentheses. Rug plots along the X-axis of each continuous variable plot illustrates the distribution of the data. Only variables with greater than 10% relative influence are shown here (refer to Supplementary Information for full model output). Abbreviated labels on the X-axis of Detailed Habitat Class include Estuarine Emergent Marsh (E.E. Marsh), Scrub/Shrub (Scrb/Shrb), Shoreline Protection (S. Protect), Unvegetated Dune (U. Dune), and Vegetated Dune (V. Dune). Abbreviated labels on the X-axis of Inundation Class include Intertidal Unvegetated (Intertidal Unveg.) and Intertidal Vegetated (Intertidal Veg.). MNDWI indicates Modified Normalized Difference Water Index and MSAVI indicates Modified Soil Adjusted Vegetation Index.

#### Piping Plover

We were able to model non-breeding abundance as well as foraging and maintenance behaviors of Piping Plover at Whiskey Island. For non-breeding abundances, we identified meaningful relationships between abundance and BI (29.9%), MSAVI (23.1%), and distance to water (11.3%; CV: 0.25 ± 0.06; Figure 8A). For the behavioral analysis, we found foraging to be meaningfully associated with BI (26.3%), inundation class (19.9%), distance to water (13.5%), MNDWI (12.4%), and MSAVI (12.0%; CV: 0.79 ± 0.02; Figure 8B). For the maintenance behavior models, we found these activities to be affiliated with BI (23.2%), inundation class (21.8%), distance to water (14.0%), and MSAVI (12.2%; CV: 0.86 ± 0.02; Figure 8C). Further information on the direction of relationships is described in the Supplemental Information along with full model outputs (Figures S20–S22).

**Figure 8.**
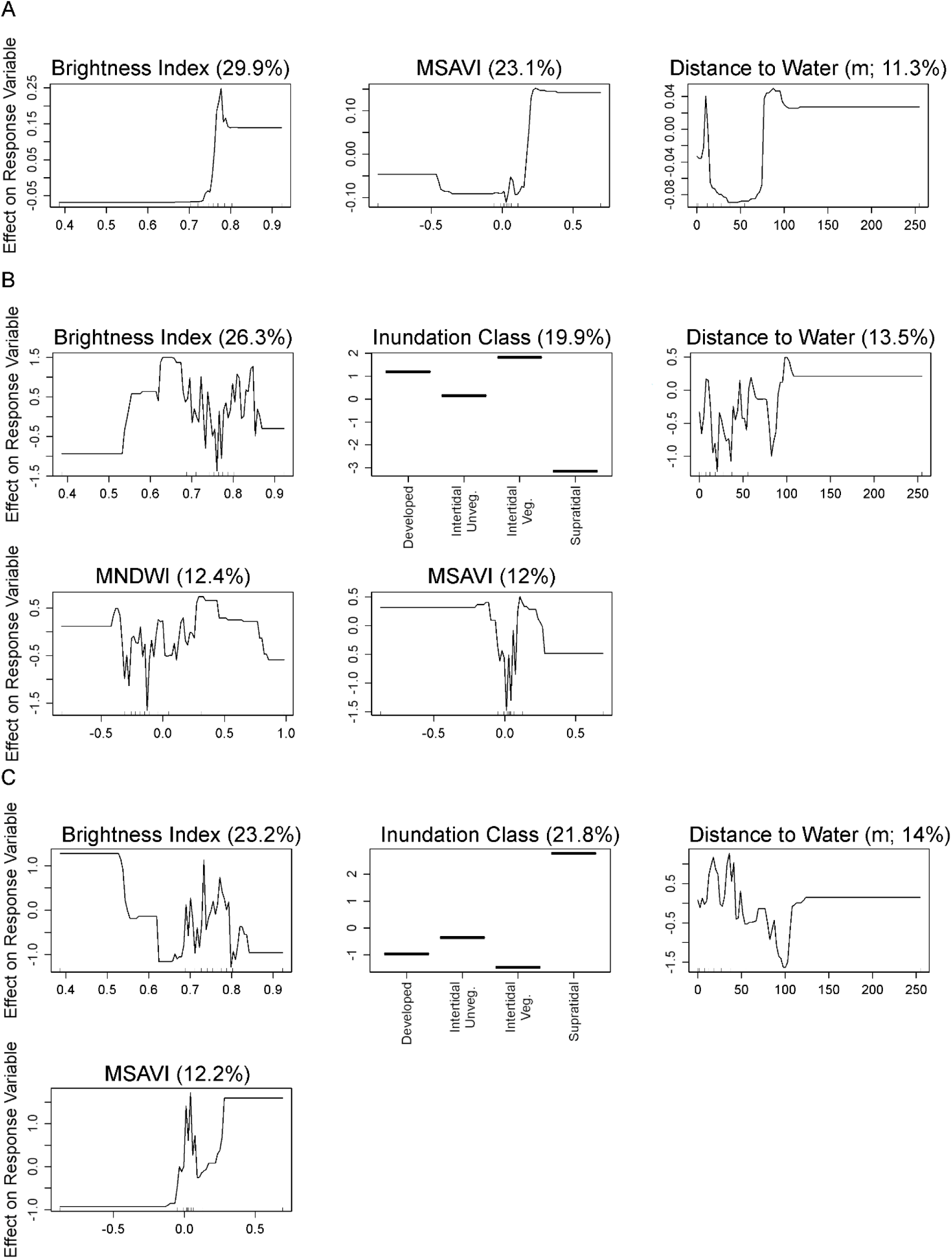
Partial dependence plots of variables related to Piping Plover (A) abundance, (B) foraging behaviors, and (C) maintenance behaviors during the non-breeding season on Whiskey Island, Louisiana, USA from a boosted regression tree model. Y-axes are centered to have a zero mean over the data distribution. The relative influence (percent) of each predictor variable is shown in parentheses. Rug plots along the X-axis of each continuous variable plot illustrates the distribution of the data. Only variables with greater than 10% relative influence are shown here (refer to Supplementary Information for full model output). Abbreviated labels on the X-axis of Detailed Habitat Class include Estuarine Emergent Marsh (E.E. Marsh), Scrub/Shrub (Scrb/Shrb), Shoreline Protection (S. Protect), Unvegetated Dune (U. Dune), and Vegetated Dune (V. Dune). Abbreviated labels on the X-axis of Inundation Class include Intertidal Unvegetated (Intertidal Unveg.) and Intertidal Vegetated (Intertidal Veg.). MNDWI indicates Modified Normalized Difference Water Index and MSAVI indicates Modified Soil Adjusted Vegetation Index.

#### Red Knot

We were able to model non-breeding abundance as well as foraging and maintenance behaviors of Red Knot at Whiskey Island. During the non-breeding season, we found meaningful relationships between abundance and distance to water (24.7%), orthoimagery-based detailed habitat classes (21.6%), BI (17.6%), and MSAVI (14.1%; CV: 0.10 ± 0.10; Figure 9A). In the behavioral analysis, we found foraging to have meaningful associations with BI (39.4%), MNDWI (20.3%), distance to water (19.0%), and MSAVI (12.6%; CV: 0.86 ± 0.01; Figure 9B). For maintenance activities, we found meaningful relationships with BI (37.2%), MSAVI (23.7%), MNDWI (21.4%), and distance to water (10.6%; CV: 0.90 ± 0.01; Figure 9C). Further information on the direction of relationships is described in the Supplemental Information along with full model outputs (Figures S23–S25).

**Figure 9.**
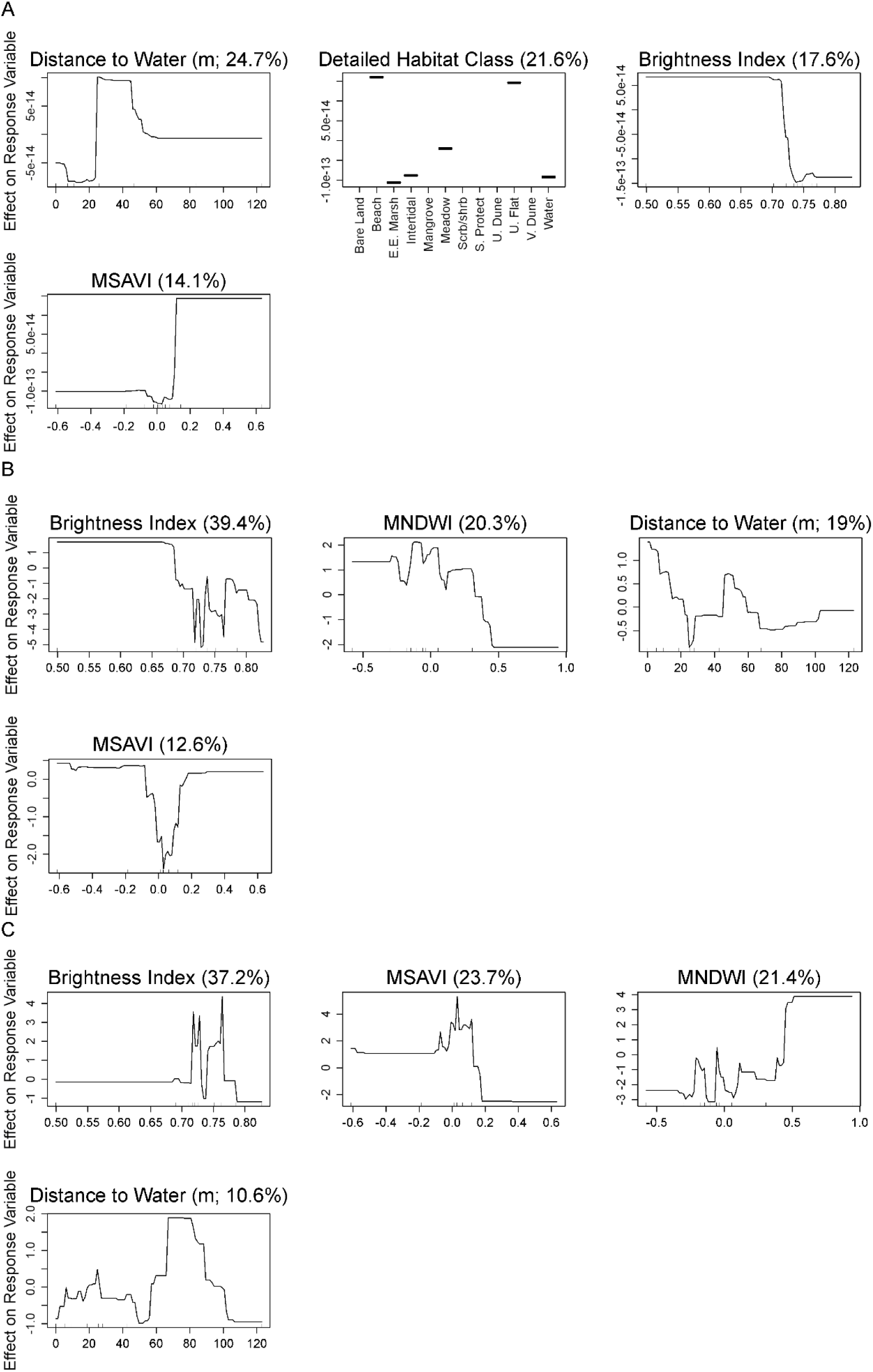
Partial dependence plots of variables related to Red Knot (A) abundance, (B) foraging behaviors, and (C) maintenance behaviors during the non-breeding seasons on Whiskey Island, Louisiana, USA from a boosted regression tree model. Y-axes are centered to have a zero mean over the data distribution. The relative influence (percent) of each predictor variable is shown in parentheses. Rug plots along the X-axis of each continuous variable plot illustrates the distribution of the data. Only variables with greater than 10% relative influence are shown here (refer to Supplementary Information for full model output). Abbreviated labels on the X-axis of Detailed Habitat Class include Estuarine Emergent Marsh (E.E. Marsh), Scrub/Shrub (Scrb/Shrb), Shoreline Protection (S. Protect), Unvegetated Dune (U. Dune), and Vegetated Dune (V. Dune). Abbreviated labels on the X-axis of Inundation Class include Intertidal Unvegetated (Intertidal Unveg.) and Intertidal Vegetated (Intertidal Veg.). MNDWI indicates Modified Normalized Difference Water Index and MSAVI indicates Modified Soil Adjusted Vegetation Index.

#### Snowy Plover

We were able to model non-breeding abundance as well as foraging and maintenance behaviors of Snowy Plover at Whiskey Island. For non-breeding abundances, we found meaningful relationships between number of individuals and BI (40.6%), MSAVI (12.2%), and MNDWI (11.1%; CV: 0.24 ± 0.06; Figure 10A). In our behavioral analysis, we found foraging behaviors to be associated with MSAVI (35.5%), BI (18.6%), MNDWI (12.4%), and inundation class (11.5%; CV: 0.78 ± 0.02; Figure 10B). Snowy Plover maintenance behaviors were associated with MSAVI (34.0%), BI (23.2%), and inundation class (14.1%; CV: 0.84 ± 0.02; Figure 10C). Further information on the direction of relationships is described in the Supplemental Information along with full model outputs (Figures S26–S28).

**Figure 10.**
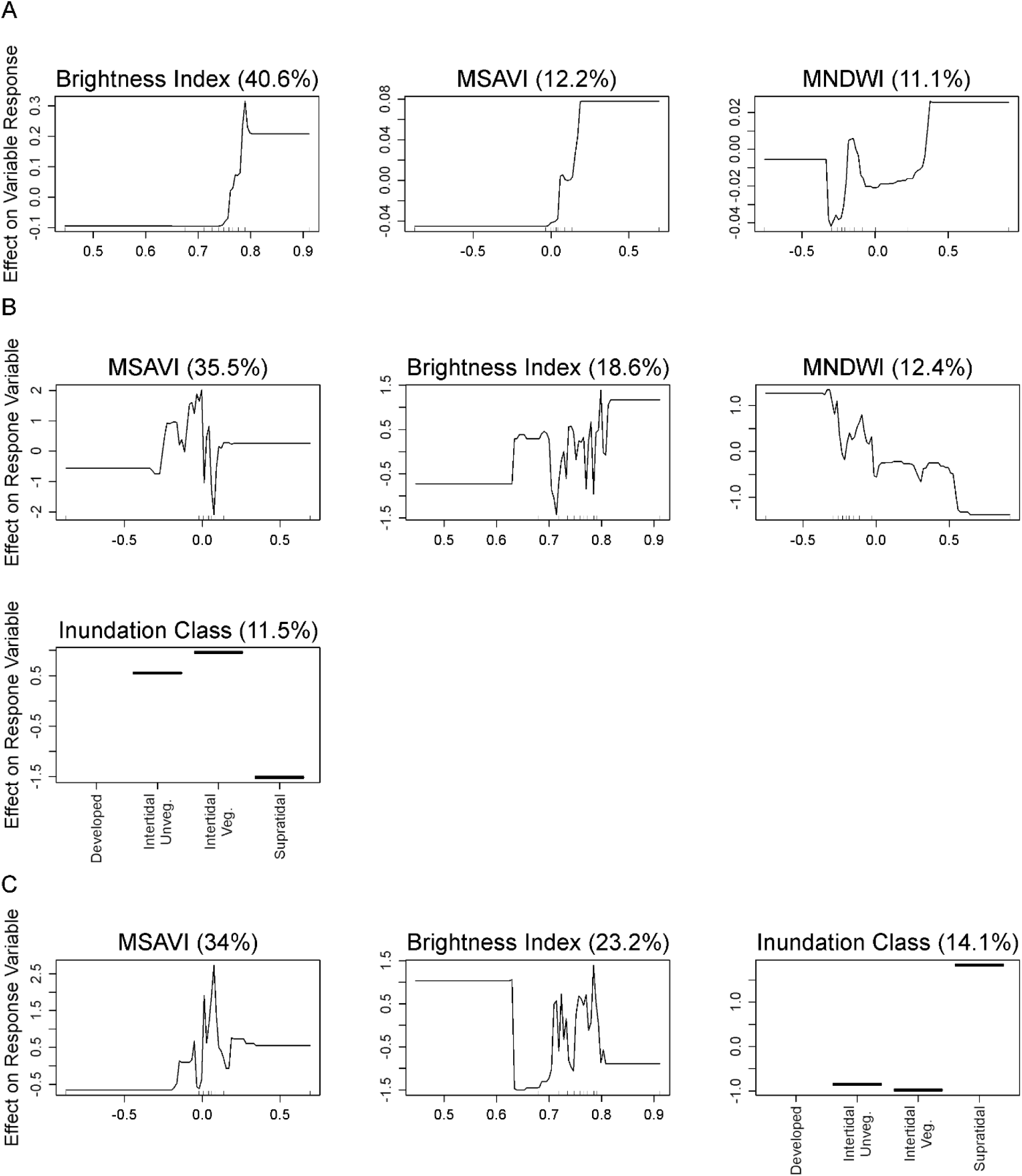
Partial dependence plots of variables related to Snowy Plover (A) abundance, (B) foraging behaviors, and (C) maintenance behaviors during the non-breeding seasons on Whiskey Island, Louisiana, USA from a boosted regression tree model. Y-axes are centered to have a zero mean over the data distribution. The relative influence (percent) of each predictor variable is shown in parentheses. Rug plots along the X-axis of each continuous variable plot illustrates the distribution of the data. Only variables with greater than 10% relative influence are shown here (refer to Supplementary Information for full model output). Abbreviated labels on the X-axis of Detailed Habitat Class include Estuarine Emergent Marsh (E.E. Marsh), Scrub/Shrub (Scrb/Shrb), Shoreline Protection (S. Protect), Unvegetated Dune (U. Dune), and Vegetated Dune (V. Dune). Abbreviated labels on the X-axis of Inundation Class include Intertidal Unvegetated (Intertidal Unveg.) and Intertidal Vegetated (Intertidal Veg.). MNDWI indicates Modified Normalized Difference Water Index and MSAVI indicates Modified Soil Adjusted Vegetation Index.

#### Wilson’s Plover

We were able to model breeding and non-breeding abundances as well as breeding, foraging, and maintenance behaviors of Wilson’s Plover at Whiskey Island. During the breeding season, we found important associations between breeding abundance of Wilson’s Plover and BI (22.7%), tidal stage (17.8%), restoration phase (12.9%), MNDWI (12.1%), and orthoimagery-based detailed habitat maps (10.3%; CV: 0.20 ± 0.01; Figure 11A). During the non-breeding season, we found meaningful associations between abundance and tidal stage (26.1%), MSAVI (23.2%), orthoimagery-based detailed habitat classes (15.1%), and BI (10.4%; CV: 0.09 ± 0.05; Figure 11B). In our behavioral analysis, we found MSAVI (22.7%), MNDWI (22.0%), BI (18.2%), and distance to water (13.0%; CV: 0.85 ± 0.00) to be strongly associated with breeding behaviors (Figure 11C). Foraging behaviors were meaningfully related to two variables, satellite-based habitat classes (61.0%) and MNDWI (12%; CV: 0.72 ± 0.01; Figure 11D). When analyzing maintenance activities, we found MSAVI (31.0%), MNDWI (17.5%), BI (17.4%), and distance to water (11.4%) to be the most important variables (CV: 0.90 ± 0.00; Figure 11E). Further information on the direction of relationships is described in the Supplemental Information along with full model outputs (Figures S29–S33).

**Figure 11.**
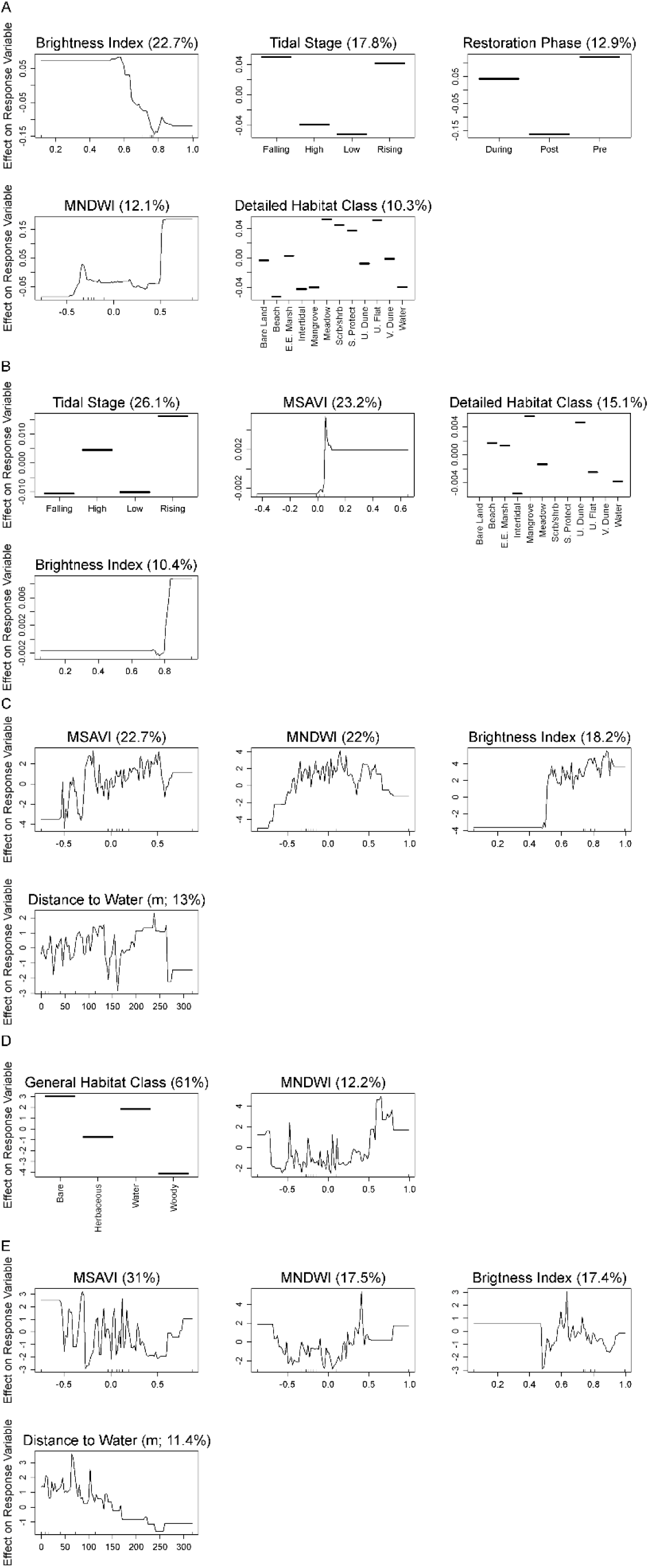
Partial dependence plots of variables related to Wilson’s Plover (A) breeding abundance, (B) non-breeding abundance, (C) breeding behaviors, (D) foraging behaviors, and (E) maintenance behaviors during the non-breeding seasons on Whiskey Island, Louisiana, USA from a boosted regression tree model. Y-axes are centered to have a zero mean over the data distribution. The relative influence (percent) of each predictor variable is shown in parentheses. Rug plots along the X-axis of each continuous variable plot illustrates the distribution of the data. Abbreviated labels on the X-axis of Detailed Habitat Class include Estuarine Emergent Marsh (E.E. Marsh), Scrub/Shrub (Scrb/Shrb), Shoreline Protection (S. Protect), Unvegetated Dune (U. Dune), and Vegetated Dune (V. Dune). Abbreviated labels on the X-axis of Inundation Class include Intertidal Unvegetated (Intertidal Unveg.) and Intertidal Vegetated (Intertidal Veg.). MNDWI indicates Modified Normalized Difference Water Index and MSAVI indicates Modified Soil Adjusted Vegetation Index.

## Discussion

Improving our understanding of how shorebirds use coastal habitats can inform restoration and management of barrier islands and coastal headlands to benefit wildlife. In our study, we found habitat factors, such as greenness, brightness, soil wetness, and distance to water, to be the most important variables in explaining abundances as well as behaviors of our focal species. This result is likely due to the complex habitat characteristics required by shorebirds to meet their functional needs (i.e., breeding, foraging, maintenance). We only found restoration phase to be considered meaningful in breeding species at Whiskey Island, which may suggest birds may be more flexible during the non-breeding season when using coastal habitats. The decrease in abundance of breeding Wilson’s Plover and American Oystercatcher during active and post restoration periods on Whiskey Island is likely due to the changes in habitat as well as the breeding community we observed following restoration (e.g., increases in number of colonial breeding birds). Given pressures shorebirds face (e.g., human disturbance, predation, and habitat loss/degradation) and documented population declines (American Oystercatcher Working Group et al., 2020; Baker et al., 2020; Elliott-Smith & Haig, 2020; Page et al., 2020; Rosenberg et al., 2019; Smith et al., 2023; Zdravkovic et al., 2023), our results show that coastal restoration projects continue to provide suitable habitat for shorebirds, particularly during the non-breeding season. Our findings also identify habitat features that could be considered when designing and managing coastal habitat projects.

### Caminada Headland

Despite the restoration at Caminada Headland being substantial, with ∼8.8 million m^3^ of offshore sediment used to create or enhance ∼429 ha of beach and dune habitat (Coastal Protection and Restoration Authority of Louisiana, 2018), we did not find restoration phase to be important in any of the models. Compared to Whiskey Island, there was more sediment deposited and more habitat created or enhanced at Caminada Headland (Coastal Protection and Restoration Authority of Louisiana, 2018). However, because the restoration occurred over a much larger area and over a longer period, the degree of change appears less dramatic. Despite substantial alteration to the existing habitat, we found no direct influence of restoration phase on our focal species. This result contradicts the few studies evaluating coastal restoration along the northern Gulf coast, which have generally found that birds avoid using areas following restoration (Arfman, 2016; Convertino et al., 2011). These studies documented declines in shorebird use within a year of the restoration occurring, which could be due to a decrease in shorebird prey items (i.e., benthic invertebrates) from being buried by new beach fill (Peterson et al., 2006, 2014). The post-restoration phase spanned ∼3 years, which is longer than previous studies and may have allowed for the habitat to resemble a non-altered state. For example, six months after sand nourishment, avian richness in parts of Holly Beach, Louisiana was generally comparable to that of nearby non-nourished areas (Arfman, 2016). Whereas current results suggest that abundance and frequency of behaviors may not be influenced by coastal restoration at longer time scales, there could still be beneficial or detrimental responses (e.g., changes in productivity, energetic condition, community diversity) outside the scope of our investigation.

When looking at meaningful predictor variables for Caminada Headland, we found MNDWI, BI, and distance to water to be important in all but one model and MSAVI in all but two models. In the majority of models, MNDWI, BI, and distance to water showed generally positive trends with the response variable. In terms of MNDWI, it may not be surprising that birds were observed to be more abundant and engage in more behaviors as the amount of wetness increased given that our focal species tend to use intertidal habitats for foraging and other activities (American Oystercatcher Working Group et al., 2020; Baker et al., 2020; Elliott-Smith & Haig, 2020; Zdravkovic et al., 2023). The results for MNDWI appear to match our distance to water metric, which found most individuals within 100 m of the water’s edge. Most of the observations occurred within a narrow band of BI values, which typically spanned ∼0.7–0.8 and appear to include mostly bare land or sparse herbaceous vegetation. For MSAVI, we found most trends tended to be negative, suggesting more birds were using habitats with less vegetative cover. However, due to differences we observed between satellite sensors for sampling MSAVI, we suggest future efforts utilize a single sensor or harmonized data (e.g., Claverie et al., 2018), when available.

### Whiskey Island

At Whiskey Island, the restoration caused a substantial amount of terrestrial habitat to be created, especially on both the Gulf and bay sides of the western portion of the island as well as the Gulf side of the eastern portion of the island. While the restoration did not seem to have any meaningful impact during the non-breeding seasons, we found abundances of American Oystercatcher and Wilson’s Plover to decrease during the breeding season following restoration. Changes to microhabitat potentially contributed to the decline in Wilson’s Plover numbers following restoration.

Wilson’s Plover tend to select nesting sites with sparse to moderate vegetation (Bergstrom, 1988; DeRose-Wilson et al., 2013; Zdravkovic et al., 2023) and much of the restoration created unvegetated habitats that may be less suitable. Additionally, chicks typically relocate to wet and/or densely vegetated areas prior to fledgling (Bergstrom, 1988; DeRose-Wilson et al., 2013; Dikun, 2008; Ray, 2011), and the restoration may have caused a greater distance between Wilson’s Plover nest sites and areas with cover and food. Change in the avian breeding community, which included an increase in colonial nesting birds like Black Skimmer (*Rynchops niger*), Gull-billed Tern (*Gelochelidon nilotica*), and Least Tern (*Sternula antillarum*), also likely impacted both American Oystercatcher and Wilson’s Plover. While both of our breeding focal species are known to nest near or even among colonial nesting birds, they are also susceptible to displacement from other nesting species (American Oystercatcher Working Group et al., 2020; Zdravkovic et al., 2023). Future studies may consider investigating the interplay between coastal restoration and interspecific interactions to understand how restoration actions could impact community ecology.

Overall, for Whiskey Island, we found BI to be considered important in almost all analyses, which was closely followed by MSAVI. In the majority of cases, BI seemed to be positively associated with abundance or behavior. Interestingly, most of the observations occurred within a narrow band of BI values, which typically spanned ∼0.7–0.8, that appear to include mostly bare land or sparse herbaceous vegetation. We also found MSAVI to have a generally positive association with abundance or behavior, but in most analyses, observations were centered around zero (bare land). We found most individuals to be in non-vegetated habitats or areas that are transitioning from bare land to sparse vegetation on the landscape.

## Conclusion

Most of our analyses found that birds did not respond to the overall effect of our “restoration phase” variable in Louisiana, but breeding abundances of American Oystercatcher and Wilson’s Plover were higher at Whiskey Island prior to restoration. However, changes in habitat (refer to Thurman et al., 2023) paired with our results of birds increasing use of newly created habitat types suggest that Piping Plover, Red Knot, Snowy Plover, and Wilson’s Plover at Caminada Headland as well as Red Knot and Wilson’s Plover (foraging and non-breeding abundance) at Whiskey Island tended to benefit from the restoration. These results suggest an overall net benefit with some relationships (i.e., Wilson’s Plover) being more complex in response to restoration activities. We also found remotely sensed variables from publicly accessible satellite imagery, which likely changed due to the restoration activities, tended to have the greatest importance across most models. This result suggests that these remote sensing metrics may serve as a tool for designing as well as evaluating restoration and construction projects in addition to predicting avian habitat use. Continuing to integrate remotely sensed data with ground-based avian data collection, as we have done, can increase knowledge on the complex life histories of wildlife species and, subsequently, may elicit changes in construction design to maximize ecosystem benefits (*sensu* Windhoffer et al., 2024).

## Acknowledgements

We thank D. Lee, P. Leberg, J. Sylvest, E. Koen, and M.J. Osland for feedback on earlier versions of this manuscript as well as K. Thompson, H. Cortes, and B. Landry for assistance with figures. This work was supported by the CPRA (contract number: 2000387259), the Barataria-Terrebonne National Estuary Program, and the U. S. Geological Survey. Any use of trade, firm, or product names is for descriptive purposes only and does not imply endorsement by the U. S. Government.

## Literature Cited

American Oystercatcher Working Group, Nol, E., & Humphrey, R. C. (2020). American Oystercatcher (Haematopus palliatus). In A. F. Poole (Ed.), Birds of the World (1.0). Cornell Lab of Ornithology.

Arfman, A. R. (2016). Evaluating the effects of coastal restoration on shorebirds and shorebird habitats in Cameron Parish, Louisiana [Master’s Thesis]. McNeese State University.

Atkinson, P. W. (2003). Can we recreate or restore intertidal habitats for shorebirds? Bulletin-Wader Study Group, 100, 67–72.

Baker, A., Gonzalez, P., Morrison, R. I. G., & Harrington, B. A. (2020). Red Knot (Calidris canutus). In S. M. Billerman (Ed.), Birds of the World: Vol. Version 1.0. Cornell Lab of Ornithology. 10.2173/bow.redkno.01

Bergstrom, P. W. (1988). Breeding biology of Wilson’s Plovers. Wilson Bulletin, 100, 25–35.

Bivand, R. S., Pebesma, E. J., & Gómez-Rubio, V. (2013). Applied spatial data analysis with R (2nd ed.). Springer.

Brusati, E. D., DuBowy, P. J., & Lacher, T. E. (2001). Comparing ecological functions of natural and created wetlands for shorebirds in Texas. Waterbirds: The International Journal of Waterbird Biology, 24(3), 371–380. 10.2307/1522067

Brush, J. M., Pruner, R. A., & Driscoll, M. J. L. (2019). Shorebirds. In R. R. Wilson, A. M. V. Fournier, J. S. Gleason, J. E. Lyons, & M. S. Woodrey (Eds.), GoMAMN Strategic Bird Monitoring Guidelines (pp. 171–202). Mississippi State University.

Byrnes, M. R., Berlinghoff, J. L., Griffee, S. F., & Lee, D. M. (2018). Louisiana Barrier Island Comprehensive Monitoring Program (BICM): Phase 2–Updated Shoreline Compilation and Change Assessment, 1880s to 2015 (p. 46). Prepared for Louisiana Coastal Protection and Restoration Authority (CPRA) by Applied Coastal Research and Engineering.

Claverie, M., Ju, J., Masek, J. G., Dungan, J. L., Vermote, E. F., Roger, J.-C., Skakun, S. V., & Justice, C. (2018). The Harmonized Landsat and Sentinel-2 surface reflectance data set. Remote Sensing of Environment, 219, 145–161. 10.1016/j.rse.2018.09.002

Coastal Protection and Restoration Authority of Louisiana. (2018). Barrier Island Status Report: Fiscal Year 2020 Annual Plan (pp. 1–30). Coastal Protection and Restoration Authority of Louisiana.

Convertino, M., Donoghue, J., Chu-Agor, M. L., Kiker, G., Munoz-Carpena, R., Fischer, R., & Linkov, I. (2011). Anthropogenic renourishment feedback on shorebirds: A multispecies bayesian perspective. Nature Precedings. 10.1038/npre.2011.5872.1

DeRose-Wilson, A., Fraser, J. D., Karpanty, S. M., & Catlin, D. H. (2013). Nest-site selection and demography of Wilson’s Plovers on a North Carolina barrier island. Journal of Field Ornithology, 84(4), 329–344. 10.1111/jofo.12033

Dikun, K. A. (2008). Nest-site selection of Wilson’s Plovers (Charadrius wilsonia) in South Carolina [Master’s Thesis]. Coastal Carolina University.

Dowle, M., & Srinivasan, A. (2021). Data.table: Extension of ‘data.fram’. R package version 1.14.0. https://CRAN.R-project.org/package=data.table

Drake, K. R., Thompson, J. E., Drake, K. L., & Zonick, C. (2001). Movements, Habitat Use, and Survival of Nonbreeding Piping Plovers. The Condor, 103(2), 259–267. 10.1093/condor/103.2.259

Elith, J., Leathwick, J. R., & Hastie, T. (2008). A working guide to boosted regression trees. Journal of Animal Ecology, 77(4), 802–813. 10.1111/j.1365-2656.2008.01390.x

Elliott-Smith, E., & Haig, S. M. (2020). Piping Plover (Charadrius melodus). In A. F. Poole (Ed.), Birds of the World: Vol. Version 1.0. Cornell Lab of Ornithology. 10.2173/bow.pipplo.01

Enwright, N. M., SooHoo, W. M., Dugas, J. L., Conzelmann, C. P., Laurenzano, C., Lee, D. M., Mouton, K., & Stelly, S. J. (2020). Louisiana Barrier Island Comprehensive Monitoring Program: Mapping habitats in beach, dune, and intertidal environments along the Louisiana Gulf of Mexico shoreline, 2008 and 2015–16 (U.S. Geological Survey Open-File Report Nos. 2020–1030; p. 57). U.S. Geological Survey. http://pubs.er.usgs.gov/publication/ofr20201030

ESA. (2015). Sentinel-2 User Handbook, Issue 1, *Revision 2*. European Space Agency. https://sentinel.esa.int/documents/247904/685211/Sentinel-2_User_Handbook

Escadafal, R., Girard, M.-C., & Courault, D. (1989). Munsell soil color and soil reflectance in the visible spectral bands of landsat MSS and TM data. Remote Sensing of Environment, 27(1), 37–46. 10.1016/0034-4257(89)90035-7

Farrell, C., Korosy, M., Wraithmell, J., & Audubon Florida. (2016). The role of coastal engineering in American Oystercatcher conservation. Audubon Florida.

Folse, T., & Lee, D. (2016). Monitoring plan for Caminada Headland beach and dune restroation incr. 2 (BA-0143). Coastal Protection and Restoration Authority of Louisiana.

Gibson, D., Chaplin, M. K., Hunt, K. L., Friedrich, M. J., Weithman, C. E., Addison, L. M., Cavalieri, V., Coleman, S., Cuthbert, F. J., Fraser, J. D., Golder, W., Hoffman, D., Karpanty, S. M., Van Zoeren, A., & Catlin, D. H. (2018). Impacts of anthropogenic disturbance on body condition, survival, and site fidelity of nonbreeding Piping Plovers. The Condor, 120(3), 566–580. 10.1650/CONDOR-17-148.1

Greenwell, B., Boehmke, B., Cunningham, J., & GBM Developers. (2020). Gbm: Generalized Boosted Regression Models. R package version 2.1.8. https://CRAN.R-project.org/package=gbm

Hijmans, R. J. (2022). raster: Geographic data analysis and modeling. R package version 3.5-21. https://CRAN.R-project.org/package=raster

Hijmans, R. J., Phillips, S., Leathwick, J., & Elith, J. (2021). dismo: Species Distribution Modeling. R package version 1.3-5. The R Foundation for Statistical Computing, Vienna Http://Cran.r-Project. Org. https://CRAN.R-project.org/package=dismo

Iglecia, M., & Winn, B. (2021). A shorebird management manual. Manomet.

Lauro, B., & Burger, J. (1989). Nest-site selection of American Oystercatchers (*Haematopus palliatus*) in salt marshes. The Auk, 106(2), 185–192.

LeBlanc, D., Anderson, A. N., Leberg, P. L., Waddle, J. H., Enwright, N. M., Thurman, H. R., & Zenzal Jr, T. J. (2023). An introduction to the evaluation of restoration for avian species at Caminada Headland and Whiskey Island in Louisiana (Evaluation of Restoration for Avian Species at Caminada Headland and Whiskey Island in Louisiana). Coastal Protection and Restoration Authority of Louisiana.

LeDee, O. E., Cuthbert, F. J., & Bolstad, P. V. (2008). A remote sensing analysis of coastal habitat composition for a threatened shorebird, the Piping Plover (Charadrius melodus). Journal of Coastal Research, *24*(3 (243)), 719–726. 10.2112/06-0734.1

Mangiafico, S. S. (2025). Rcompanion: Functions to Support Extension Education Program Evaluation. Version 2.5.0. Rutgers Cooperative Extension. https://CRAN.R-project.org/package=rcompanion

McIntyre, A. F., & Heath, J. A. (2011). Evaluating the effects of foraging habitat restoration on shorebird reproduction: The importance of performance criteria and comparative design. Journal of Coastal Conservation, 15(1), 151–157. 10.1007/s11852-010-0128-x

Müller, D., Leitão, P. J., & Sikor, T. (2013). Comparing the determinants of cropland abandonment in Albania and Romania using boosted regression trees. Agricultural Systems, 117, 66–77.

Noel, B. L., & Chandler, C. R. (2008). Spatial distribution and site fidelity of non-breeding Piping Plovers on the Georgia coast. Waterbirds, 31(2), 241–251. 10.1675/1524-4695(2008)31%255B241:SDASFO%255D2.0.CO;2

Page, G. W., Stenzel, L. E., Warriner, J. S., Warriner, J. C., & Paton, P. W. (2020). Snowy Plover (Charadrius nivosus). In A. F. Poole (Ed.), Birds of the World: Vol. Version 1.0. Cornell Lab of Ornithology. 10.2173/bow.snoplo5.01

Pebesma, E., & Bivand, R. S. (2005). Classes and methods for spatial data in R. R News, 5(2), 9–13.

Peterson, C. H., Bishop, M. J., D’Anna, L. M., & Johnson, G. A. (2014). Multi-year persistence of beach habitat degradation from nourishment using coarse shelly sediments. Science of The Total Environment, 487, 481–492. 10.1016/j.scitotenv.2014.04.046

Peterson, C. H., Bishop, M. J., Johnson, G. A., D’Anna, L. M., & Manning, L. M. (2006). Exploiting beach filling as an unaffordable experiment: Benthic intertidal impacts propagating upwards to shorebirds. Journal of Experimental Marine Biology and Ecology, 338(2), 205–221. 10.1016/j.jembe.2006.06.021

Qi, J., Chehbouni, A., Huete, A. R., Kerr, Y. H., & Sorooshian, S. (1994). A modified soil adjusted vegetation index. Remote Sensing of Environment, 48(2), 119–126. 10.1016/0034-4257(94)90134-1

R Core Team. (2021). R: A language and environment for statistical computing (Version 4.1.0) [Computer software]. R Foundation for Statistical Computing. https://www.R-project.org

Ray, K. L. (2011). Factor’s affecting Wilson’s Plover (Charadrius wilsonia) demography and habitat use at Onslow Beach, Marine Corps Base Camp Lejune, North Carolina [Master’s Thesis]. Virginia Polytechnic Institute and State University.

Remsen, J. V., Jr., Wallace, B. P., Seymour, M. A., O’Malley, D. A., & Johnson, E. I. (2019). The regional, national, and international importance of Louisiana’s coastal avifauna. The Wilson Journal of Ornithology, 131(2), 221–434. 10.1676/18-111

Rosenberg, K. V., Dokter, A. M., Blancher, P. J., Sauer, J. R., Smith, A. C., Smith, P. A., Stanton, J. C., Panjabi, A., Helft, L., Parr, M., & Marra, P. P. (2019). Decline of the North American avifauna. Science, 366(6461), 120–124. 10.1126/science.aaw1313

Schofield, L. N., Deppe, J. L., Zenzal, T. J., Ward, M. P., Diehl, R. H., Bolus, R. T., & Moore, F. R. (2018). Using automated radio telemetry to quantify activity patterns of songbirds during stopover. The Auk, 135(4), 949–963. 10.1642/AUK-17-229.1

Schulte, S. A., & Simons, T. R. (2014). Factors affecting the reproductive success of American Oystercatchers *Haematopus palliatus* on the outer banks of North Carolina. Marine Ornithology, 43, 37–47.

Schulz, J. L., & Leberg, P. L. (2019). Factors Affecting Prey Availability and Habitat Use of Nonbreeding Piping Plovers (Charadrius melodus) in Coastal Louisiana. Journal of Coastal Research, 35(4), 861–871. 10.2112/JCOASTRES-D-17-00147.1

Smith, P. A., Smith, A. C., Andres, B., Francis, C. M., Harrington, B., Friis, C., Morrison, R. I. G., Paquet, J., Winn, B., & Brown, S. (2023). Accelerating declines of North America’s shorebirds signal the need for urgent conservation action. Ornithological Applications, 125(2), duad003. 10.1093/ornithapp/duad003

Thurman, H. R., Enwright, N. M., Cheney, W. C., Dugas, J. L., Lee, D. M., & Jones, W. R. (2023). Mapping habitats and shorelines pre-, during, and post-restoration on Caminada Headland and Whiskey Island, Louisiana, 2012–2020 (p. 51p.) [Final Report]. Louisiana Coastal Protection and Restoration Authority.

U.S. Fish and Wildlife Service. (2015). Recovery Plan for the Northern Great Plains piping plover (Charadrius melodus) in two volumes. Volume I: Draft breeding recovery plan for the Northern Great Plains piping plover (Charadrius melodus) and Volume II: Draft revised recovery plan for the wintering range of the Northern Great Plains piping plover (Charadrius melodus) and Comprehensive conservation strategy for the piping plover (Charadrius melodus) in its coastal migration and wintering range in the continental United States (p. 166).

USGS. (2021). Landsat 8 Collection 2 (C2) Level 2 Science Product (L2SP) Guide v. 3.0 (No. USGS LSDS-1619; p. 37). U.S. Geological Survey, Earth Resources Observation and Science (EROS) Center. https://d9-wret.s3.us-west-2.amazonaws.com/assets/palladium/production/s3fs-public/media/files/LSDS-1619_Landsat8-C2-L2-ScienceProductGuide-v3.pdf

Wei, T., & Simko, V. (2021). R package “corrplot”: Visualization of a Correlation Matrix (Version 0.90). https://github.com/taiyun/corrplot

Wickham, H., François, R., Henry, L., & Müller, K. (2022). Dplyr: A grammar of data manipulation. R package version 1.0.9. https://CRAN.R-project.org/package=dplyr

Windhoffer, E. D., Carruthers, T. J. B., Henkel, J., Gleason, J. S., & Wiebe, J. J. (2024). Leveraging co-production within ecosystem restoration to maximize benefits to coastal birds. Journal of Environmental Management, 360, 121093. 10.1016/j.jenvman.2024.121093

Withers, K. (2002). Shorebird Use of Coastal Wetland and Barrier Island Habitat in the Gulf of Mexico. The Scientific World Journal, 2, 514–536. 10.1100/tsw.2002.112

Xu, H. (2006). Modification of normalised difference water index (NDWI) to enhance open water features in remotely sensed imagery. International Journal of Remote Sensing, 27(14), 3025–3033. 10.1080/01431160600589179

Zdravkovic, M. G., Corbat, C. A., & Bergstrom, P. W. (2023). Wilson’s Plover (*Charadrius wilsonia*). In P. G. Rodewald (Ed.), Birds of the World (1.1). Cornell Lab of Ornithology. 10.2173/bow.wilplo.01.1

Zenzal Jr., T. J., Anderson, A. N., Geary, B. J., Schulz, J. L., Dobbs, R. C., Barrow Jr, W. C., & Waddle, J. H. (2025). Early season tropical cyclones affect birds breeding on a barrier island. Gulf and Caribbean Research 36: 38–48. 10.18785/gcr.3601.08

Zenzal Jr., T. J., Anderson, A. N., & LeBlanc, D. (2023). Evaluating if abundance and behavior of shorebird species are related to restoration and habitat at Whiskey Island and Caminada Headland, Louisiana from 2012 to 2020 [Dataset]. ScienceBase. 10.5066/P93MVS0S

