## Supplemental Information for "Understanding Abundances and Behaviors of Shorebirds in Coastal Louisiana"

*This information product has been peer reviewed and approved for publication as a preprint by the U.S. Geological Survey.*

**Supplemental Information**

### Understanding Abundances and Behaviors of Shorebirds in Coastal Louisiana

T.J. Zenzal Jr. [1]*, Amanda N. Anderson[1], Nicholas M. Enwright[1], Hana R. Thurman[2], Wyatt C. Cheney[3], Delaina LeBlanc[4], Robert C. Dobbs[1+], Brock Geary[1†], and J. Hardin Waddle[5]

[1]U.S. Geological Survey (USGS), Wetland and Aquatic Research Center (WARC), Lafayette, LA

[2]Cherokee Nation System Solutions, Contractor to the USGS WARC, Lafayette, LA

[3]Lafayette, LA, Contractor to the USGS WARC,

[4]Barataria-Terrebonne National Estuary Program, Thibodaux, LA

[5]USGS, WARC, Gainesville, FL

[+] Present address: Louisiana Department of Wildlife and Fisheries, Lafayette, LA

[†]Present address: University of Pennsylvania School of Veterinary Medicine, Kennett Square, PA

#### Methods

##### Geospatial Analyses

Orthoimagery-based habitat maps – The methods for orthoimagery-based habitat map development are found in Thurman et al. (2023) and some details are repeated here for clarity. We used orthoimagery to produce annual, detailed habitat classifications through a semi-automated approach. Habitat maps were created for the years 2012, 2015, and 2017–2019, spanning the time periods before, during, and after restoration. The two sources of orthoimagery for this mapping effort were the National Agriculture Imagery Program (NAIP) statewide imagery for Louisiana, and the Coastwide Reference Monitoring System (CRMS) digital orthophoto quarter quadrangles (DOQQs) for coastal Louisiana (Tables S3 and S4). Aerial orthoimagery from both sources was four-band with a spatial resolution of 1 m, except for the 2019 NAIP imagery, which had a resolution of 0.6 m. The 2019 NAIP imagery was resampled to 1 m using bilinear interpolation for consistency with other maps in this series. Descriptions of the habitat classes mapped are found in Table S1.

Map-based inundation classes and distance to water – The orthoimagery-based habitat maps were converted to Barrier Island Comprehensive Monitoring Program (BICM) inundation zone-based classes (Enwright et al., 2020). Table S2 shows the crosswalk between the detailed classes and the inundation zone-based classes. For each of the orthoimagery-based habitat maps, we developed a Euclidean distance raster to indicate the distance to the nearest water pixel to determine the minimal distance a bird may have been from a water source.

Satellite-based habitat maps – The methods for satellite-based habitat map development are found in Thurman et al. (2023) and some details are repeated here for the convenience of the reader. We used satellite imagery to produce interannual maps that were at or near the time of the avian surveys and covered the time period before, during, and after restoration. Cloud-free satellite imagery was inventoried for Whiskey Island from 2013–2020 and Caminada Headland from 2013–2019 during the time windows of the avian surveys (Table S5 and S6). Both Landsat 8 and Sentinel-2 imagery were used for this assessment. Landsat 8 was launched in mid-February 2013 and Sentinel-2 was launched in late June 2015.

These satellite data were pan-sharpened (refer to Thurman et al. [2023] for details). We calculated several spectral indices from each satellite image (Thurman et al., 2023). The spectral indices that were found to be important included greenness indices, wetness indices, and a brightness index (Table S7). The classes for the satellite-based habitat maps included: 1) bare, 2) herbaceous vegetation, 3) woody vegetation, and 4) water. More detailed information on the satellite-based habitat maps including the accuracy of the maps can be found in Thurman et al. (2023).

Extraction to avian surveys – Esri point shapefiles were created for the Whiskey Island and the Caminada Headland avian survey databases. We extracted geospatial predictor variables to each bird location. Specifically, these included: 1) detailed habitat classes from the orthoimagery-based maps, 2) map-based inundation classes, 3) distance to water, 4) satellite-based habitat classes, and 5) a subset of the satellite indices focused on greenness, wetness, and brightness. For Whiskey Island, we extracted the predictor values at the exact location of the bird observations. Since the Caminada bird locations include some ambiguity (refer to main text), we used focal statistics with a 50-m circular kernel to determine the majority non-water habitat class and mean value for each spectral index. The date of the avian survey was used to determine what geospatial predictor was most relevant. Bird observations that overlapped with areas that were included in the cloud and cloud shadow mask were set to “no data” for satellite-based predictors (i.e., satellite-based habitat classes and spectral indices). To assist with the interpretation of the spectral indices, we have developed violin plots using points from the satellite-based maps that were assessed for accuracy (Table S5 and S6).

Inundation Assignments – We assigned a field or map-based inundation class to each bird observation (Table S2; Enwright et al., 2020). For Whiskey Island, we cross-walked the habitat type assigned during field surveys to fit within a map-based inundation class for each bird observation instead of using the map-based inundation class assigned from remote sensing. We categorized the habitat types into the following inundation classes: 1) intertidal-unvegetated (e.g., foreshore beach, tidal flat, washover flat), 2) intertidal-vegetated (e.g., estuarine emergent marsh, algal flat), 3) supratidal (e.g., backshore beach, sand flat, vegetated dune, shell), and 4) developed/shoreline protection (e.g., jetty and platform spoil; Enwright et al., 2020; Table S6). Additionally, we assigned a tidal stage for each Whiskey Island survey to be high or low if the midpoint of a survey fell within 1.5 hours of predicted high or low tide per the National Oceanic and Atmospheric Administration Caillou Boca tide station (Station ID: 8763206); otherwise rising or falling was used as appropriate (Schulz & Leberg, 2019). For Caminada Headland, we assigned a map-based inundation class based on the habitat classification schemes determined by remotely sensed habitat mapping from the Louisiana BICM Program (Enwright et al., 2020; Table S1 and S2). We used the habitat classification maps to assign a map-based inundation class to each bird observation at Caminada Headland. We categorized the habitat types into the following inundation classes: 1) intertidal-unvegetated (i.e., intertidal), 2) intertidal-vegetated (i.e., estuarine emergent marsh, or mangrove), 3) supratidal (i.e., beach, meadow, scrub/shrub, unvegetated dune, or unvegetated flat), and 4) water (water; Enwright et al., 2020; Table S2).

#### Results

##### Spectral Indices

Violin plots help show the variation of spectral indices for each of the four satellite-based habitat types (Figures S34–S35). In this study we opted to use the native Landsat and Sentinel-2 data to take advantage of the high resolution of Sentinel-2 data. Some differences in spectral indices were found by habitat class (Figures S34–S35). For the dates sampled for the violin plots, MSAVI tended to be higher for vegetated areas for Sentinel-2 compared to Landsat (Figures S34–S35). Other variables tended to be more similar between sensors.

##### Caminada Headland

Piping Plover – The relationship between distance to water and non-breeding abundance was bimodal. The highest abundances of non-breeding Piping Plovers were within ~10 m of water with another uptick in abundance >75 m from the water. The detailed habitat types with the highest abundance of individuals were estuarine emergent marsh and water; all other habitat types had relatively lower abundances, except scrub/shrub where no individuals were observed. Non-breeding abundance was highest in interior areas, closely followed by areas along the bay. In terms of MNDWI, we found high abundances in the tails of the distribution (MNDWI values <-0.5 and >0.5 but when examining the bulk of the data there is a slight increase between MNDWI values of ~-0.25 and 0.0 (Figure S1).

Foraging behaviors were most prevalent along the bay and Gulf portions of the headland, with few occurring within the interior. In terms of BI, we found high occurrences of foraging activity in the tails of the distribution (BI values <0.7 and > 0.9), but the bulk of the foraging occurrences spans a narrow window of brightness values (0.7–0.8) that show a positive relationship. Foraging tended to increase with MSAVI but was especially high in areas transitioning from bare ground to sporadic vegetative cover. Foraging generally increased as distance to water increased through 50 m but then decreased with increasing distance. MNDWI tended to have a positive relationship with foraging as soil moisture went from dry to open water (Figure S2). Opposite of the foraging analysis, we found most maintenance activities to occur within the interior area. Maintenance activities tended to decrease as BI and MNDWI values increased over the bulk of the data. In contrast, maintenance activities increased as MSAVI values increased. Maintenance activities slightly decreased between 0 m and 50 m from water but then increased from 50 m to 150 m from the water (Figure S3).

Red Knot –Aside from a small spike in numbers at the water's edge, we found the number of non-breeding Red Knots to increase with proximity from water. We found Red Knots abundance to decrease with increasing BI values until a BI value of ~0.76, then increase with brighter landcover (Figure S4). We found foraging activity to generally decrease as distance away from water increased, but foraging was relatively high within 50 m of water. Foraging generally had a positive relationship with both BI and MSAVI. Overall, foraging activity tended to decrease as MNDWI values increased, but in the area with the majority of the data (MNDWI values ~-0.3–0.0) we tended to see foraging increase with an increase in MSAVI. Most foraging took place on the Gulf-side of the headland as opposed to the bay side or interior areas (Figure S5). We found an opposite pattern between maintenance activities and distance to water when compared to foraging, such that individuals engaged in more maintenance activities as distance from water increased. Overall, maintenance activities decreased with BI, although there was a slight uptick between 0.75 and 0.80 where a majority of the data lie. Over most of the data, we found the prevalence of maintenance behaviors to decrease with MSAVI. There was a positive relationship between maintenance activity and MNDWI. Another opposing result when compared to foraging is that maintenance activities were more prevalent within areas classified as interior or bay compared to the Gulf-side of the headland (Figure S6).

Snowy Plover – Snowy Plover non-breeding abundance tended to increase as MNDWI values increased, whereas abundance tended to decrease as MSAVI values increased. The greatest abundance of Snowy Plover occurred at ~25 m from the water's edge, which decreased as distance from the water increased or decreased. In terms of habitat, we had the highest abundances in areas classified as unvegetated dune, beach, water, and unvegetated flat. Habitats classified as meadow, estuarine emergent marsh, and intertidal had relatively lower abundances, while no individuals were observed in areas classified as mangrove, scrub/shrub, or vegetated dune. Finally, abundance decreased as BI values approached 0.73 but then increased as BI values increased above 0.73 (Figure S7).

Foraging decreased as BI values increased. Foraging primarily took place on the bay-side of the headland, followed closely by the Gulf side; little foraging occurred in the interior. Foraging was highest at the water's edge with foraging activity generally decreasing with distance through 50 m, before then increasing between ~50–75 m. The majority of foraging observations were obtained in areas with MSAVI values between 0.0 and 0.1, which showed a general decrease in foraging as MSAVI values increased. Foraging tended to increase as MNDWI values increased with the majority of data from areas that would be considered irregularly to regularly flooded (Figure S8). Maintenance activities tended to increase as BI values, MNDWI values, and distance to water increased. Across the bulk of data, we found maintenance activities to decrease as MSAVI values increased to areas that are likely getting some vegetative cover. Opposite of the foraging analysis, we found maintenance activities to be most prominent in areas classified as interior (Figure S9).

Wilson's Plover –Breeding Wilson's Plover abundances generally increased with increasing BI values. We found most individuals to be in areas with MNDWI values between -0.5 to 0.0, with abundances increasing from -0.5 to about -0.2 then decreasing between -0.2 to 0.0. We found breeding abundance to generally decrease as MSAVI values increased, suggesting individuals used areas with little to no vegetative cover. Despite a peak in breeding abundance at the water's edge, we found breeding abundance tended to increase with distance away from water. In terms of habitat, those classified as intertidal, water, and unvegetated flat had the highest abundances of breeding Wilson's Plover (Figure S10). Similar to breeding abundances of Wilson's Plover, we found non-breeding abundance to generally decrease with MSAVI across most of the data and increase with distance away from water. We found abundance to be lowest at BI values below 0.7, peak around 0.7, and then slightly decrease but remain generally stable above 0.7. Abundance decreased as values of MNDWI values increased with few individuals being in areas of open water (Figure S11).

Breeding behaviors increased as MSAVI increased until ~0.2 before declining as vegetative cover increased. Breeding activities occurred most on the Gulf-side of the headland and least on the bay side. We also found breeding activities to occur least at the Bayou site and most at the West Beach site. Overall, breeding tended to increase with BI from 0.75 to 0.8 and then decreased at higher BI values. Breeding activities tended to increase with MNDWI with most data occurring from -0.5 to 0.0 (Figure S12). Foraging generally increased with MNDWI values. Foraging decreased between MSAVI values of 0.0–0.2, which includes most of the data. Foraging also tended to trend lower with increasing BI values with most data bounded by BI values of 0.7–0.8. Foraging was greatest nearest to water and decreased as distance to water increased, which also explains why foraging was greatest on the Gulf-side of the headland and lowest within the interior (Figure S13). The analysis found maintenance behaviors to decrease as MSAVI values increase. Opposite of the foraging analysis, we found most maintenance activities to occur within the headland's interior. We also found an overall negative relationship between maintenance and MNDWI between -0.5 and 0.0, which encompasses most of the data. With BI values, we found maintenance activities, besides a peak around ~0.65, activities generally increased between 0.7 and 0.8 where most of the data fell. We also found a negative relationship between maintenance activities and distance to water over the bulk of the data (0–100 m from water). Maintenance activities mainly occurred ~20 m to 140 m from water, which matches our finding that most maintenance activities occurred within the headland's interior (Figure S14).

Whiskey Island

American Oystercatcher – During the breeding season, our model found that the abundance of oystercatchers was higher prior to restoration compared to the during or post-restoration phases. Abundance generally increased with MSAVI values as well with peak abundance found in areas with values indicative of bare ground. The relationship between abundance of breeding individuals and MNDWI was such that most individuals were observed in areas with moderately dry conditions to areas experiencing regular inundation (i.e., intertidal habitats and emergent marsh); few individuals were observed in water bodies. Focusing on the majority of data, we found a peak in abundance in areas classified with high brightness values (~0.75–0.78) that are indicative of drier soils (Figure S15).

We found a positive relationship between non-breeding abundance and MNDWI values with most individuals in areas that experience regular inundation (i.e., intertidal habitats and emergent marsh) to open water. Most individuals were observed utilizing developed/shoreline protection areas as opposed to supratidal or unvegetated intertidal habitats; no individuals were observed in vegetated intertidal habitats during the non-breeding season. Oystercatchers tended to be in areas with bare soil primarily, based on MSAVI values, with abundance decreasing with increased vegetative cover. We found non-breeding abundance to have a negative relationship with brightness, such that based on BI values individuals tended to select less reflective habitats (e.g., wet sand; Figure S16).

In the logistic response to individuals exhibiting breeding behaviors analysis, the only meaningful variable was inundation class (80.8%); all other variables had <5% relative influence (CV: 0.86 ± 0.02). Most breeding behaviors were observed in supratidal habitats, followed by intertidal vegetated habitats; developed and intertidal unvegetated habitats had the least observations of individuals engaged in breeding behaviors (Figure S17). Foraging tended to increase as BI values increased with two peaks occurring around 0.71 and 0.84. Foraging tended to be highest in areas with no vegetation based on low MSAVI values (<0.1) and decrease as MSAVI increased. Foraging had a bimodal distribution with respect to MNDWI, such that foraging was lowest in areas with regular inundation and peaked in areas with relatively low (dry) and high (open water) MNDWI values (Figure S18). Opposite of the breeding behavior analysis, we found areas classified as developed to be where most maintenance activities occurred, while intertidal vegetated habitats were where maintenance activities were least likely to occur. We found maintenance activities to increase with BI until ~0.81 then decreases when approaching the brightest values (Figure S19).

Piping Plover – We found most non-breeding individuals to occupy the brightest areas (>0.75), which may be drier soils. We found abundance to increase with MSAVI values greater than zero, suggesting more birds in areas with sparse to moderate vegetative cover. Distance to water showed a bimodal distribution, where individuals were found relatively close to water (i.e., within 25 m) or relatively far from water (> ~70 m; Figure S20). Foraging increased with BI values between 0.40 and 0.65 and then generally decreased beyond 0.65. Foraging was highest in areas classified as intertidal vegetated, closely followed by developed and intertidal unvegetated; supratidal habitats had the least occurrence of foraging. Foraging occurrences decreased as distance to water increased between the water's edge and ~30 m, then increased at distances greater than ~30 m. Probability of foraging tended to increase as MNDWI values increased, suggesting more foraging in wetter areas. Most foraging occurred in areas that are likely bare soil or transitioning from bare soil to sporadic vegetation based on the MSAVI values (Figure S21). We found maintenance activities to be highest at low BI values, but there was a secondary peak when BI values were at 0.75. Opposite of foraging behaviors, we found most maintenance activities to take place in habitats that are classified as supratidal according to our inundation class, whereas all other classes illustrated low use for maintenance activities. Maintenance activities showed a bimodal distribution with distance to water, which shows that maintenance activities were greatest ~25 m from water and >100 m from water. Maintenance activities showed a positive relationship with MSAVI such that most maintenance activities occurred around at MSAVI values >0, which would be areas transitioning from bare soil to slight and moderate vegetation cover (Figure S22).

Red Knot – We found the highest abundance of non—breeding Red Knots between ~20–50 m from water. In terms of habitat types used by Red Knots, we found most individuals to be in habitats classified as beach and unvegetated flat. Red Knot abundance tended to decrease with an increase in BI values, such that most individuals were in areas with a BI value between 0.50 and 0.75. Finally, Red Knot abundance tended to increase as the soil transitioned from bare ground to sporadic or moderately vegetated ground cover (0.0–0.6 MSAVI; Figure S23). Foraging activity generally decreased as brightness increased, although there is a small peak in foraging activity at relatively bright habitats (~0.80). Foraging tended to decrease with increasing moisture levels based on MNDWI values. Foraging was highest along the water's edge (0–20 m from water), followed by a secondary peak in foraging activity ~50 m away from water. Foraging also showed a bimodal distribution relative to MSAVI where foraging was highest between -0.2 and -0.1, which would be indicative of areas that are bare soil, and areas with MSAVI values >0.1, which would be slight to moderate vegetative ground cover (Figure S24). Maintenance activities peaked when soil brightness was between ~0.72 and 0.78. Maintenance activities occurred mostly in areas with bare soil according to MSAVI values (~-0.1–0.15) and decreased as MSAVI values increased. Maintenance activities generally increased as MNDWI values increased. Finally, the relationship between maintenance activities and distance to water illustrated a bimodal distribution where there was an initial small peak in activity around 20 m from water and a larger peak in maintenance activities around 80 m from the water (Figure S25).

Snowy Plover – For Snowy Plovers, we found abundance to increase with BI values, suggesting more individuals occupying more reflective habitats (dry sand or white water). Similarly, we found abundance to increase as MSAVI values increased, suggesting higher abundances in areas with slight to moderate vegetative ground cover. MNDWI values also tended to have a positive relationship with abundance, suggesting individuals increased in abundance with soil moisture (Figure S26). Snowy Plover foraging tended to increase with MSAVI values until foraging activity peaked when MSAVI values were ~-0.1 before decreasing with greater MSAVI values. Foraging activity generally increased with BI values. Snowy Plovers foraging showed a negative relationship with MNDWI, suggesting foraging activity was highest in relatively dry habitats. Intertidal vegetated and unvegetated were the inundation classes with the most foraging, while no foraging occurred in areas classified as developed (Figure S27). Maintenance activities generally increased with MSAVI values with the largest peak in activity occurring in areas of bare soil just before bare ground transitions to some sporadic cover (~0–0.2). In terms of BI, maintenance activities generally increased with brightness values when considering the bulk of the data. The relationship between inundation class and maintenance activities was opposite of foraging considering most individuals engaged in maintenance activities in areas considered supratidal; no birds engaged in maintenance activities within areas classified as developed (Figure S28).

Wilson's Plover – We found breeding abundance to decrease as soil brightness increased. Wilson's Plover breeding abundance was highest when tides were falling or rising, but lowest during high or low tide. We observed the highest breeding abundances before restoration occurred, followed by the during restoration period; abundances were lowest after restoration occurred. Breeding abundances showed a bimodal distribution with MNDWI such that there was an initial small peak in abundance at MNDWI values of ~-0.4 (i.e., non-water areas) and a larger peak in areas considered open water. We also found breeding abundance to be greatest in habitats classified as meadow, scrub/shrub, shoreline protection, and unvegetated flat (Figure S29). Non-breeding abundance was highest when the tide was rising and lowest when the tide was falling or low. Non-breeding abundances showed a positive relationship with MSAVI, but the highest peak in abundance occurred around an MSAVI value of 0.1. In terms of habitat use, Wilson's Plovers were most observed in habitats classified as mangrove or unvegetated dune and least observed in habitats classified as intertidal. We did not observe non-breeding Wilson's Plover in habitats classified as bare land, scrub/shrub, shoreline protection, or vegetated dune. Finally, we found a positive relationship between non-breeding abundance and BI values with most individuals occurring between brightness values of ~0.65 to 0.8 (Figure S30).

When analyzing the relationship between MSAVI as well as BI with breeding behavior, we found a positive relationship — prevalence of breeding behavior generally increased with increasing values of MSAVI and BI. Breeding behaviors generally increased with MNDWI until MNDWI values of 0.5 then decreased. However, most breeding observations occurred in a narrow range of MNDWI values (~-0.25–0.0) that did not appear to increase or decrease with changing MNDWI values. Distance to water showed two peaks where breeding behaviors were highest (~100 m and 225 m) as well as a valley around 150 m when the prevalence of breeding behaviors appeared to be lowest (Figure S31). We found most foraging behaviors to be related to habitat classified as bare land, closely followed by water and herbaceous; few foraging observations occurred in woody habitats. In terms of MNDWI, we generally found foraging to increase as birds moved from drier habitats to open water; similar to breeding behaviors, most observations occurred within a narrow band of MNDWI values (~-0.25–0.10) that appeared to not increase or decrease with changing MNDWI values (Figure S32). We found maintenance activities to decrease as MSAVI values increased.

In terms of MNDWI, we generally found a positive relationship with maintenance activities. Maintenance activities showed a negative relationship with BI. Finally, maintenance activities were highest close to water and decreased as distance to water increased (Figure S33).

Literature Cited

Cowardin, L. M., Carter, V., Golet, F. C., & LaRoe, E. T. (1979). *Classification of wetlands and deepwater habitats of the United States* (No. Report FWS/ OBS–79/31; p. 131). U.S. Department of the Interior, Fish and Wildlife Service.

Enwright, N. M., SooHoo, W. M., Dugas, J. L., Conzelmann, C. P., Laurenzano, C., Lee, D. M., Mouton, K., & Stelly, S. J. (2020). *Louisiana Barrier Island Comprehensive Monitoring Program: Mapping habitats in beach, dune, and intertidal environments along the Louisiana Gulf of Mexico shoreline, 2008 and 2015–16* (U.S. Geological Survey Open-File Report Nos. 2020–1030; p. 57). U.S. Geological Survey. http://pubs.er.usgs.gov/publication/ofr20201030

Escadafal, R., Girard, M.-C., & Courault, D. (1989). Munsell soil color and soil reflectance in the visible spectral bands of landsat MSS and TM data. *Remote Sensing of Environment*, *27*(1), 37–46. https://doi.org/10.1016/0034-4257(89)90035-7

Fearnley, S., Brien, L., Martinez, L., Miner, M., Kulp, M., & Penland, S. (2009). *Chenier Plain, South-Central Louisiana, and Chandeleur Islands, Habitat mapping and change analysis 1996 to 2005, Part 1—Methods for habitat mapping and change analysis 1996 to 2005—Louisiana Barrier Island Comprehensive Monitoring Program (BICM) 5* (p. 11). University of New Orleans, Pontchartrain Institute for Environmental Sciences.

Homer, C., Dewitz, J., Yang, L., Jin, S., Danielson, P., Xian, G., Coulston, J., Herold, N., Wickham, J., & Megown, K. (2015). Completion of the 2011 National Land Cover Database for the Conterminous United States – Representing a Decade of Land Cover Change Information. *Photogrammetric Engineering and Remote Sensing*, *81*, 345–354.

Leatherman, S. P. (1979). *Barrier Island Handbook*. National Park Service, Cooperative Research Unit, The Environmental Institute, University of Massachusetts at Amherst.

Lucas, K. L., & Carter, G. A. (2010). Decadal Changes in Habitat-Type Coverage on Horn Island, Mississippi, U.S.A. *Journal of Coastal Research*, *26*(6), 1142–1148. https://doi.org/10.2112/JCOASTRES-D-09-00018.1

Psuty, N. P. (1989). An application of science to the management of coastal dunes along the Atlantic coast of the U.S.A. *Proceedings of the Royal Society of Edinburgh, Section B: Biological Sciences*, *96*, 289–307. https://doi.org/10.1017/S0269727000010988

Qi, J., Chehbouni, A., Huete, A. R., Kerr, Y. H., & Sorooshian, S. (1994). A modified soil adjusted vegetation index. *Remote Sensing of Environment*, *48*(2), 119–126. https://doi.org/10.1016/0034-4257(94)90134-1

Schulz, J. L., & Leberg, P. L. (2019). Factors Affecting Prey Availability and Habitat Use of Nonbreeding Piping Plovers (Charadrius melodus) in Coastal Louisiana. *Journal of Coastal Research*, *35*(4), 861–871. https://doi.org/10.2112/JCOASTRES-D-17-00147.1

Thurman, H. R., Enwright, N. M., Cheney, W. C., Dugas, J. L., Lee, D. M., & Jones, W. R. (2023). *Mapping habitats and shorelines pre-, during, and post-restoration on Caminada Headland and Whiskey Island, Louisiana, 2012–2020* (p. 51p.) [Final Report]. Louisiana Coastal Protection and Restoration Authority.

Xu, H. (2006). Modification of normalised difference water index (NDWI) to enhance open water features in remotely sensed imagery. *International Journal of Remote Sensing*, *27*(14), 3025–3033. https://doi.org/10.1080/01431160600589179

Tables

Table S1. Louisiana Barrier Island Comprehensive Monitoring (BICM) program's detailed habitat classification schemes. Note, while the forest class is part of the BICM detailed classification scheme, it is not present in these habitat maps. [NA, not applicable]

| **Detailed class** | **Description** | **Description source** |
|---|---|---|
| Beach | Beach habitat includes supratidal bare or sparsely vegetated areas (i.e., above the extreme high water springs tide level) located along coastlines with high wave energy (i.e., Gulf-facing shorelines). Vegetation cover is generally less than 30%. Beach transitions into dunes, meadow, or unvegetated flat where overwash is evident. Beach includes the backshore zone of a beach. | Modified from Cowardin et al., 1979 |
| Unvegetated dune | Dunes are supratidal features (i.e., above the extreme high water springs tide level) developed via Aeolian processes. Dunes are often located above typical storm water levels and have a well-defined relative elevation (i.e., upper slope or | Modified from Psuty, 1989 |

| | | |
|---|---|---|
| | ridge). Unvegetated dune includes dune habitat that has less than 10% vegetation cover. | |
| Vegetated dune | Dunes are supratidal features (i.e., above the extreme high water springs tide level) developed via Aeolian processes. Dunes are often located above typical storm water levels and have a well-defined relative elevation (i.e., upper slope or ridge). Vegetated dune includes dune habitat that has greater than 10% vegetation cover. | Modified from Psuty, 1989 |
| Unvegetate d flat | Unvegetated barrier flat includes flat or gently sloping supratidal unvegetated or sparsely vegetated areas (i.e., areas located above extreme high water springs tide level) that are located on the backslope of dunes, unvegetated washover fans, and along low-energy shorelines. Vegetation coverage should be generally less than 30%. | Modified from Leatherman , 1979 |
| Meadow | Meadow includes supratidal areas (i.e., above the extreme high water springs tide level) with sparse to dense herbaceous vegetation located in areas leading up to dunes and on the barrier flat (i.e., backslope of dunes and supratidal, back-barrier habitat). Vegetation coverage should generally be greater than 30%. Classification of meadow habitat is restricted by geomorphic settings. Meadow is reserved for areas located on barrier flats of barrier islands, backslopes of dunes, transitional vegetated areas in dune/beach habitats. | Modified from Lucas & Carter, 2010 |
| Intertidal | Intertidal includes bare or sparsely vegetated areas located between the extreme low water springs and extreme high water springs tide levels. Vegetation cover should generally be less than 30%. Intertidal includes the foreshore zone of a beach. | Cowardin et al., 1979 |

| | | |
|---|---|---|
| Estuarine emergent marsh | Estuarine emergent marsh includes intertidal saline emergent marsh (i.e., located above extreme low water springs and below extreme high water springs tide levels) and supratidal brackish emergent marsh. Vegetation cover should be generally 30% or greater cover by erect, rooted, herbaceous hydrophytes. Note, supratidal emergent vegetation that is located on the backslopes of dunes will be classified as meadow. | Cowardin et al., 1979 |
| Mangrove | Mangrove habitat includes areas with black mangrove (*Avicennia germinans*). Mangrove vegetation coverage should generally be greater than 30%. | NA |
| Bare land | Bare land includes bare or sparsely vegetated areas that are often located above typical storm water levels and are associated with unvegetated spoil or inland ridges. Vegetation cover should generally be less than 30%. | Modified from Fearnley et al., 2009 |
| Grassland | Grassland includes upland areas covered by herbaceous vegetation often located above typical storm water levels and are associated with inland spoil banks with herbaceous vegetation, freshwater emergent marsh, and upland areas along the mainland in the Chenier Plain BICM regions. | Modified from Homer et al., 2015 |
| Scrub/shrub | Scrub/shrub includes areas where woody vegetation height is greater than about 0.5 m, but less than 6 m. Woody vegetation coverage should generally be greater than 30%. | Cowardin et al., 1979 |
| Forest | Forest includes areas where woody vegetation height is greater than 6 m. Woody vegetation coverage should generally be greater than 30%. | Cowardin et al., 1979 |

| | | |
|---|---|---|
| Shoreline protection | Shoreline protection includes any material used to protect shorelines against erosion (e.g., breakwater, groins, and jetties). | Fearnley et al., 2009 |
| Developed | Developed includes areas dominated by constructed materials (i.e., transportation infrastructure, and residential and commercial areas) and open developed areas. | Modified from Homer et al., 2015 |
| Water | Water includes areas of open water with generally less than 30% cover of vegetation. | Modified from Cowardin et al., 1979 |

Table S2. Crosswalk for ortho-based habitat classes and Barrier Island Comprehensive Monitoring (BICM) inundation zone-based classes.

| **Inundation zone-based class** | **Ortho-based habitat class** |
|---|---|
| Water | Water |
| Intertidal-unvegetated | Intertidal |
| Intertidal-vegetated | Estuarine emergent marsh and Mangrove |
| Supratidal | Bare land, Beach, Forest, Grassland, Meadow, Scrub/shrub, Unvegetated dune, Unvegetated flat, and Vegetated dune |
| Developed/shoreline protection | Developed and Shoreline protection |

Table S3. Aerial orthoimagery used to produce habitat maps of Caminada Headland in 2012, 2015, and 2017–2019. The 2019 imagery had a resolution of 0.6 m but was resampled to 1 m to create the maps. [CRMS, Coastwide Reference Monitoring System; NAIP, National Agriculture Imagery Program]

| **Year** | **Image date(s)** | **Image source** | **Image resolution (m)** | **Restoration status** |
|---|---|---|---|---|
| 2012 | 11/07/2012; 11/13/2012 | CRMS | 1 | Pre-restoration |
| 2015 | 04/30/2015 | NAIP | 1 | During restoration |
| 2017 | 09/08/2017; 09/12/2017 | NAIP | 1 | Post-restoration |
| 2018 | 11/16/2018 | CRMS | 1 | Post-restoration |
| 2019 | 08/30/2019 | NAIP | 0.6 | Post-restoration |

Table S4. Aerial orthoimagery used to produce habitat maps of Whiskey Island in 2012, 2015, and 2017–2019. The 2019 imagery had a resolution of 0.6 m but was resampled to 1 m to create the maps. [CRMS, Coastwide Reference Monitoring System; NAIP, National Agriculture Imagery Program]

| Year | Image date(s) | Image source | Image resolution (m) | Restoration status |
|---|---|---|---|---|
| 2012 | 11/07/2012 | CRMS | 1 | Pre-restoration |
| 2015 | 08/26/2015 | NAIP | 1 | Pre-restoration |
| 2017 | 09/10/2017; 09/11/2017 | NAIP | 1 | During restoration |
| 2018 | 11/15/2018 | CRMS | 1 | Post-restoration |
| 2019 | 08/30/2019 | NAIP | 0.6 | Post-restoration |

Table S5. Landsat 8 and Sentinel-2 satellite imagery of Caminada Headland from 2013–2019, and the closest avian survey dates to each image.

| Image source | Image number | Image date | Avian survey date(s) |
|---|---|---|---|
| Landsat | 1* | 3/29/2013 | 1/11/2013–4/5/2013 |
| Landsat | 2 | 6/25/2013 | 5/6/2013–9/25/2013 |
| Landsat | 3 | 12/18/2013 | 10/9/2013–12/18/2013 |
| Landsat | 4 | 1/19/2014 | 1/15/2014–2/27/2014 |
| Landsat | 5 | 4/9/2014 | 3/12/2014–6/5/2014 |
| Landsat | 7 | 8/15/2014 | 7/30/2014–9/24/2014 |
| Landsat | 8 | 11/19/2014 | 10/8/2014–12/17/2014 |
| Landsat | 9 | 2/7/2015 | 1/14/2015–2/26/2015 |
| Landsat | 10* | 3/27/2015 | 3/16/2015–6/10/2015 |
| Landsat | 11 | 9/19/2015 | 7/29/2015–9/23/2015 |
| Sentinel | 2* | 10/6/2015 | 10/7/2015–10/21/2015 |
| Landsat | 12 | 11/22/2015 | 11/4/2015–11/19/2015 |
| Landsat | 13 | 12/8/2015 | 12/2/2015 |
| Sentinel | 3 | 12/18/2015 | 12/15/2015 |
| Sentinel | 4 | 1/4/2016 | 1/13/2016 |
| Sentinel | 6 | 1/24/2016 | 1/22/2016–1/25/2016 |
| Landsat | 14 | 2/10/2016 | 2/10/2016 |
| Landsat | 15 | 2/26/2016 | 2/24/2016 |
| Landsat | 16 | 3/13/2016 | 3/8/2016–4/6/2016 |

| | | | |
|---|---|---|---|
| Sentinel | 8 | 5/23/2016 | 4/20/2016–6/8/2016 |
| Landsat | 17* | 10/7/2016 | 7/27/2016–10/19/2016 |
| Sentinel | 9* | 11/19/2016 | 11/2/2016–11/16/2016 |
| Sentinel | 10 | 11/29/2016 | 12/1/2016 |
| Landsat | 18 | 12/10/2016 | 12/14/2016 |
| Sentinel | 11 | 1/8/2017 | 2/6/2017–2/9/2017 |
| Landsat | 19 | 3/16/2017 | 2/22/2017–3/22/2017 |
| Landsat | 20 | 4/1/2017 | 4/5/2017–4/19/2017 |
| Sentinel | 17 | 5/8/2017 | 5/10/2017–6/15/2017 |
| Sentinel | 18 | 7/25/2017 | 7/26/2017 |
| Sentinel | 19* | 8/16/2017 | 8/9/2017–8/23/2017 |
| Landsat | 21* | 9/8/2017 | 9/6/2017–9/19/2017 |
| Sentinel | 20 | 10/5/2017 | 10/4/2017 |
| Landsat | 22 | 10/26/2017 | 10/18/2017–11/1/2017 |
| Sentinel | 23 | 11/19/2017 | 11/15/2017 |
| Landsat | 24 | 11/27/2017 | 11/29/2017 |
| Sentinel | 24 | 12/12/2017 | 12/13/2017 |
| Sentinel | 25 | 1/3/2018 | 1/9/2018 |
| Sentinel | 26 | 1/23/2018 | 1/24/2018 |
| Landsat | 26 | 1/30/2018 | 2/6/2018 |
| Sentinel | 28 | 3/4/2018 | 2/20/2018–3/6/2018 |
| Sentinel | 30 | 3/22/2018 | 3/20/2018–4/4/2018 |

| | | | |
|---|---|---|---|
| Sentinel | 32 | 4/23/2018 | 4/17/2018 |
| Sentinel | 33 | 5/8/2018 | 5/1/2018–6/5/2018 |
| Sentinel | 35 | 7/22/2018 | 7/24/2018–8/7/2018 |
| Sentinel | 36* | 9/15/2018 | 8/21/2018–9/25/2018 |
| Landsat | 27* | 10/13/2018 | 10/2/2018–12/4/2018 |
| Sentinel | 41 | 1/28/2019 | 12/11/2018–1/29/2019 |
| Sentinel | 42 | 2/7/2019 | 2/5/2019–2/19/2019 |
| Landsat | 28 | 3/6/2019 | 3/7/2019 |
| Landsat | 29* | 3/22/2019 | 3/19/2019 |
| Sentinel | 43* | 4/3/2019 | 4/3/2019–6/5/2019 |

*Indicates if the satellite-based map was included in the accuracy assessment.

Table S6. Landsat 8 and Sentinel-2 satellite imagery of Whiskey Island from 2013–2020, and the closest avian survey dates to each image.

| Image source | Image number | Image date | Avian survey date(s) |
|---|---|---|---|
| Landsat | 1* | 3/29/2013 | 8/7/2012–3/8/2013 |
| Landsat | 2 | 6/25/2013 | 5/8/2013–9/17/2013 |
| Landsat | 3 | 12/18/2013 | 10/22/2013–12/17/2013 |
| Landsat | 4 | 1/19/2014 | 1/31/2014–2/28/2014 |
| Landsat | 5 | 4/9/2014 | 3/19/2014–5/4/2014 |
| Landsat | 6 | 7/30/2014 | 6/4/2014–7/31/2014 |
| Landsat | 7 | 8/15/2014 | 8/21/2014–9/17/2014 |
| Landsat | 8 | 11/19/2014 | 10/6/2014–12/17/2014 |
| Landsat | 9 | 2/7/2015 | 1/6/2015–2/11/2015 |
| Landsat | 10 | 3/27/2015 | 3/4/2015–5/29/2015 |
| Sentinel | 1* | 8/10/2015 | 6/3/2015–8/27/2015 |
| Landsat | 11* | 9/19/2015 | 9/10/2015–10/6/2015 |
| Landsat | 12 | 11/22/2015 | 10/22/2015–11/30/2015 |
| Landsat | 13 | 12/8/2015 | 12/15/2015–12/29/2015 |
| Sentinel | 5 | 1/17/2016 | 1/12/2016–1/25/2016 |
| Landsat | 14 | 2/10/2016 | 2/11/2016 |
| Landsat | 15 | 2/26/2016 | 2/26/2016 |

| | | | |
|---|---|---|---|
| Landsat | 16 | 3/13/2016 | 3/6/2016–4/3/2016 |
| Sentinel | 7 | 5/6/2016 | 4/12/2016–7/21/2016 |
| Landsat | 17* | 10/7/2016 | 7/28/2016–11/3/2016 |
| Landsat | 18 | 12/10/2016 | 11/16/2016–12/27/2016 |
| Sentinel | 12* | 1/31/2017 | 1/9/2017–1/24/2017 |
| Sentinel | 13 | 2/10/2017 | 2/8/2017 |
| Sentinel | 14 | 3/2/2017 | 2/23/2017–3/9/2017 |
| Sentinel | 15 | 3/22/2017 | 3/23/2017–3/23/2017 |
| Landsat | 20 | 4/1/2017 | 4/4/2017–4/19/2017 |
| Sentinel | 16 | 5/1/2017 | 4/24/2017–6/8/2017 |
| Sentinel | 18 | 7/25/2017 | 6/15/2017–8/8/2017 |
| Landsat | 21* | 9/8/2017 | 8/23/2017–9/20/2017 |
| Sentinel | 21* | 10/8/2017 | 10/5/2017 |
| Sentinel | 22 | 10/18/2017 | 10/18/2017 |
| Landsat | 22 | 10/26/2017 | 10/31/2017 |
| Landsat | 23 | 11/11/2017 | 11/14/2017 |
| Landsat | 24 | 11/27/2017 | 11/28/2017 |
| Sentinel | 24 | 12/12/2017 | 12/13/2017–12/29/2017 |
| Landsat | 25 | 1/14/2018 | 1/10/2018 |
| Landsat | 26 | 1/30/2018 | 1/24/2018–2/8/2018 |
| Sentinel | 27* | 2/20/2018 | 2/22/2018 |

| | | | |
|---|---|---|---|
| Sentinel | 29 | 3/12/2018 | 3/7/2018–3/13/2018 |
| Sentinel | 30 | 3/22/2018 | 3/22/2018–3/26/2018 |
| Sentinel | 31 | 4/11/2018 | 4/5/2018–4/16/2018 |
| Sentinel | 34 | 5/11/2018 | 4/25/2018–5/30/2018 |
| Landsat | 27 | 10/13/2018 | 8/14/2018–10/5/2018 |
| Sentinel | 37 | 10/18/2018 | 10/30/2018 |
| Sentinel | 38* | 11/17/2018 | 11/16/2018–11/28/2018 |
| Sentinel | 39 | 12/22/2018 | 12/11/2018 |
| Sentinel | 40 | 1/21/2019 | 2/13/2019 |
| Landsat | 28 | 3/6/2019 | 3/7/2019 |
| Landsat | 29 | 3/22/2019 | 4/10/2019 |
| Sentinel | 44 | 5/16/2019 | 4/26/2019–7/3/2019 |
| Sentinel | 45* | 8/29/2019 | 7/18/2019–10/23/2019 |
| Landsat | 30* | 1/20/2020 | 11/7/2019–2/28/2020 |
| Landsat | 31 | 3/24/2020 | 3/5/2020–3/18/2020 |
| Sentinel | 46 | 5/5/2020 | 5/6/2020 |
| Landsat | 32 | 5/11/2020 | 5/13/2020–5/28/2020 |
| Landsat | 33 | 6/12/2020 | 6/4/2020–6/26/2020 |
| Landsat | 34 | 7/14/2020 | 7/2/2020–8/19/2020 |

*Indicates if the satellite-based map was included in the accuracy assessment.

Table S7. Indices used for satellite-based habitat map development from 2013–2020 for Whiskey Island and Caminada Headland, Louisiana. NIR: the near-infrared band of the orthoimagery; RED: the red band of the imagery; GREEN: the green band of the imagery; SWIR: the shortwave infrared band of the imagery.

| Index | Formula | Interpretation | Source |
|---|---|---|---|
| Modified soil adjusted vegetation index (MSAVI) | $(2 * \mathrm{NIR} + 1 - \sqrt{((2 * \mathrm{NIR} + 1)^2 - 8 * (\mathrm{NIR} - \mathrm{RED})))}$ /2) | Measure of greenness; positive relationship between value and greenness | Qi et al., 1994 |
| Modified normalized difference water index (MNDWI) | $(\mathrm{GREEN} - \mathrm{SWIR})/(\mathrm{GREEN} + \mathrm{SWIR})$ | Measure of wetness; positive relationship between value and wetness | Xu, 2006 |
| Brightness index (BI) | $\sqrt{\frac{\frac{\mathrm{RED}^2}{\mathrm{GREEN}^2}}{2}}$ | Brightness of soil and substrate; positive relationship between value and brightness | Escadafal et al., 1989 |

Figures

Figure S1. Partial dependence plots of variables predicting Piping Plover abundance during the non-breeding seasons on Caminada Headland, Louisiana, USA from a boosted regression tree model. Y-axes are centered to have a zero mean over the data distribution. The relative influence (percent) of each predictor variable is shown in parentheses. Rug plots along the X-axis of each continuous variable plot illustrates the distribution of the data. Abbreviated labels on the X-axis of Detailed Habitat Class include Estuarine Emergent Marsh (E.E. Marsh), Scrub/Shrub (Scrb/Shrb), Unvegetated Dune (U. Dune), Unvegetated Flat (U. Flat), and Vegetated Dune (V. Dune). Abbreviated labels on the X-axis of Inundation Class include Intertidal Unvegetated (Intertidal Unveg.) and Intertidal Vegetated (Intertidal Veg.). Abbreviated labels on the X-axis of Site include East Beach (E. Beach), West Belle Pass (W. Belle Pass), and West Beach (W. Beach). MNDWI indicates Modified Normalized Difference Water Index and MSAVI indicates Modified Soil Adjusted Vegetation Index.

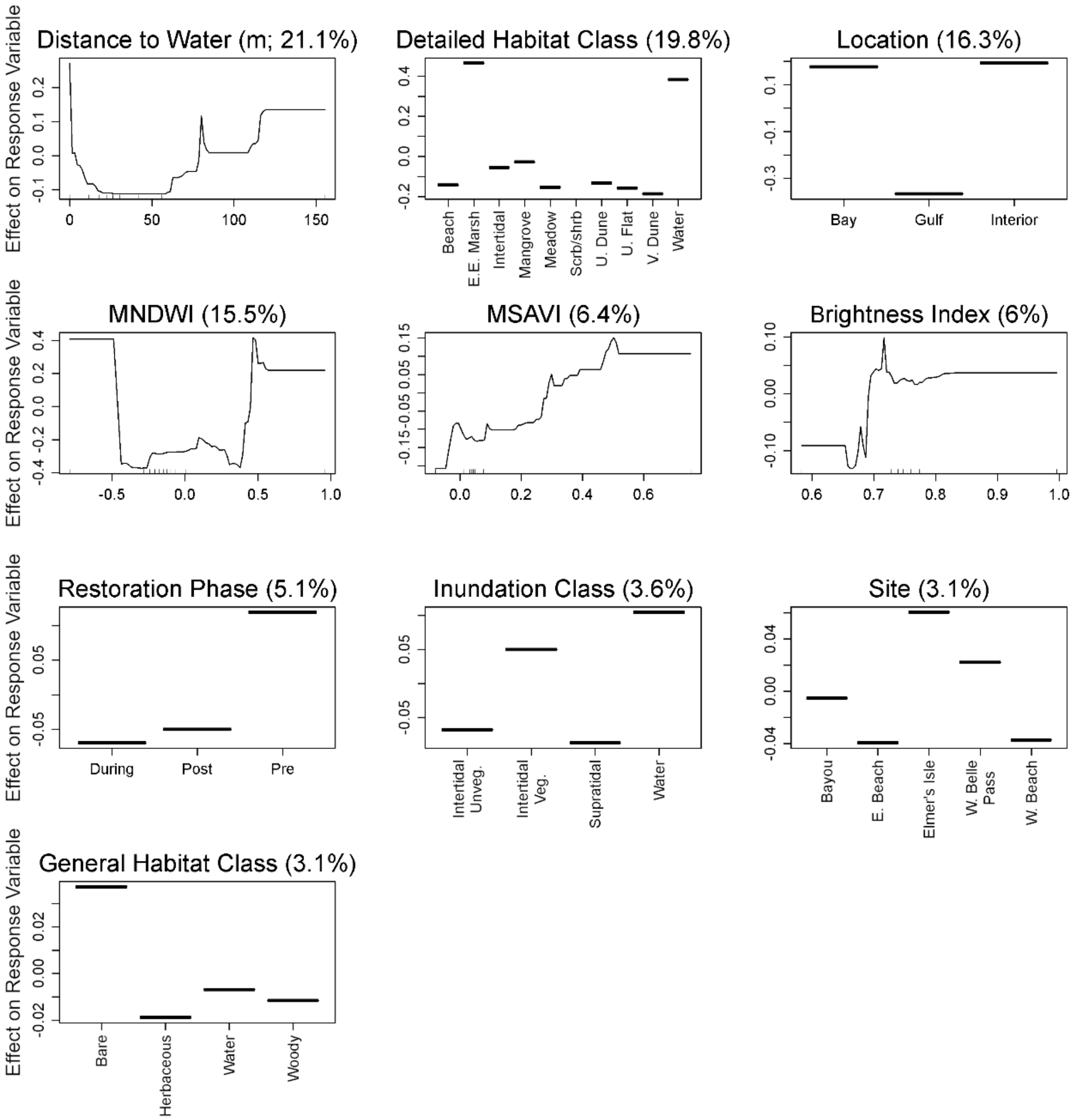
Distance to Water (m; 21.1%)
Detailed Habitat Class (19.8%)
Location (16.3%)
MNDWI (15.5%)
MSAVI (6.4%)
Brightness Index (6%)
Restoration Phase (5.1%)
Inundation Class (3.6%)
Site (3.1%)
General Habitat Class (3.1%)
Effect on Response Variable
Beach
E.E. Marsh
Intertidal
Mangrove
Meadow
Scrb/shrb
U. Dune
U. Flat
V. Dune
Water
Bay
Gulf
Interior
During
Post
Pre
Intertidal Unveg.
Intertidal Veg.
Supratidal
Water
Bayou
E. Beach
Elmer's Isle
W. Belle Pass
W. Beach
Bare
Herbaceous
Water
Woody

Figure S2. Partial dependence plots of variables predicting occurrence of Piping Plover foraging behaviors during the non-breeding seasons on Caminada Headland, Louisiana, USA from a boosted regression tree model. Y-axes are centered to have a zero mean over the data distribution. The relative influence (percent) of each predictor variable is shown in parentheses. Rug plots along the X-axis of each continuous variable plot illustrates the distribution of the data. Abbreviated labels on the X-axis of Detailed Habitat Class include Estuarine Emergent Marsh (E.E. Marsh), Scrub/Shrub (Scrb/Shrb), Unvegetated Dune (U. Dune), Unvegetated Flat (U. Flat), and Vegetated Dune (V. Dune). Abbreviated labels on the X-axis of Inundation Class include Intertidal Unvegetated (Intertidal Unveg.) and Intertidal Vegetated (Intertidal Veg.). Abbreviated labels on the X-axis of Site include East Beach (E. Beach), West Belle Pass (W. Belle Pass), and West Beach (W. Beach). MNDWI indicates Modified Normalized Difference Water Index and MSAVI indicates Modified Soil Adjusted Vegetation Index.

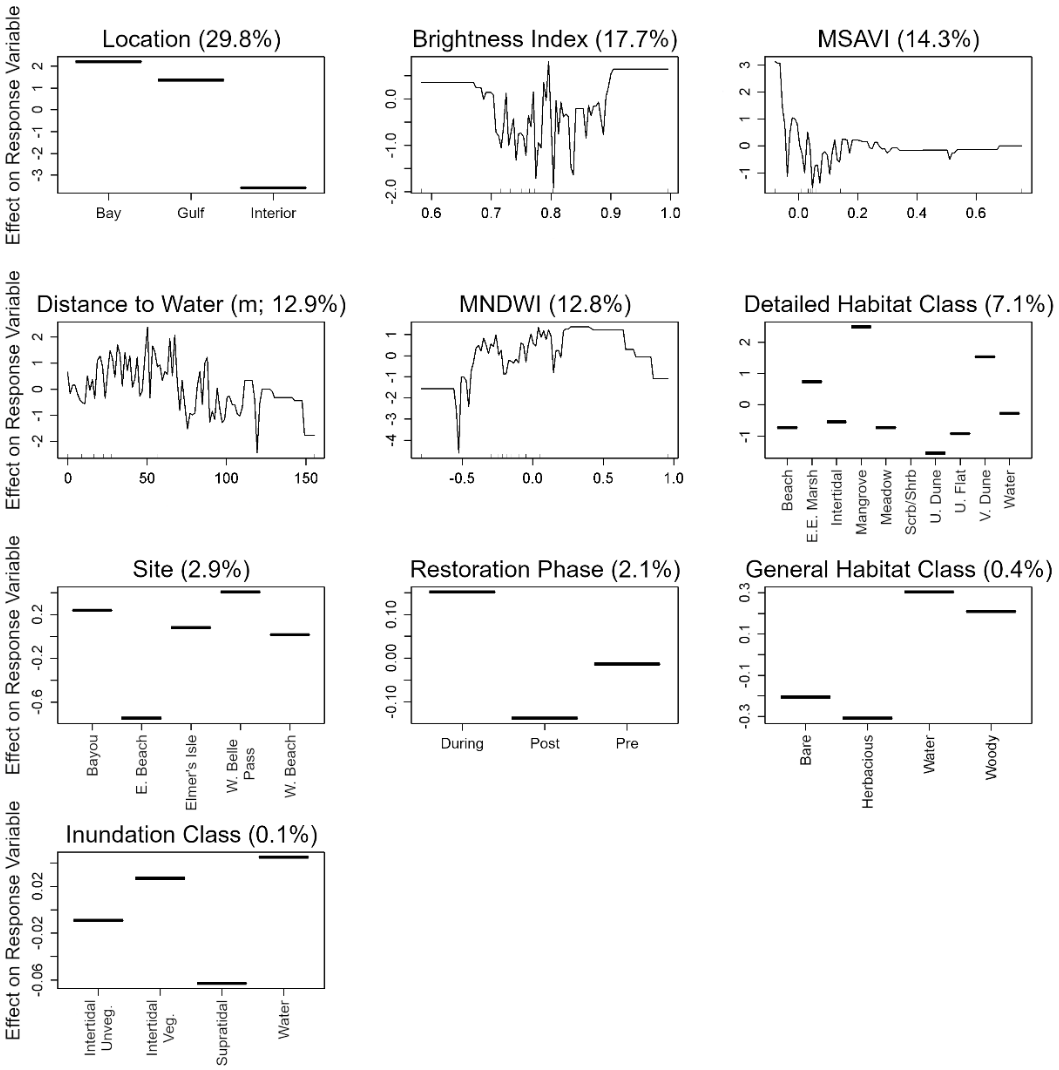
Location (29.8%)
Bay
Gulf
Interior
Brightness Index (17.7%)
MSAVI (14.3%)
Distance to Water (m; 12.9%)
MNDWI (12.8%)
Detailed Habitat Class (7.1%)
Beach
E.E. Marsh
Intertidal
Mangrove
Meadow
Scrb/Shrb
U. Dune
U. Flat
V. Dune
Water
Site (2.9%)
Bayou
E. Beach
Elmer's Isle
W. Belle Pass
W. Beach
Restoration Phase (2.1%)
During
Post
Pre
General Habitat Class (0.4%)
Bare
Herbacious
Water
Woody
Inundation Class (0.1%)
Intertidal Unveg.
Intertidal Veg.
Supratidal
Water
Effect on Response Variable

Figure S3. Partial dependence plots of variables predicting occurrence of Piping Plover maintenance behaviors during the non-breeding seasons on Caminada Headland, Louisiana, USA from a boosted regression tree model. Y-axes are centered to have a zero mean over the data distribution. The relative influence (percent) of each predictor variable is shown in parentheses. Rug plots along the X-axis of each continuous variable plot illustrates the distribution of the data. Abbreviated labels on the X-axis of Detailed Habitat Class include Estuarine Emergent Marsh (E.E. Marsh), Scrub/Shrub (Scrb/Shrb), Unvegetated Dune (U. Dune), Unvegetated Flat (U. Flat), and Vegetated Dune (V. Dune). Abbreviated labels on the X-axis of Inundation Class include Intertidal Unvegetated (Intertidal Unveg.) and Intertidal Vegetated (Intertidal Veg.). Abbreviated labels on the X-axis of Site include East Beach (E. Beach), West Belle Pass (W. Belle Pass), and West Beach (W. Beach). MNDWI indicates Modified Normalized Difference Water Index and MSAVI indicates Modified Soil Adjusted Vegetation Index.

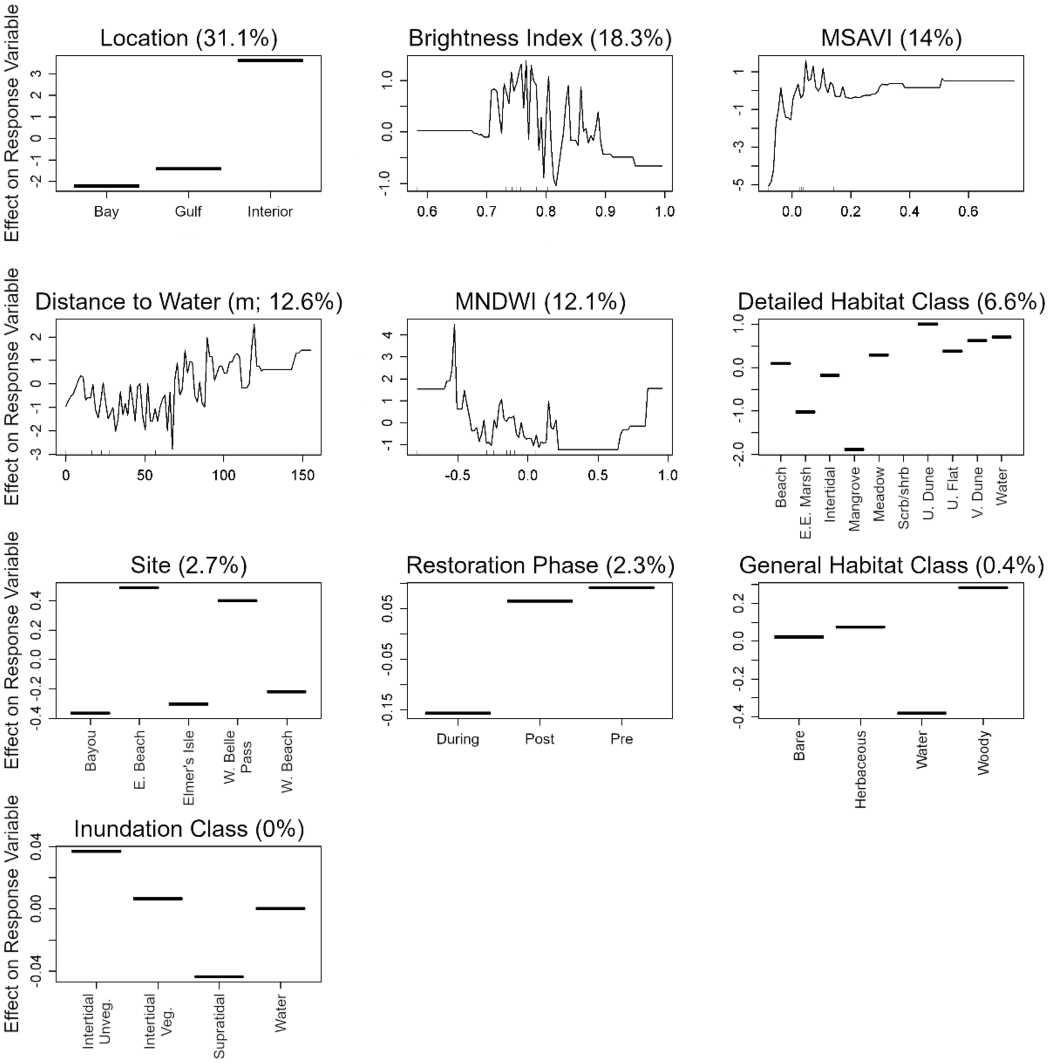
Location (31.1%)
Brightness Index (18.3%)
MSAVI (14%)
Distance to Water (m; 12.6%)
MNDWI (12.1%)
Detailed Habitat Class (6.6%)
Site (2.7%)
Restoration Phase (2.3%)
General Habitat Class (0.4%)
Inundation Class (0%)
Effect on Response Variable
Bay
Gulf
Interior
Beach
E.E. Marsh
Intertidal
Mangrove
Meadow
Scrb/shrb
U. Dune
U. Flat
V. Dune
Water
Bayou
E. Beach
Elmer's Isle
W. Belle Pass
W. Beach
During
Post
Pre
Bare
Herbaceous
Water
Woody
Intertidal Unveg.
Intertidal Veg.
Supratidal
Water

Figure S4. Partial dependence plots of variables predicting Red Knot abundance during the non-breeding seasons on Caminada Headland, Louisiana, USA from a boosted regression tree model. Y-axes are centered to have a zero mean over the data distribution. The relative influence (percent) of each predictor variable is shown in parentheses. Rug plots along the X-axis of each continuous variable plot illustrates the distribution of the data. Abbreviated labels on the X-axis of Detailed Habitat Class include Estuarine Emergent Marsh (E.E. Marsh), Scrub/Shrub (Scrb/Shrb), Unvegetated Dune (U. Dune), Unvegetated Flat (U. Flat), and Vegetated Dune (V. Dune). Abbreviated labels on the X-axis of Inundation Class include Intertidal Unvegetated (Intertidal Unveg.) and Intertidal Vegetated (Intertidal Veg.). Abbreviated labels on the X-axis of Site include East Beach (E. Beach), West Belle Pass (W. Belle Pass), and West Beach (W. Beach). MNDWI indicates Modified Normalized Difference Water Index and MSAVI indicates Modified Soil Adjusted Vegetation Index.

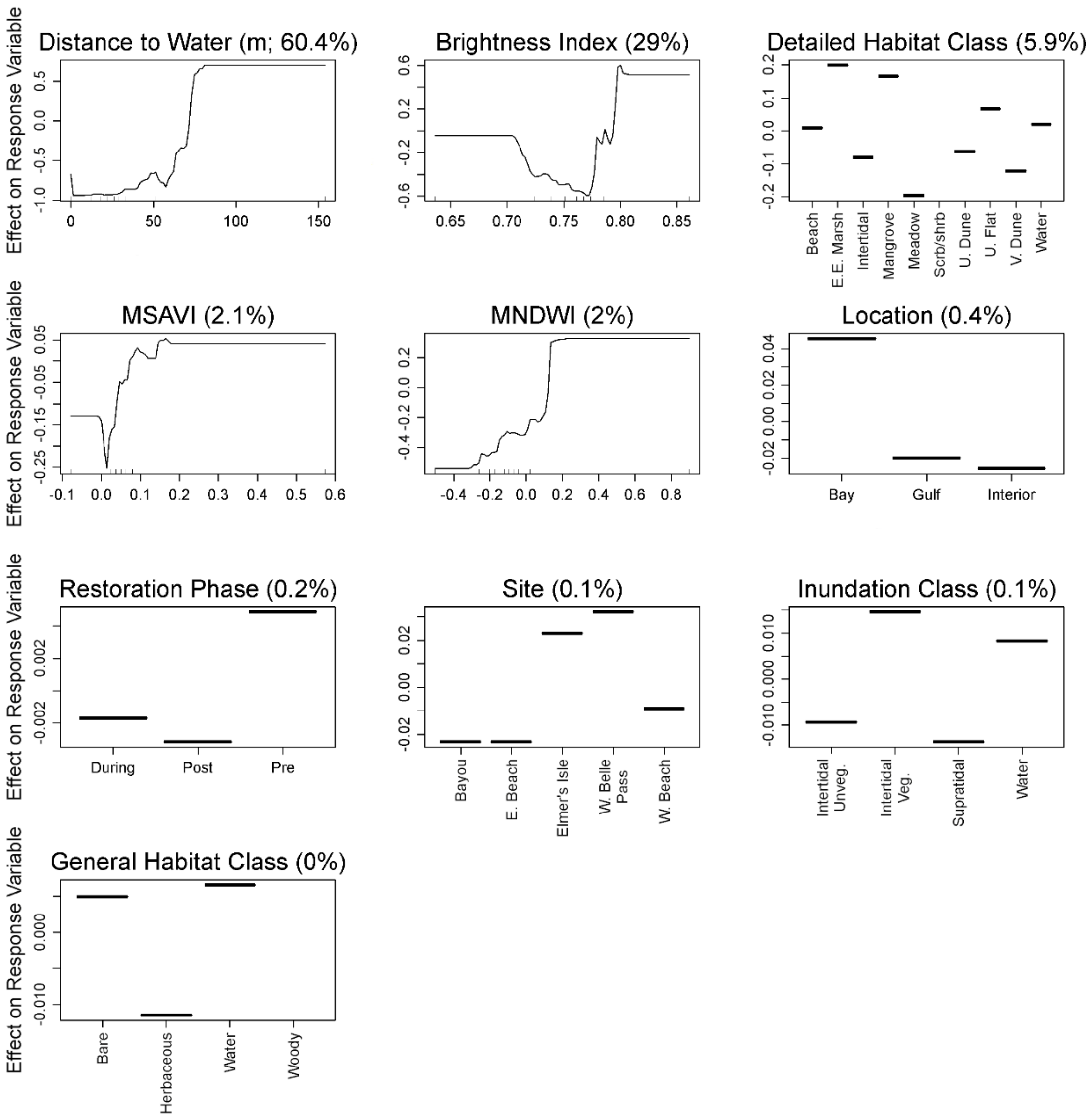
Effect on Response Variable
Distance to Water (m; 60.4%)
Brightness Index (29%)
Detailed Habitat Class (5.9%)
Beach
E.E. Marsh
Intertidal
Mangrove
Meadow
Scrb/shrb
U. Dune
U. Flat
V. Dune
Water
MSAVI (2.1%)
MNDWI (2%)
Location (0.4%)
Bay
Gulf
Interior
Restoration Phase (0.2%)
During
Post
Pre
Site (0.1%)
Bayou
E. Beach
Elmer's Isle
W. Belle Pass
W. Beach
Inundation Class (0.1%)
Intertidal Unveg.
Intertidal Veg.
Supratidal
Water
General Habitat Class (0%)
Bare
Herbaceous
Water
Woody

Figure S5. Partial dependence plots of variables predicting occurrence of Red Knot foraging behaviors during the non-breeding seasons on Caminada Headland, Louisiana, USA from a boosted regression tree model. Y-axes are centered to have a zero mean over the data distribution. The relative influence (percent) of each predictor variable is shown in parentheses. Rug plots along the X-axis of each continuous variable plot illustrates the distribution of the data. Abbreviated labels on the X-axis of Detailed Habitat Class include Estuarine Emergent Marsh (E.E. Marsh), Scrub/Shrub (Scrb/Shrb), Unvegetated Dune (U. Dune), Unvegetated Flat (U. Flat), and Vegetated Dune (V. Dune). Abbreviated labels on the X-axis of Inundation Class include Intertidal Unvegetated (Intertidal Unveg.) and Intertidal Vegetated (Intertidal Veg.). Abbreviated labels on the X-axis of Site include East Beach (E. Beach), West Belle Pass (W. Belle Pass), and West Beach (W. Beach). MNDWI indicates Modified Normalized Difference Water Index and MSAVI indicates Modified Soil Adjusted Vegetation Index.

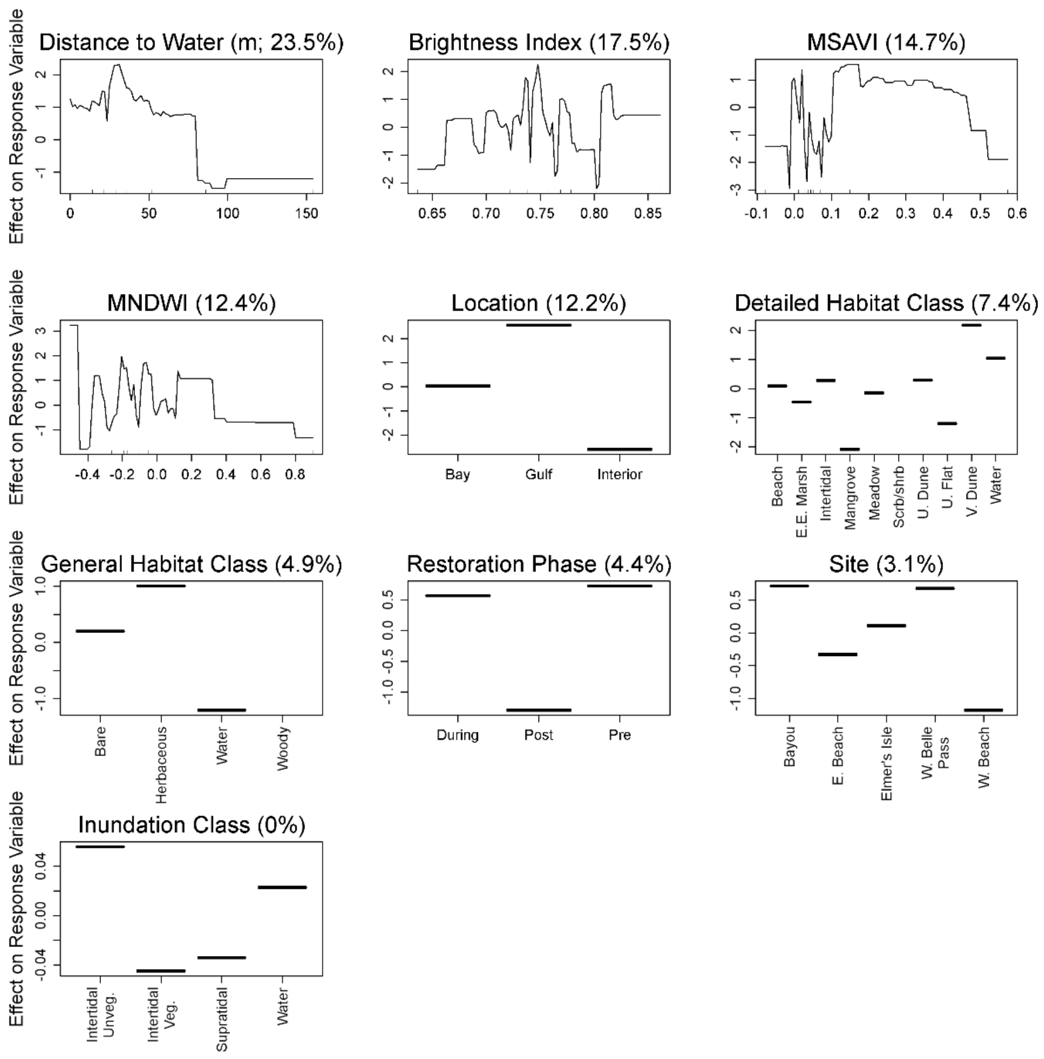
Distance to Water (m; 23.5%)
Brightness Index (17.5%)
MSAVI (14.7%)
MNDWI (12.4%)
Location (12.2%)
Detailed Habitat Class (7.4%)
General Habitat Class (4.9%)
Restoration Phase (4.4%)
Site (3.1%)
Inundation Class (0%)
Effect on Response Variable
Bay
Gulf
Interior
Beach
E.E. Marsh
Intertidal
Mangrove
Meadow
Scrb/shrb
U. Dune
U. Flat
V. Dune
Water
Bare
Herbaceous
Water
Woody
During
Post
Pre
Bayou
E. Beach
Elmer's Isle
W. Belle Pass
W. Beach
Intertidal Unveg.
Intertidal Veg.
Supratidal
Water

Figure S6. Partial dependence plots of variables predicting occurrence of Red Knot maintenance behaviors during the non-breeding seasons on Caminada Headland, Louisiana, USA from a boosted regression tree model. Y-axes are centered to have a zero mean over the data distribution. The relative influence (percent) of each predictor variable is shown in parentheses. Rug plots along the X-axis of each continuous variable plot illustrates the distribution of the data. Abbreviated labels on the X-axis of Detailed Habitat Class include Estuarine Emergent Marsh (E.E. Marsh), Scrub/Shrub (Scrb/Shrb), Unvegetated Dune (U. Dune), Unvegetated Flat (U. Flat), and Vegetated Dune (V. Dune). Abbreviated labels on the X-axis of Inundation Class include Intertidal Unvegetated (Intertidal Unveg.) and Intertidal Vegetated (Intertidal Veg.). Abbreviated labels on the X-axis of Site include East Beach (E. Beach), West Belle Pass (W. Belle Pass), and West Beach (W. Beach). MNDWI indicates Modified Normalized Difference Water Index and MSAVI indicates Modified Soil Adjusted Vegetation Index.

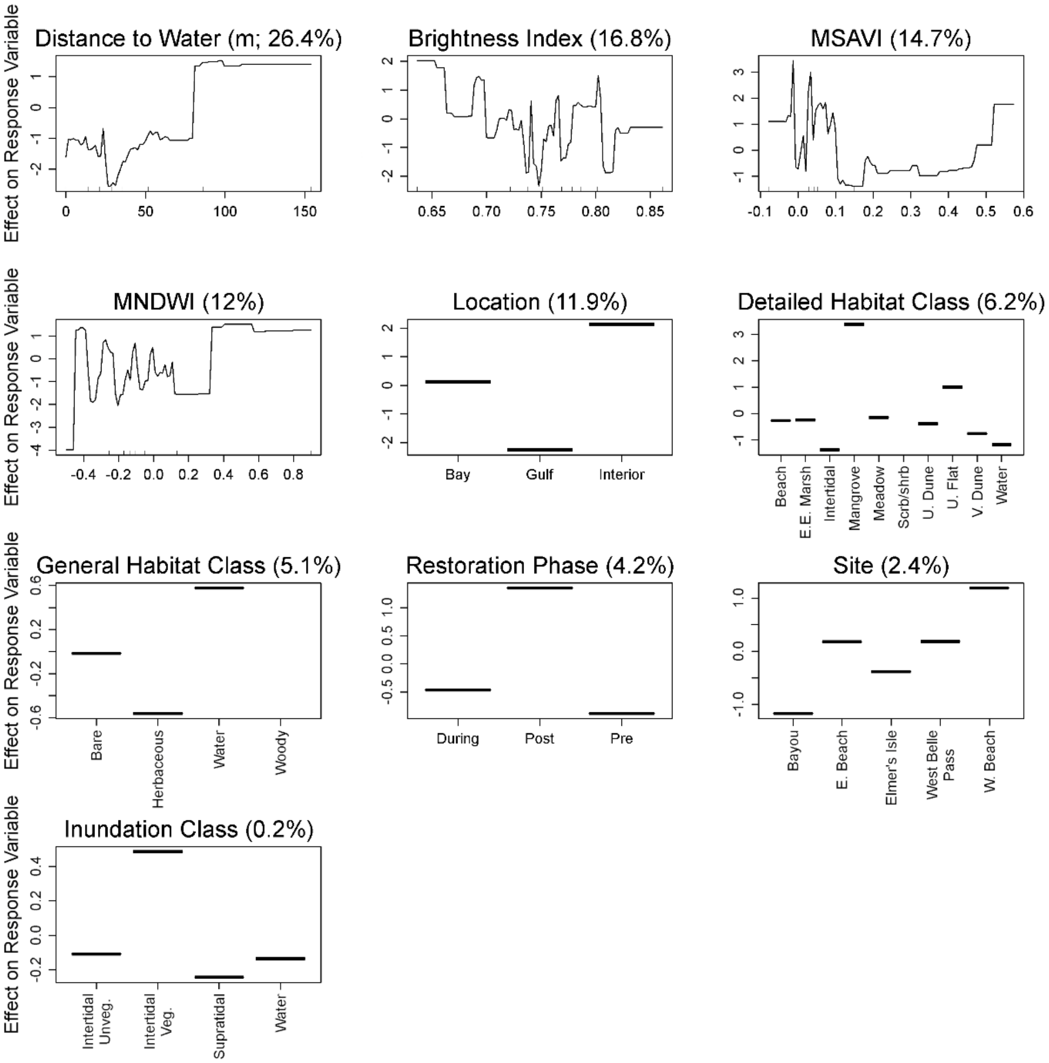
Effect on Response Variable
Distance to Water (m; 26.4%)
Brightness Index (16.8%)
MSAVI (14.7%)
MNDWI (12%)
Location (11.9%)
Bay
Gulf
Interior
Detailed Habitat Class (6.2%)
Beach
E.E. Marsh
Intertidal
Mangrove
Meadow
Scrb/shrb
U. Dune
U. Flat
V. Dune
Water
General Habitat Class (5.1%)
Bare
Herbaceous
Water
Woody
Restoration Phase (4.2%)
During
Post
Pre
Site (2.4%)
Bayou
E. Beach
Elmer's Isle
West Belle Pass
W. Beach
Inundation Class (0.2%)
Intertidal Unveg.
Intertidal Veg.
Supratidal
Water

Figure S7. Partial dependence plots of variables predicting Snowy Plover abundance during the non-breeding seasons on Caminada Headland, Louisiana, USA from a boosted regression tree model. Y-axes are centered to have a zero mean over the data distribution. The relative influence (percent) of each predictor variable is shown in parentheses. Rug plots along the X-axis of each continuous variable plot illustrates the distribution of the data. Abbreviated labels on the X-axis of Detailed Habitat Class include Estuarine Emergent Marsh (E.E. Marsh), Scrub/Shrub (Scrb/Shrb), Unvegetated Dune (U. Dune), Unvegetated Flat (U. Flat), and Vegetated Dune (V. Dune). Abbreviated labels on the X-axis of Inundation Class include Intertidal Unvegetated (Intertidal Unveg.) and Intertidal Vegetated (Intertidal Veg.). Abbreviated labels on the X-axis of Site include East Beach (E. Beach), West Belle Pass (W. Belle Pass), and West Beach (W. Beach). MNDWI indicates Modified Normalized Difference Water Index and MSAVI indicates Modified Soil Adjusted Vegetation Index.

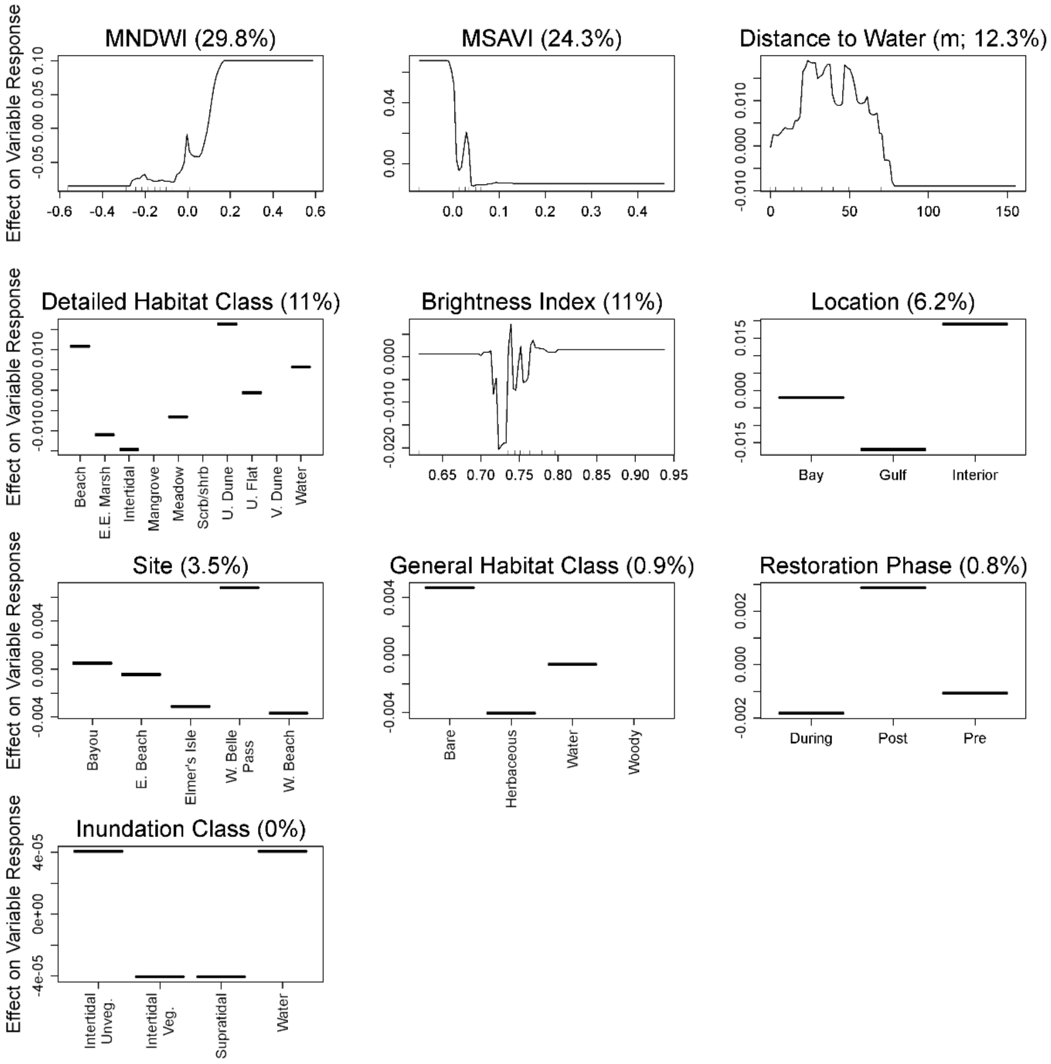

Effect on Variable Response
MNDWI (29.8%)
MSAVI (24.3%)
Distance to Water (m; 12.3%)
Detailed Habitat Class (11%)
Beach
E.E. Marsh
Intertidal
Mangrove
Meadow
Scrb/shrb
U. Dune
U. Flat
V. Dune
Water
Brightness Index (11%)
Location (6.2%)
Bay
Gulf
Interior
Site (3.5%)
Bayou
E. Beach
Elmer's Isle
W. Belle Pass
W. Beach
General Habitat Class (0.9%)
Bare
Herbaceous
Water
Woody
Restoration Phase (0.8%)
During
Post
Pre
Inundation Class (0%)
Intertidal Unveg.
Intertidal Veg.
Supratidal
Water

Figure S8. Partial dependence plots of variables predicting occurrence of Snowy Plover foraging behaviors during the non-breeding seasons on Caminada Headland, Louisiana, USA from a boosted regression tree model. Y-axes are centered to have a zero mean over the data distribution. The relative influence (percent) of each predictor variable is shown in parentheses. Rug plots along the X-axis of each continuous variable plot illustrates the distribution of the data. Abbreviated labels on the X-axis of Detailed Habitat Class include Estuarine Emergent Marsh (E.E. Marsh), Scrub/Shrub (Scrb/Shrb), Unvegetated Dune (U. Dune), Unvegetated Flat (U. Flat), and Vegetated Dune (V. Dune). Abbreviated labels on the X-axis of Inundation Class include Intertidal Unvegetated (Intertidal Unveg.) and Intertidal Vegetated (Intertidal Veg.). Abbreviated labels on the X-axis of Site include East Beach (E. Beach), West Belle Pass (W. Belle Pass), and West Beach (W. Beach). MNDWI indicates Modified Normalized Difference Water Index and MSAVI indicates Modified Soil Adjusted Vegetation Index.

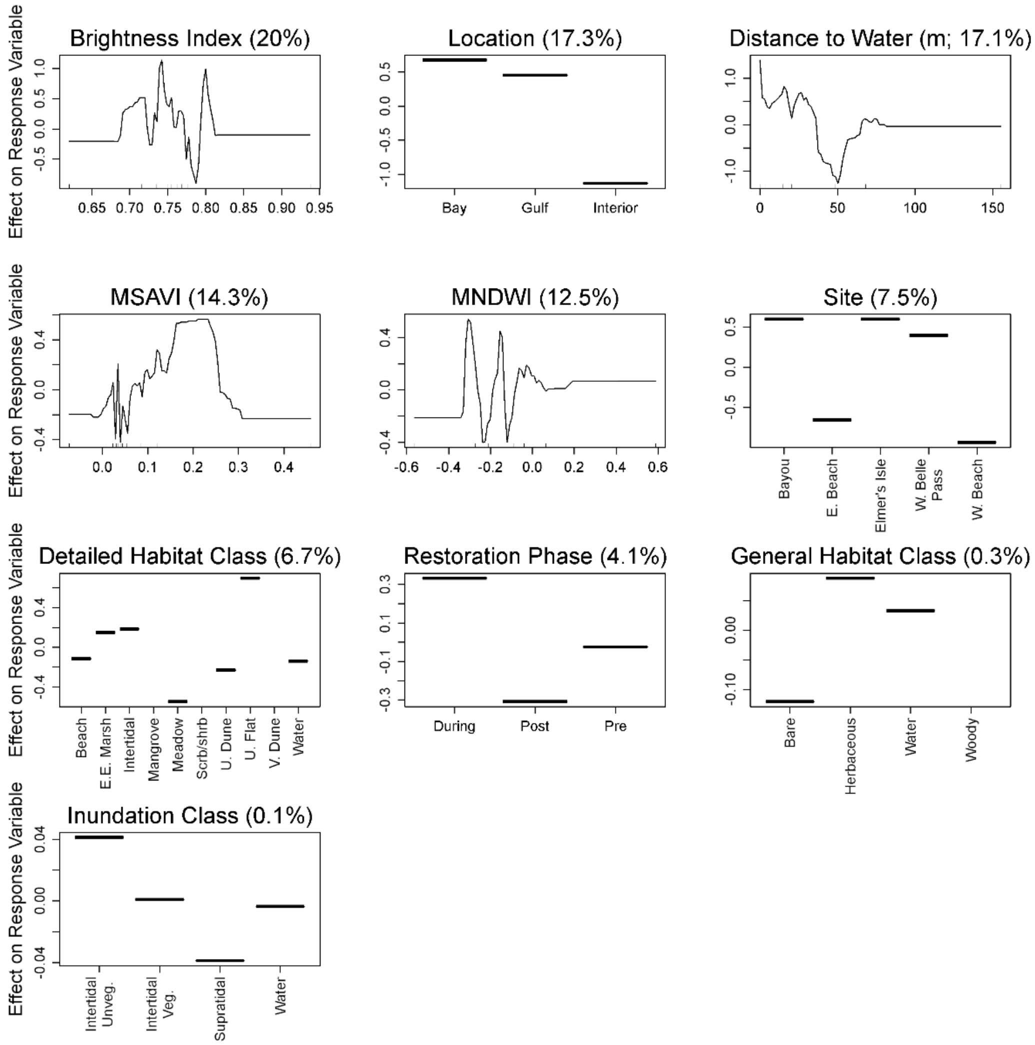
Brightness Index (20%)
Location (17.3%)
Distance to Water (m; 17.1%)
MSAVI (14.3%)
MNDWI (12.5%)
Site (7.5%)
Detailed Habitat Class (6.7%)
Restoration Phase (4.1%)
General Habitat Class (0.3%)
Inundation Class (0.1%)
Effect on Response Variable
Bay
Gulf
Interior
Bayou
E. Beach
Elmer's Isle
W. Belle Pass
W. Beach
Beach
E.E. Marsh
Intertidal
Mangrove
Meadow
Scrb/shrb
U. Dune
U. Flat
V. Dune
Water
During
Post
Pre
Bare
Herbaceous
Water
Woody
Intertidal Unveg.
Intertidal Veg.
Supratidal
Water

Figure S9. Partial dependence plots of variables predicting occurrence of Snowy Plover maintenance behaviors during the non-breeding seasons on Caminada Headland, Louisiana, USA from a boosted regression tree model. Y-axes are centered to have a zero mean over the data distribution. The relative influence (percent) of each predictor variable is shown in parentheses. Rug plots along the X-axis of each continuous variable plot illustrates the distribution of the data. Abbreviated labels on the X-axis of Detailed Habitat Class include Estuarine Emergent Marsh (E.E. Marsh), Scrub/Shrub (Scrb/Shrb), Unvegetated Dune (U. Dune), Unvegetated Flat (U. Flat), and Vegetated Dune (V. Dune). Abbreviated labels on the X-axis of Inundation Class include Intertidal Unvegetated (Intertidal Unveg.) and Intertidal Vegetated (Intertidal Veg.). Abbreviated labels on the X-axis of Site include East Beach (E. Beach), West Belle Pass (W. Belle Pass), and West Beach (W. Beach). MNDWI indicates Modified Normalized Difference Water Index and MSAVI indicates Modified Soil Adjusted Vegetation Index.

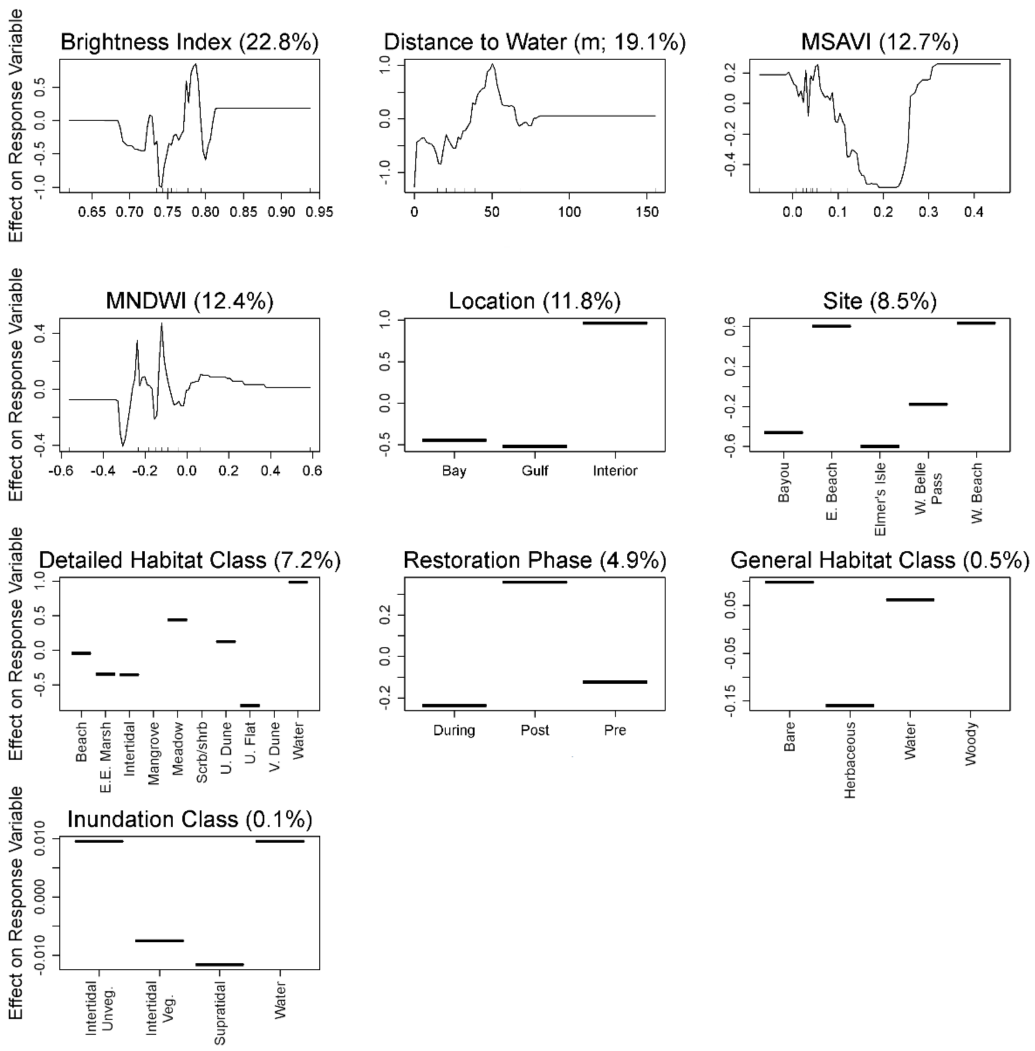

Brightness Index (22.8%)
Distance to Water (m; 19.1%)
MSAVI (12.7%)
MNDWI (12.4%)
Location (11.8%)
Site (8.5%)
Detailed Habitat Class (7.2%)
Restoration Phase (4.9%)
General Habitat Class (0.5%)
Inundation Class (0.1%)
Effect on Response Variable
Bay
Gulf
Interior
Bayou
E. Beach
Elmer's Isle
W. Belle Pass
W. Beach
Beach
E.E. Marsh
Intertidal
Mangrove
Meadow
Scrb/shrb
U. Dune
U. Flat
V. Dune
Water
During
Post
Pre
Bare
Herbaceous
Water
Woody
Intertidal Unveg.
Intertidal Veg.
Supratidal
Water

Figure S10. Partial dependence plots of variables predicting Wilson's Plover abundance during the breeding season on Caminada Headland, Louisiana, USA from a boosted regression tree model. Y-axes are centered to have a zero mean over the data distribution. The relative influence (percent) of each predictor variable is shown in parentheses. Rug plots along the X-axis of each continuous variable plot illustrates the distribution of the data. Abbreviated labels on the X-axis of Detailed Habitat Class include Estuarine Emergent Marsh (E.E. Marsh), Scrub/Shrub (Scrb/Shrb), Unvegetated Dune (U. Dune), Unvegetated Flat (U. Flat), and Vegetated Dune (V. Dune). Abbreviated labels on the X-axis of Inundation Class include Intertidal Unvegetated (Intertidal Unveg.) and Intertidal Vegetated (Intertidal Veg.). Abbreviated labels on the X-axis of Site include East Beach (E. Beach), West Belle Pass (W. Belle Pass), and West Beach (W. Beach). MNDWI indicates Modified Normalized Difference Water Index and MSAVI indicates Modified Soil Adjusted Vegetation Index.

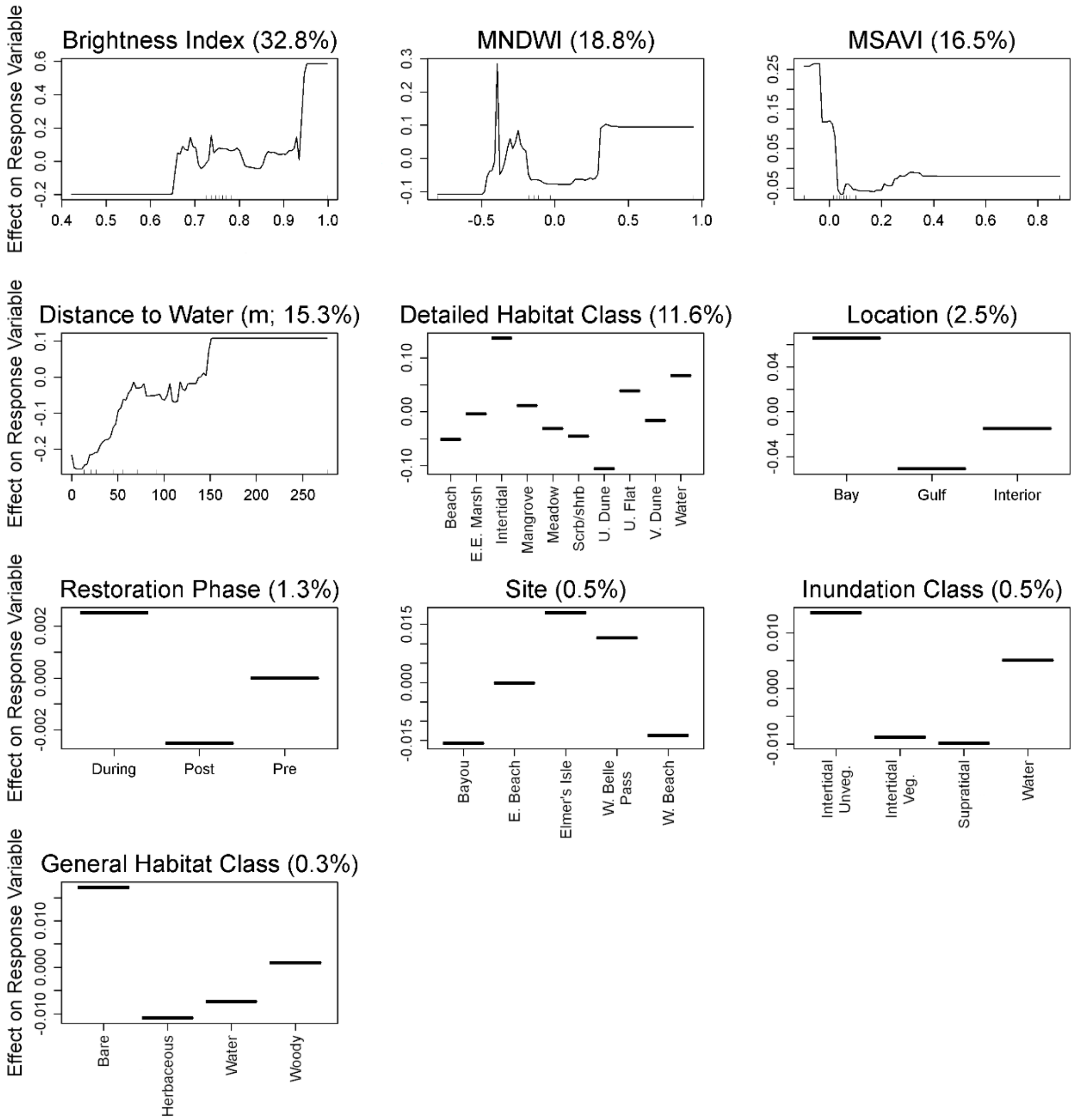

Effect on Response Variable
Brightness Index (32.8%)
MNDWI (18.8%)
MSAVI (16.5%)
Distance to Water (m; 15.3%)
Detailed Habitat Class (11.6%)
Beach
E.E. Marsh
Intertidal
Mangrove
Meadow
Scrb/shrb
U. Dune
U. Flat
V. Dune
Water
Location (2.5%)
Bay
Gulf
Interior
Restoration Phase (1.3%)
During
Post
Pre
Site (0.5%)
Bayou
E. Beach
Elmer's Isle
W. Belle Pass
W. Beach
Inundation Class (0.5%)
Intertidal Unveg.
Intertidal Veg.
Supratidal
Water
General Habitat Class (0.3%)
Bare
Herbaceous
Water
Woody

Figure S11. Partial dependence plots of variables predicting Wilson's Plover abundance during the non-breeding seasons on Caminada Headland, Louisiana, USA from a boosted regression tree model. Y-axes are centered to have a zero mean over the data distribution. The relative influence (percent) of each predictor variable is shown in parentheses. Rug plots along the X-axis of each continuous variable plot illustrates the distribution of the data. Abbreviated labels on the X-axis of Detailed Habitat Class include Estuarine Emergent Marsh (E.E. Marsh), Scrub/Shrub (Scrb/Shrb), Unvegetated Dune (U. Dune), Unvegetated Flat (U. Flat), and Vegetated Dune (V. Dune). Abbreviated labels on the X-axis of Inundation Class include Intertidal Unvegetated (Intertidal Unveg.) and Intertidal Vegetated (Intertidal Veg.). Abbreviated labels on the X-axis of Site include East Beach (E. Beach), West Belle Pass (W. Belle Pass), and West Beach (W. Beach). MNDWI indicates Modified Normalized Difference Water Index and MSAVI indicates Modified Soil Adjusted Vegetation Index.

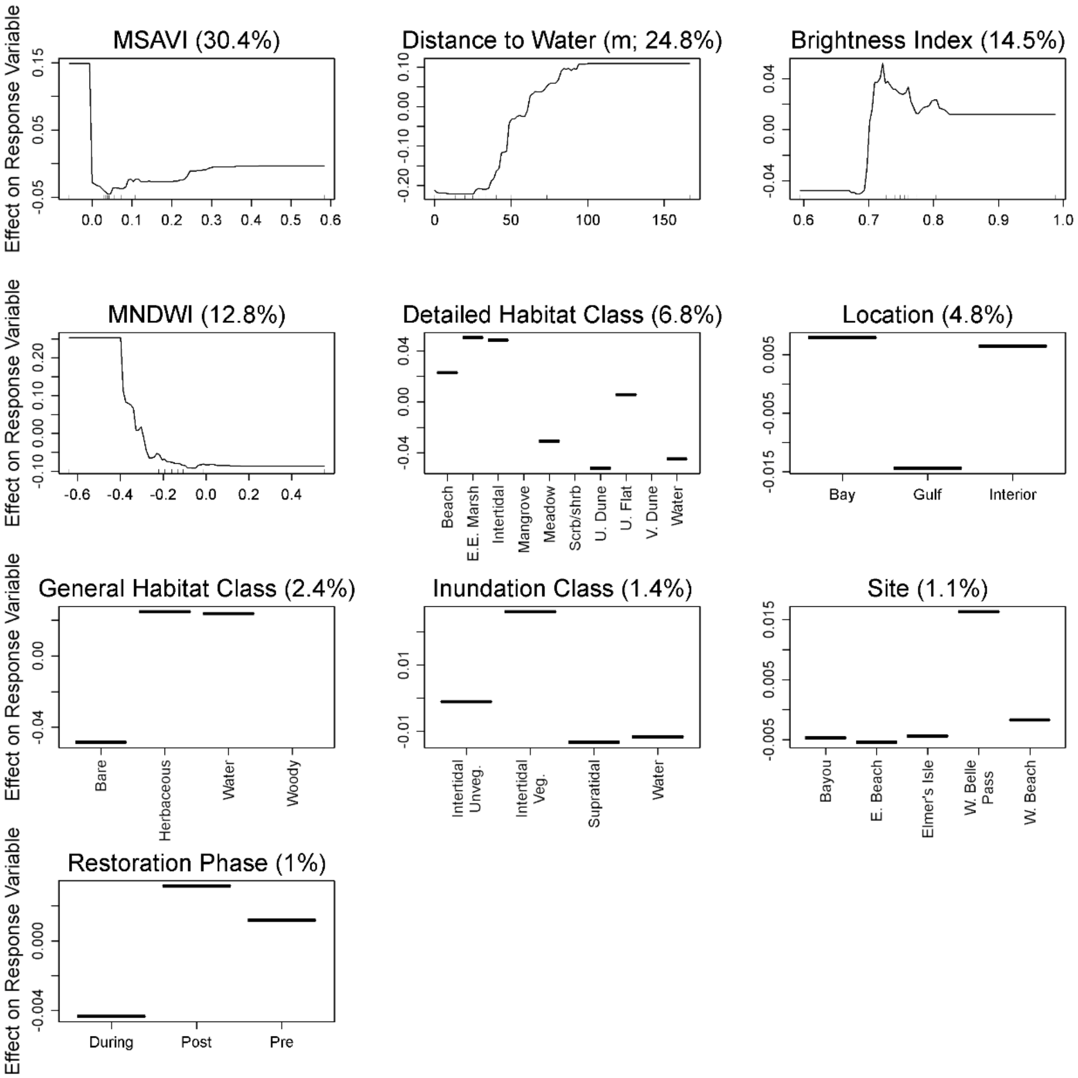

MSAVI (30.4%)
Distance to Water (m; 24.8%)
Brightness Index (14.5%)
MNDWI (12.8%)
Detailed Habitat Class (6.8%)
Location (4.8%)
General Habitat Class (2.4%)
Inundation Class (1.4%)
Site (1.1%)
Restoration Phase (1%)
Effect on Response Variable
Beach
E.E. Marsh
Intertidal
Mangrove
Meadow
Scrb/shrb
U. Dune
U. Flat
V. Dune
Water
Bay
Gulf
Interior
Bare
Herbaceous
Woody
Intertidal Unveg.
Intertidal Veg.
Supratidal
Bayou
E. Beach
Elmer's Isle
W. Belle Pass
W. Beach
During
Post
Pre

Figure S12. Partial dependence plots of variables predicting occurrence of Wilson's Plover breeding behaviors on Caminada Headland, Louisiana, USA from a boosted regression tree model. Y-axes are centered to have a zero mean over the data distribution. The relative influence (percent) of each predictor variable is shown in parentheses. Rug plots along the X-axis of each continuous variable plot illustrates the distribution of the data. Abbreviated labels on the X-axis of Detailed Habitat Class include Estuarine Emergent Marsh (E.E. Marsh), Scrub/Shrub (Scrb/Shrb), Unvegetated Dune (U. Dune), Unvegetated Flat (U. Flat), and Vegetated Dune (V. Dune). Abbreviated labels on the X-axis of Inundation Class include Intertidal Unvegetated (Intertidal Unveg.) and Intertidal Vegetated (Intertidal Veg.). Abbreviated labels on the X-axis of Site include East Beach (E. Beach), West Belle Pass (W. Belle Pass), and West Beach (W. Beach). MNDWI indicates Modified Normalized Difference Water Index and MSAVI indicates Modified Soil Adjusted Vegetation Index.

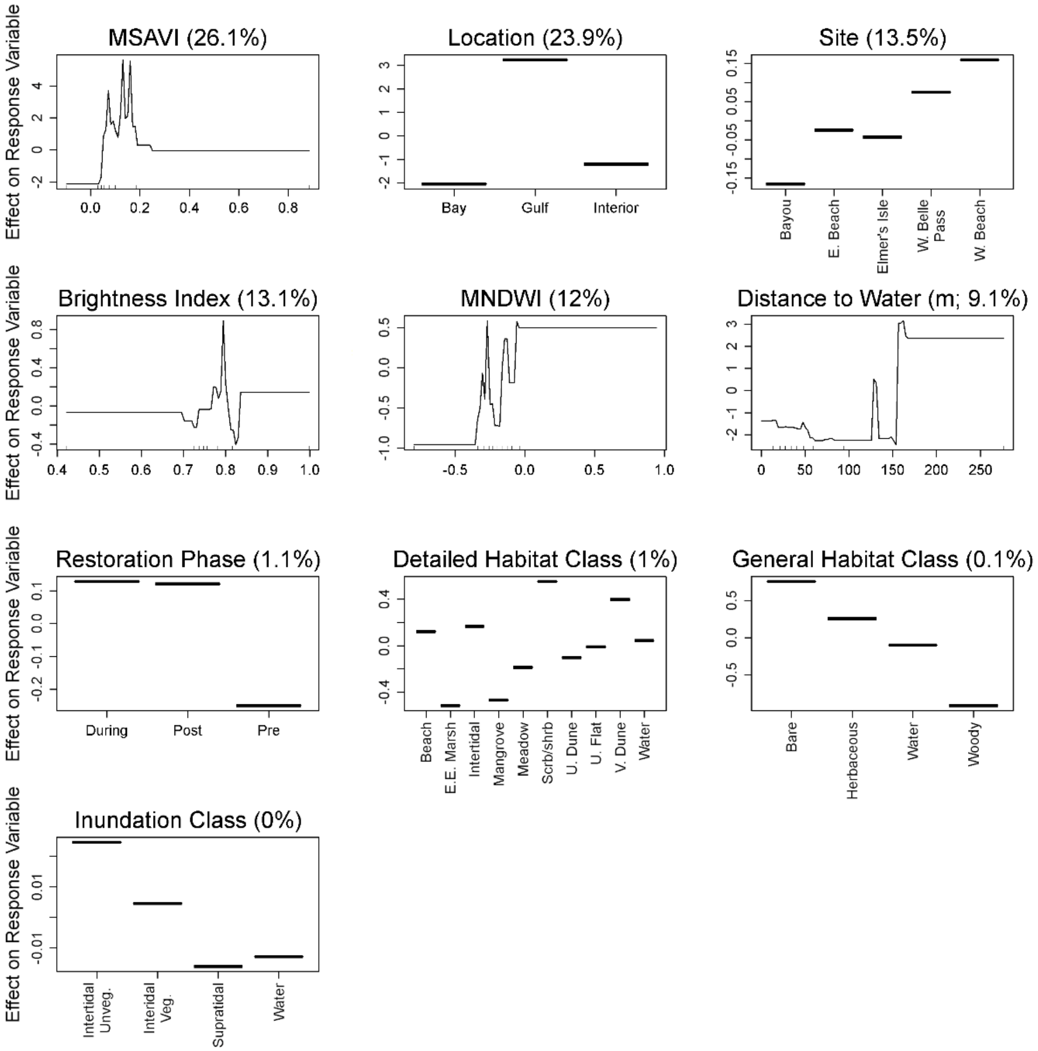
MSAVI (26.1%)
Location (23.9%)
Site (13.5%)
Brightness Index (13.1%)
MNDWI (12%)
Distance to Water (m; 9.1%)
Restoration Phase (1.1%)
Detailed Habitat Class (1%)
General Habitat Class (0.1%)
Inundation Class (0%)
Effect on Response Variable
Bay
Gulf
Interior
Bayou
E. Beach
Elmer's Isle
W. Belle Pass
W. Beach
During
Post
Pre
Beach
E.E. Marsh
Intertidal
Mangrove
Meadow
Scrb/shrb
U. Dune
U. Flat
V. Dune
Water
Bare
Herbaceous
Water
Woody
Intertidal Unveg.
Interidal Veg.
Supratidal
Water

Figure S13. Partial dependence plots of variables predicting occurrence of Wilson's Plover foraging behaviors on Caminada Headland, Louisiana, USA from a boosted regression tree model. Y-axes are centered to have a zero mean over the data distribution. The relative influence (percent) of each predictor variable is shown in parentheses. Rug plots along the X-axis of each continuous variable plot illustrates the distribution of the data. Abbreviated labels on the X-axis of Detailed Habitat Class include Estuarine Emergent Marsh (E.E. Marsh), Scrub/Shrub (Scrb/Shrb), Unvegetated Dune (U. Dune), Unvegetated Flat (U. Flat), and Vegetated Dune (V. Dune). Abbreviated labels on the X-axis of Inundation Class include Intertidal Unvegetated (Intertidal Unveg.) and Intertidal Vegetated (Intertidal Veg.). Abbreviated labels on the X-axis of Site include East Beach (E. Beach), West Belle Pass (W. Belle Pass), and West Beach (W. Beach). MNDWI indicates Modified Normalized Difference Water Index and MSAVI indicates Modified Soil Adjusted Vegetation Index.

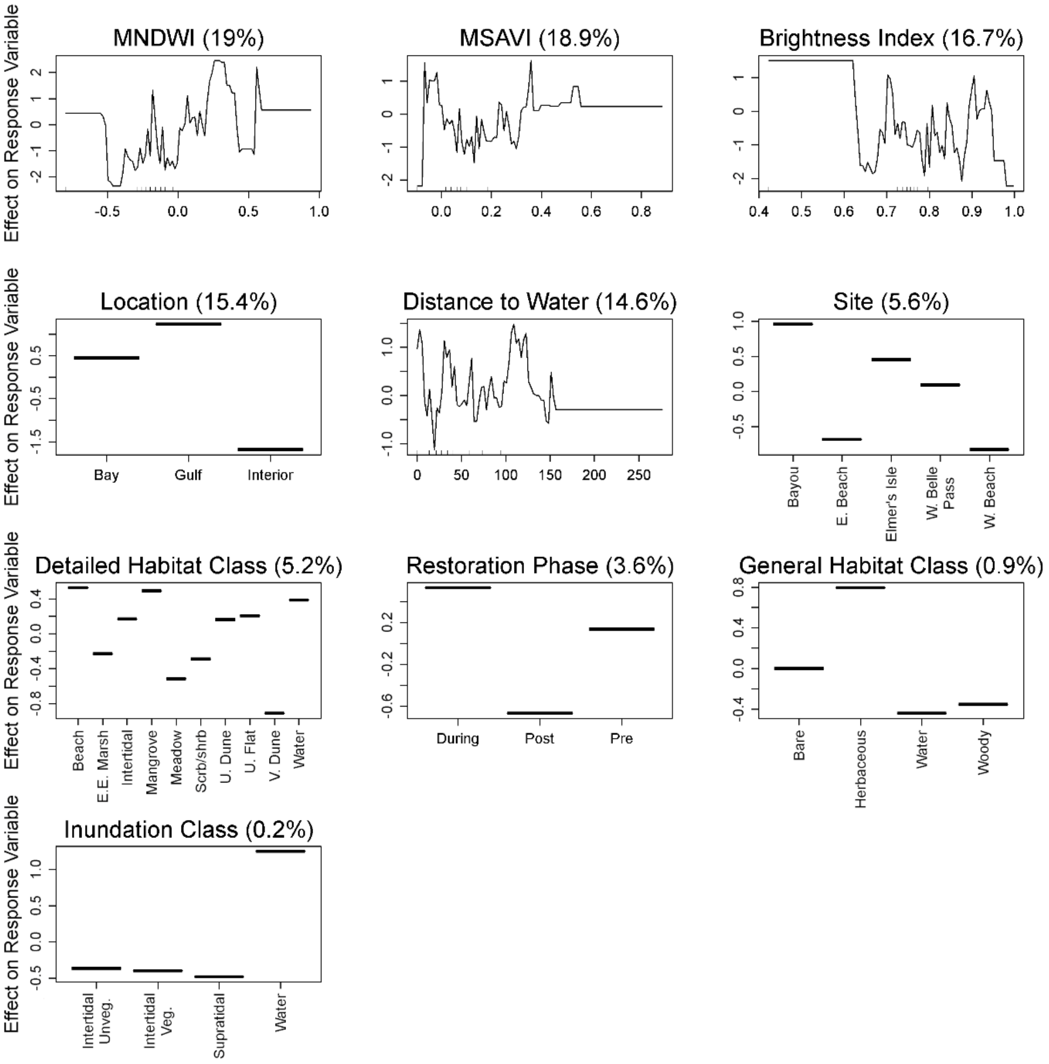

Effect on Response Variable
MNDWI (19%)
MSAVI (18.9%)
Brightness Index (16.7%)
Location (15.4%)
Bay
Gulf
Interior
Distance to Water (14.6%)
Site (5.6%)
Bayou
E. Beach
Elmer's Isle
W. Belle Pass
W. Beach
Detailed Habitat Class (5.2%)
Beach
E.E. Marsh
Intertidal
Mangrove
Meadow
Scrb/shrb
U. Dune
U. Flat
V. Dune
Water
Restoration Phase (3.6%)
During
Post
Pre
General Habitat Class (0.9%)
Bare
Herbaceous
Water
Woody
Inundation Class (0.2%)
Intertidal Unveg.
Intertidal Veg.
Supratidal
Water

Figure S14. Partial dependence plots of variables predicting occurrence of Wilson's Plover maintenance behaviors on Caminada Headland, Louisiana, USA from a boosted regression tree model. Y-axes are centered to have a zero mean over the data distribution. The relative influence (percent) of each predictor variable is shown in parentheses. Rug plots along the X-axis of each continuous variable plot illustrates the distribution of the data. Abbreviated labels on the X-axis of Detailed Habitat Class include Estuarine Emergent Marsh (E.E. Marsh), Scrub/Shrub (Scrb/Shrb), Unvegetated Dune (U. Dune), Unvegetated Flat (U. Flat), and Vegetated Dune (V. Dune). Abbreviated labels on the X-axis of Inundation Class include Intertidal Unvegetated (Intertidal Unveg.) and Intertidal Vegetated (Intertidal Veg.). Abbreviated labels on the X-axis of Site include East Beach (E. Beach), West Belle Pass (W. Belle Pass), and West Beach (W. Beach). MNDWI indicates Modified Normalized Difference Water Index and MSAVI indicates Modified Soil Adjusted Vegetation Index.

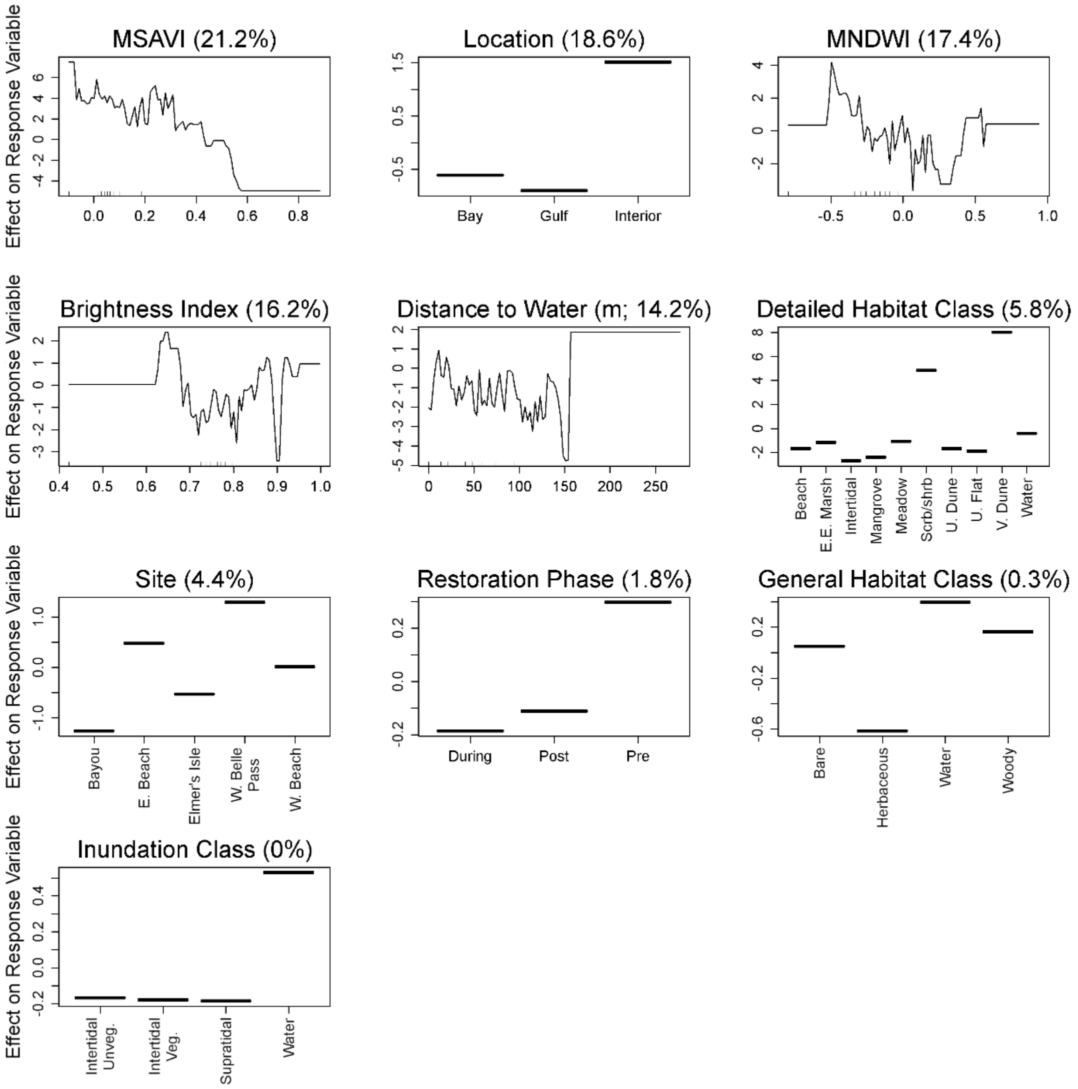

Effect on Response Variable
MSAVI (21.2%)
Location (18.6%)
Bay
Gulf
Interior
MNDWI (17.4%)
Brightness Index (16.2%)
Distance to Water (m; 14.2%)
Detailed Habitat Class (5.8%)
Beach
E.E. Marsh
Intertidal
Mangrove
Meadow
Scrb/shrb
U. Dune
U. Flat
V. Dune
Water
Site (4.4%)
Bayou
E. Beach
Elmer's Isle
W. Belle Pass
W. Beach
Restoration Phase (1.8%)
During
Post
Pre
General Habitat Class (0.3%)
Bare
Herbaceous
Water
Woody
Inundation Class (0%)
Intertidal Unveg.
Intertidal Veg.
Supratidal
Water

Figure S15. Partial dependence plots of variables predicting American Oystercatcher abundance during the breeding season on Whiskey Island, Louisiana, USA from a boosted regression tree model. Y-axes are centered to have a zero mean over the data distribution. The relative influence (percent) of each predictor variable is shown in parentheses. Rug plots along the X-axis of each continuous variable plot illustrates the distribution of the data. Abbreviated labels on the X-axis of Detailed Habitat Class include Estuarine Emergent Marsh (E.E. Marsh), Scrub/Shrub (Scrb/Shrb), Shoreline Protection (S. Protect), Unvegetated Dune (U. Dune), and Vegetated Dune (V. Dune). Abbreviated labels on the X-axis of Inundation Class include Intertidal Unvegetated (Intertidal Unveg.) and Intertidal Vegetated (Intertidal Veg.). MNDWI indicates Modified Normalized Difference Water Index and MSAVI indicates Modified Soil Adjusted Vegetation Index.

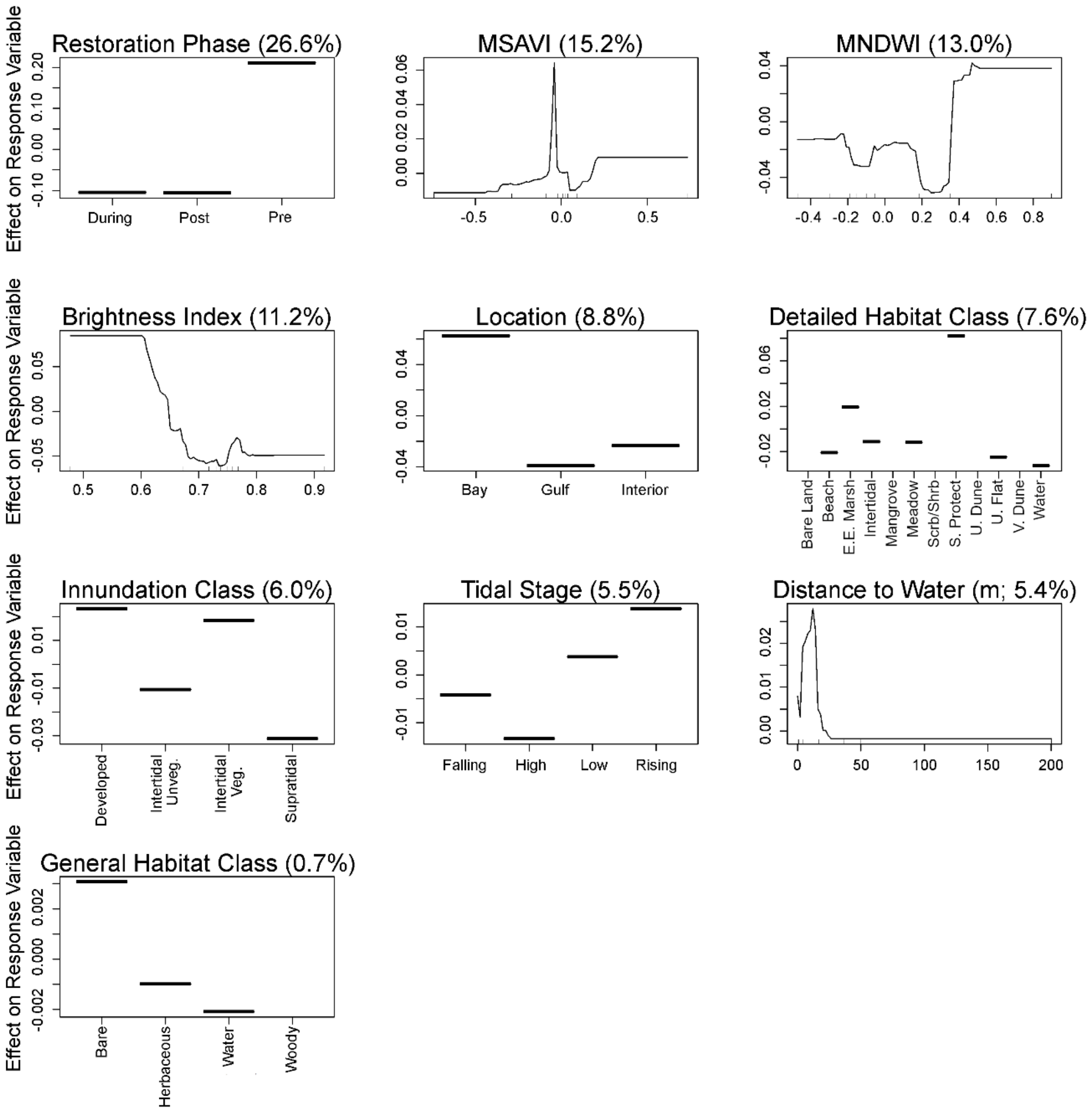
Effect on Response Variable
Restoration Phase (26.6%)
-0.10 0.00 0.10 0.20
During
Post
Pre
MSAVI (15.2%)
0.00 0.02 0.04 0.06
-0.5 0.0 0.5
MNDWI (13.0%)
-0.04 0.00 0.04
-0.4 -0.2 0.0 0.2 0.4 0.6 0.8
Brightness Index (11.2%)
-0.05 0.00 0.05
0.5 0.6 0.7 0.8 0.9
Location (8.8%)
-0.04 0.00 0.04
Bay
Gulf
Interior
Detailed Habitat Class (7.6%)
-0.02 0.02 0.06
Bare Land
Beach
E.E. Marsh
Intertidal
Mangrove
Meadow
Scrb/Shrb
S. Protect
U. Dune
U. Flat
V. Dune
Water
Innundation Class (6.0%)
-0.03 -0.01 0.01
Developed
Intertidal Unveg.
Intertidal Veg.
Supratidal
Tidal Stage (5.5%)
-0.01 0.00 0.01
Falling
High
Low
Rising
Distance to Water (m; 5.4%)
0.00 0.01 0.02
0 50 100 150 200
General Habitat Class (0.7%)
-0.002 0.000 0.002
Bare
Herbaceous
Water
Woody

Figure S16. Partial dependence plots of variables predicting American Oystercatcher abundance during non-breeding seasons on Whiskey Island, Louisiana, USA from a boosted regression tree model. Y-axes are centered to have a zero mean over the data distribution. The relative influence (percent) of each predictor variable is shown in parentheses. Rug plots along the X-axis of each continuous variable plot illustrates the distribution of the data. Abbreviated labels on the X-axis of Detailed Habitat Class include Estuarine Emergent Marsh (E.E. Marsh), Scrub/Shrub (Scrb/Shrb), Shoreline Protection (S. Protect), Unvegetated Dune (U. Dune), and Vegetated Dune (V. Dune). Abbreviated labels on the X-axis of Inundation Class include Intertidal Unvegetated (Intertidal Unveg.) and Intertidal Vegetated (Intertidal Veg.). MNDWI indicates Modified Normalized Difference Water Index and MSAVI indicates Modified Soil Adjusted Vegetation Index.

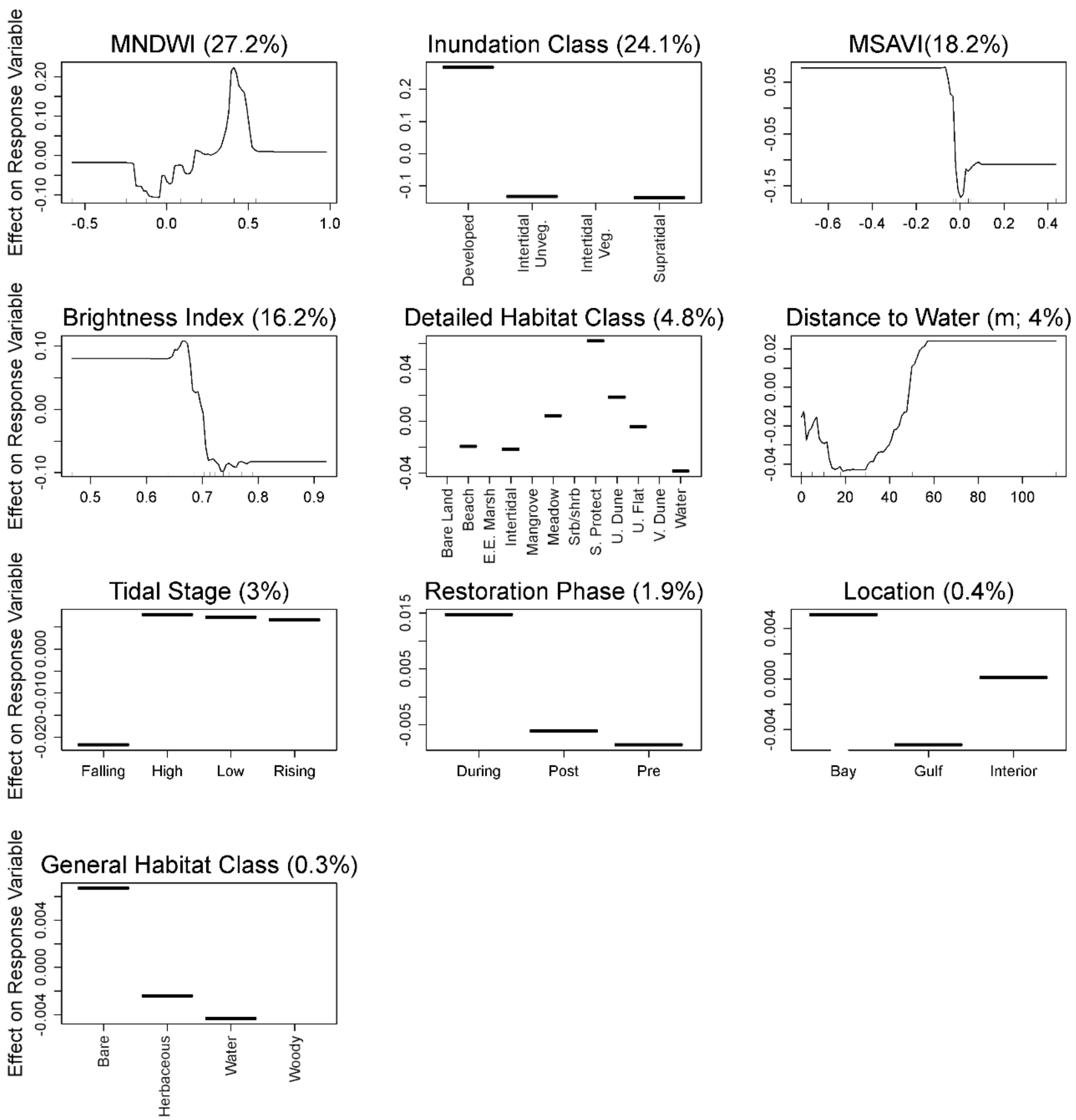

Effect on Response Variable
MNDWI (27.2%)
Inundation Class (24.1%)
Developed
Intertidal Unveg.
Intertidal Veg.
Supratidal
MSAVI(18.2%)
Brightness Index (16.2%)
Detailed Habitat Class (4.8%)
Bare Land
Beach
E.E. Marsh
Intertidal
Mangrove
Meadow
Srb/shrb
S. Protect
U. Dune
U. Flat
V. Dune
Water
Distance to Water (m; 4%)
Tidal Stage (3%)
Falling
High
Low
Rising
Restoration Phase (1.9%)
During
Post
Pre
Location (0.4%)
Bay
Gulf
Interior
General Habitat Class (0.3%)
Bare
Herbaceous
Water
Woody

Figure S17. Partial dependence plots of variables predicting the occurrence of American Oystercatcher breeding behaviors on Whiskey Island, Louisiana, USA from a boosted regression tree model. Y-axes are centered to have a zero mean over the data distribution. The relative influence (percent) of each predictor variable is shown in parentheses. Rug plots along the X-axis of each continuous variable plot illustrates the distribution of the data. Abbreviated labels on the X-axis of Detailed Habitat Class include Estuarine Emergent Marsh (E.E. Marsh), Scrub/Shrub (Scrb/Shrb), Shoreline Protection (S. Protect), Unvegetated Dune (U. Dune), and Vegetated Dune (V. Dune). Abbreviated labels on the X-axis of Inundation Class include Intertidal Unvegetated (Intertidal Unveg.) and Intertidal Vegetated (Intertidal Veg.). MNDWI indicates Modified Normalized Difference Water Index and MSAVI indicates Modified Soil Adjusted Vegetation Index.

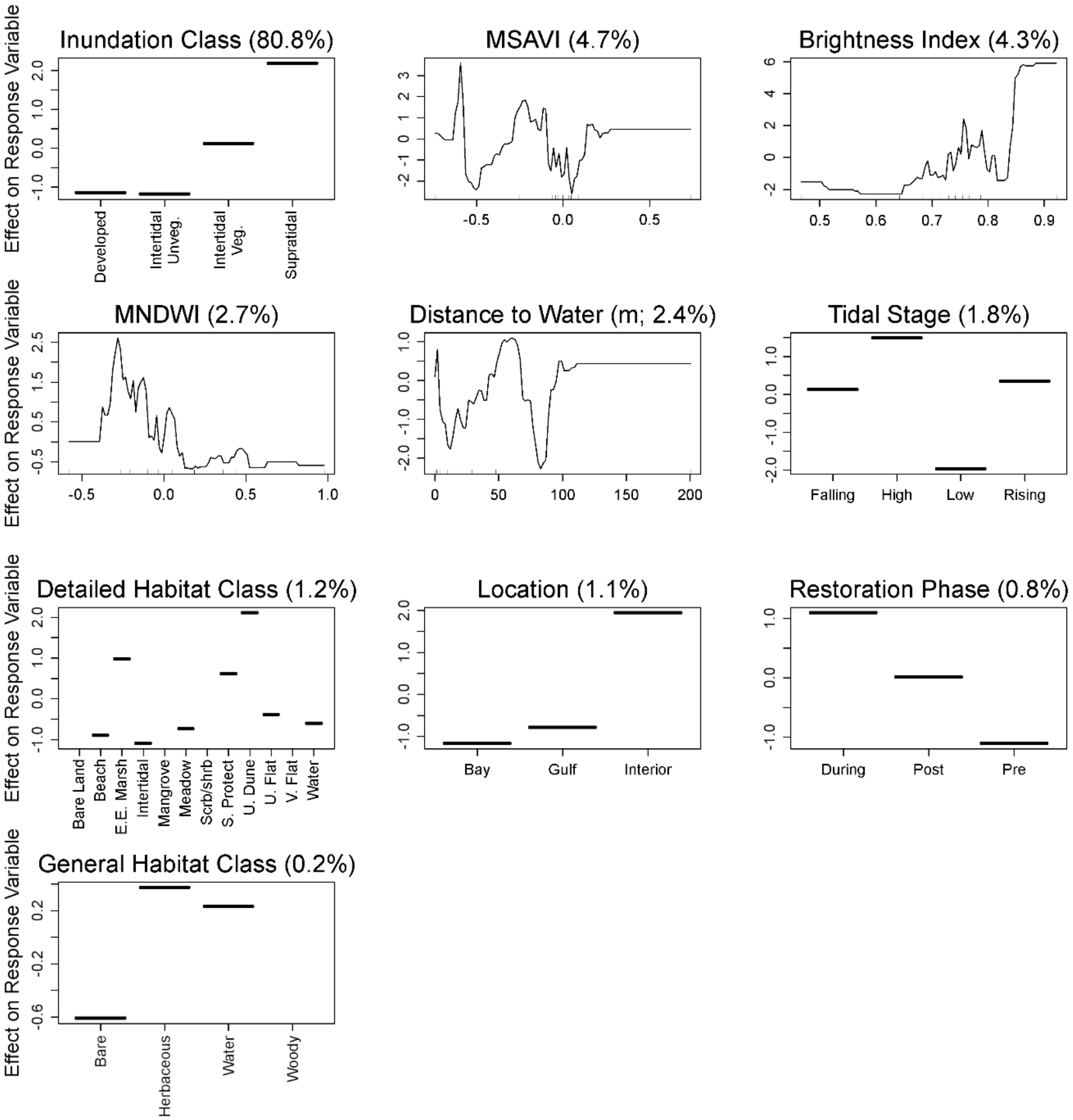
Effect on Response Variable
Inundation Class (80.8%)
Developed
Intertidal Unveg.
Intertidal Veg.
Supratidal
MSAVI (4.7%)
Brightness Index (4.3%)
MNDWI (2.7%)
Distance to Water (m; 2.4%)
Tidal Stage (1.8%)
Falling
High
Low
Rising
Detailed Habitat Class (1.2%)
Bare Land
Beach
E.E. Marsh
Intertidal
Mangrove
Meadow
Scrb/shrb
S. Protect
U. Dune
U. Flat
V. Flat
Water
Location (1.1%)
Bay
Gulf
Interior
Restoration Phase (0.8%)
During
Post
Pre
General Habitat Class (0.2%)
Bare
Herbaceous
Water
Woody

Figure S18. Partial dependence plots of variables predicting the occurrence of American Oystercatcher foraging behaviors on Whiskey Island, Louisiana, USA from a boosted regression tree model. Y-axes are centered to have a zero mean over the data distribution. The relative influence (percent) of each predictor variable is shown in parentheses. Rug plots along the X-axis of each continuous variable plot illustrates the distribution of the data. Abbreviated labels on the X-axis of Detailed Habitat Class include Estuarine Emergent Marsh (E.E. Marsh), Scrub/Shrub (Scrb/Shrb), Shoreline Protection (S. Protect), Unvegetated Dune (U. Dune), and Vegetated Dune (V. Dune). Abbreviated labels on the X-axis of Inundation Class include Intertidal Unvegetated (Intertidal Unveg.) and Intertidal Vegetated (Intertidal Veg.). MNDWI indicates Modified Normalized Difference Water Index and MSAVI indicates Modified Soil Adjusted Vegetation Index.

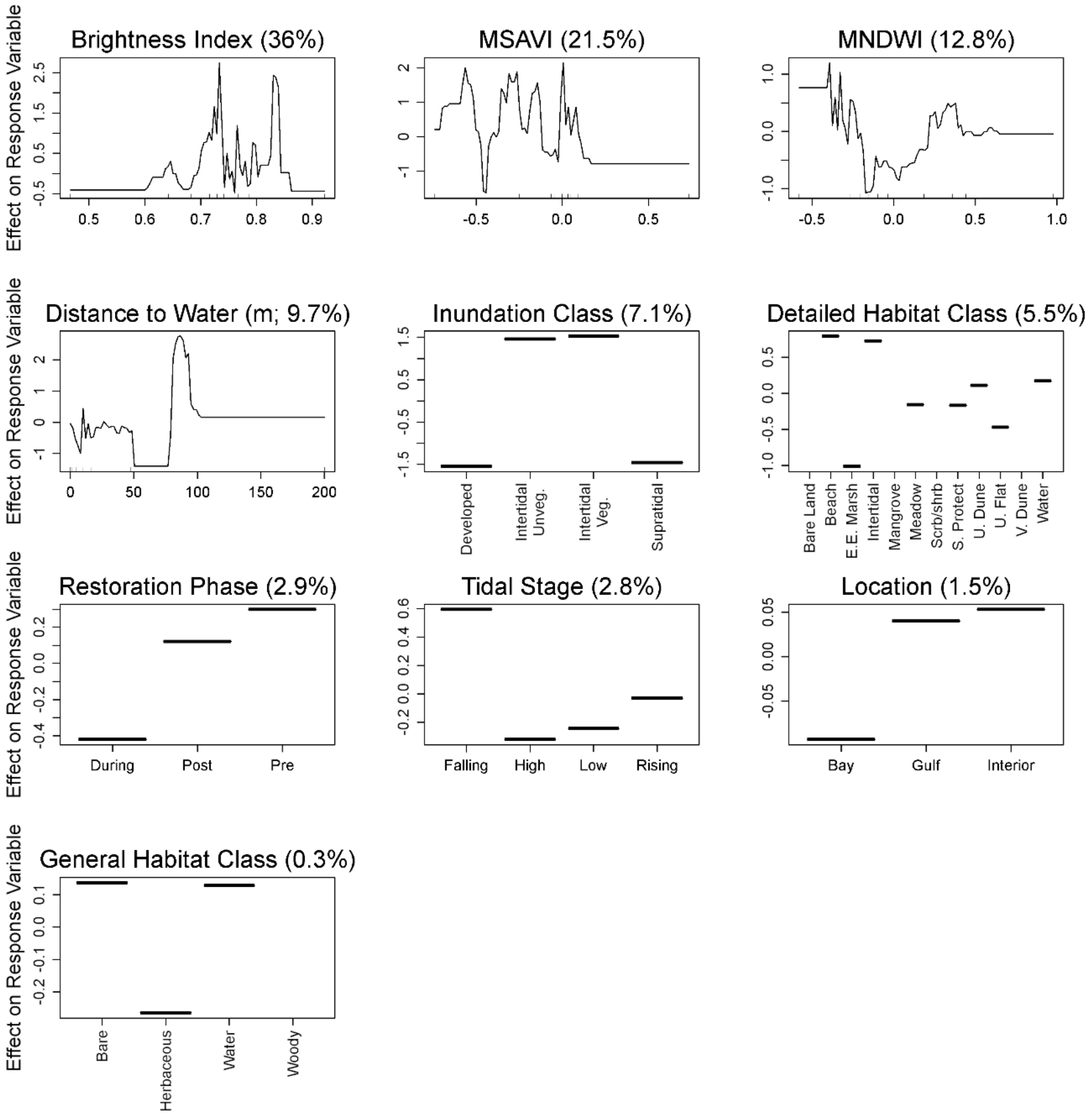
Effect on Response Variable
Brightness Index (36%)
MSAVI (21.5%)
MNDWI (12.8%)
Distance to Water (m; 9.7%)
Inundation Class (7.1%)
Developed
Intertidal Unveg.
Intertidal Veg.
Supratidal
Detailed Habitat Class (5.5%)
Bare Land
Beach
E.E. Marsh
Intertidal
Mangrove
Meadow
Scrb/shrb
S. Protect
U. Dune
U. Flat
V. Dune
Water
Restoration Phase (2.9%)
During
Post
Pre
Tidal Stage (2.8%)
Falling
High
Low
Rising
Location (1.5%)
Bay
Gulf
Interior
General Habitat Class (0.3%)
Bare
Herbaceous
Water
Woody

Figure S19. Partial dependence plots of variables predicting the occurrence of American Oystercatcher maintenance behaviors on Whiskey Island, Louisiana, USA from a boosted regression tree model. Y-axes are centered to have a zero mean over the data distribution. The relative influence (percent) of each predictor variable is shown in parentheses. Rug plots along the X-axis of each continuous variable plot illustrates the distribution of the data. Abbreviated labels on the X-axis of Detailed Habitat Class include Estuarine Emergent Marsh (E.E. Marsh), Scrub/Shrub (Scrb/Shrb), Shoreline Protection (S. Protect), Unvegetated Dune (U. Dune), and Vegetated Dune (V. Dune). Abbreviated labels on the X-axis of Inundation Class include Intertidal Unvegetated (Intertidal Unveg.) and Intertidal Vegetated (Intertidal Veg.). MNDWI indicates Modified Normalized Difference Water Index and MSAVI indicates Modified Soil Adjusted Vegetation Index.

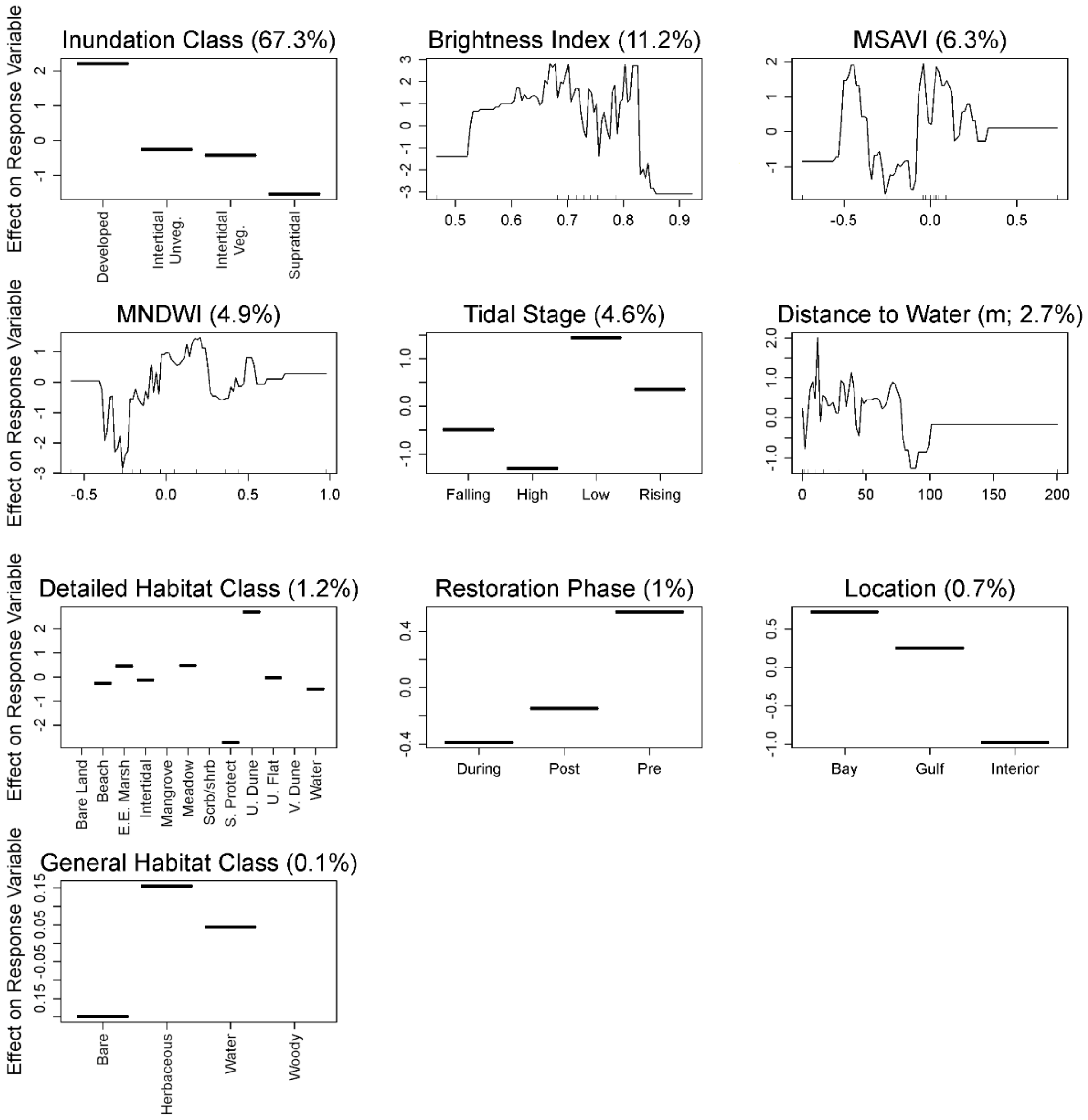

Effect on Response Variable
Inundation Class (67.3%)
Developed
Intertidal Unveg.
Intertidal Veg.
Supratidal
Brightness Index (11.2%)
MSAVI (6.3%)
MNDWI (4.9%)
Tidal Stage (4.6%)
Falling
High
Low
Rising
Distance to Water (m; 2.7%)
Detailed Habitat Class (1.2%)
Bare Land
Beach
E.E. Marsh
Intertidal
Mangrove
Meadow
Scrb/shrb
S. Protect
U. Dune
U. Flat
V. Dune
Water
Restoration Phase (1%)
During
Post
Pre
Location (0.7%)
Bay
Gulf
Interior
General Habitat Class (0.1%)
Bare
Herbaceous
Water
Woody

Figure S20. Partial dependence plots of variables predicting Piping Plover abundance during the non-breeding seasons on Whiskey Island, Louisiana, USA from a boosted regression tree model. Y-axes are centered to have a zero mean over the data distribution. The relative influence (percent) of each predictor variable is shown in parentheses. Rug plots along the X-axis of each continuous variable plot illustrates the distribution of the data. Abbreviated labels on the X-axis of Detailed Habitat Class include Estuarine Emergent Marsh (E.E. Marsh), Scrub/Shrub (Scrb/Shrb), Shoreline Protection (S. Protect), Unvegetated Dune (U. Dune), and Vegetated Dune (V. Dune). Abbreviated labels on the X-axis of Inundation Class include Intertidal Unvegetated (Intertidal Unveg.) and Intertidal Vegetated (Intertidal Veg.). MNDWI indicates Modified Normalized Difference Water Index and MSAVI indicates Modified Soil Adjusted Vegetation Index.

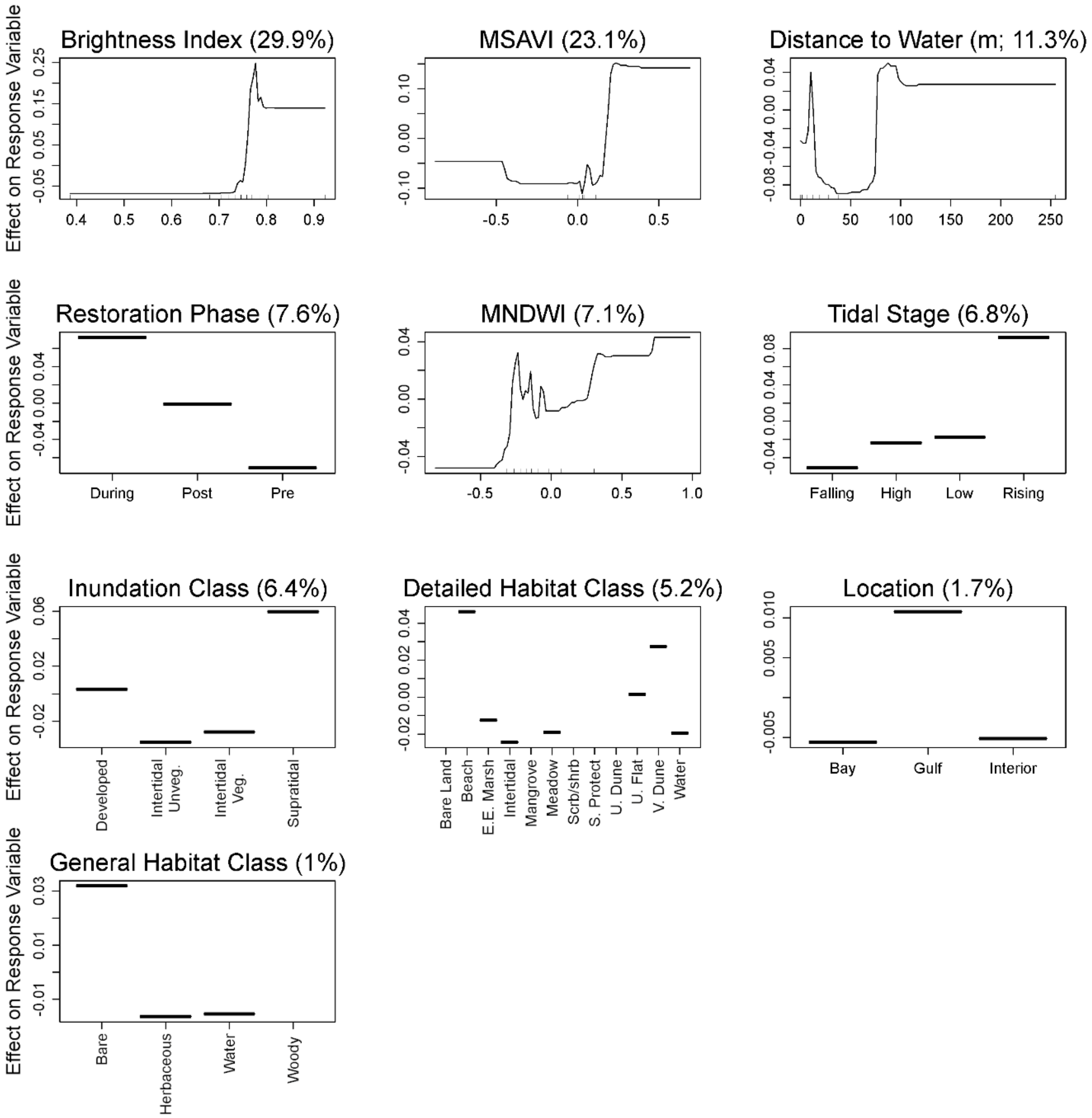
Brightness Index (29.9%)
MSAVI (23.1%)
Distance to Water (m; 11.3%)
Restoration Phase (7.6%)
MNDWI (7.1%)
Tidal Stage (6.8%)
Inundation Class (6.4%)
Detailed Habitat Class (5.2%)
Location (1.7%)
General Habitat Class (1%)
Effect on Response Variable
During
Post
Pre
Falling
High
Low
Rising
Developed
Intertidal Unveg.
Intertidal Veg.
Supratidal
Bare Land
Beach
E.E. Marsh
Intertidal
Mangrove
Meadow
Scrb/shrb
S. Protect
U. Dune
U. Flat
V. Dune
Water
Bay
Gulf
Interior
Bare
Herbaceous
Water
Woody

Figure S21. Partial dependence plots of variables predicting occurrences of Piping Plover foraging behaviors during the non-breeding seasons on Whiskey Island, Louisiana, USA from a boosted regression tree model. Y-axes are centered to have a zero mean over the data distribution. The relative influence (percent) of each predictor variable is shown in parentheses. Rug plots along the X-axis of each continuous variable plot illustrates the distribution of the data. Abbreviated labels on the X-axis of Detailed Habitat Class include Estuarine Emergent Marsh (E.E. Marsh), Scrub/Shrub (Scrb/Shrb), Shoreline Protection (S. Protect), Unvegetated Dune (U. Dune), and Vegetated Dune (V. Dune). Abbreviated labels on the X-axis of Inundation Class include Intertidal Unvegetated (Intertidal Unveg.) and Intertidal Vegetated (Intertidal Veg.). MNDWI indicates Modified Normalized Difference Water Index and MSAVI indicates Modified Soil Adjusted Vegetation Index.

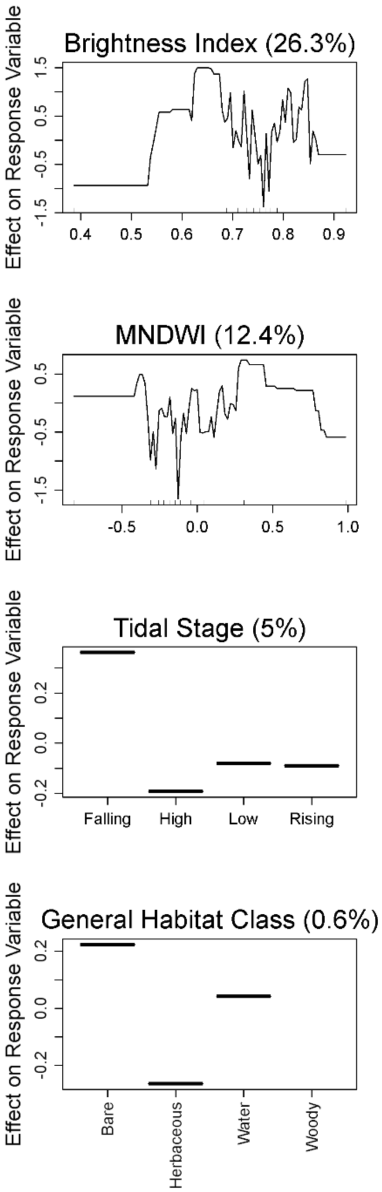
Brightness Index (26.3%)
Inundation Class (19.9%)
Distance to Water (13.5%)
MNDWI (12.4%)
MSAVI (12%)
Detailed Habitat Class (6%)
Tidal Stage (5%)
Location (3%)
Restoration Phase (1.2%)
General Habitat Class (0.6%)
Effect on Response Variable
Developed
Intertidal Unveg.
Intertidal Veg.
Supratidal
Bare Land
Beach
E.E. Marsh
Intertidal
Mangrove
Meadow
Scrb/shrb
S. Protect
U. Dune
U. Flat
V. Dune
Water
Falling
High
Low
Rising
Bay
Gulf
Interior
During
Post
Pre
Bare
Herbaceous
Water
Woody

Figure S22. Partial dependence plots of variables predicting occurrences of Piping Plover maintenance behaviors during the non-breeding seasons on Whiskey Island, Louisiana, USA from a boosted regression tree model. Y-axes are centered to have a zero mean over the data distribution. The relative influence (percent) of each predictor variable is shown in parentheses. Rug plots along the X-axis of each continuous variable plot illustrates the distribution of the data. Abbreviated labels on the X-axis of Detailed Habitat Class include Estuarine Emergent Marsh (E.E. Marsh), Scrub/Shrub (Scrb/Shrb), Shoreline Protection (S. Protect), Unvegetated Dune (U. Dune), and Vegetated Dune (V. Dune). Abbreviated labels on the X-axis of Inundation Class include Intertidal Unvegetated (Intertidal Unveg.) and Intertidal Vegetated (Intertidal Veg.). MNDWI indicates Modified Normalized Difference Water Index and MSAVI indicates Modified Soil Adjusted Vegetation Index.

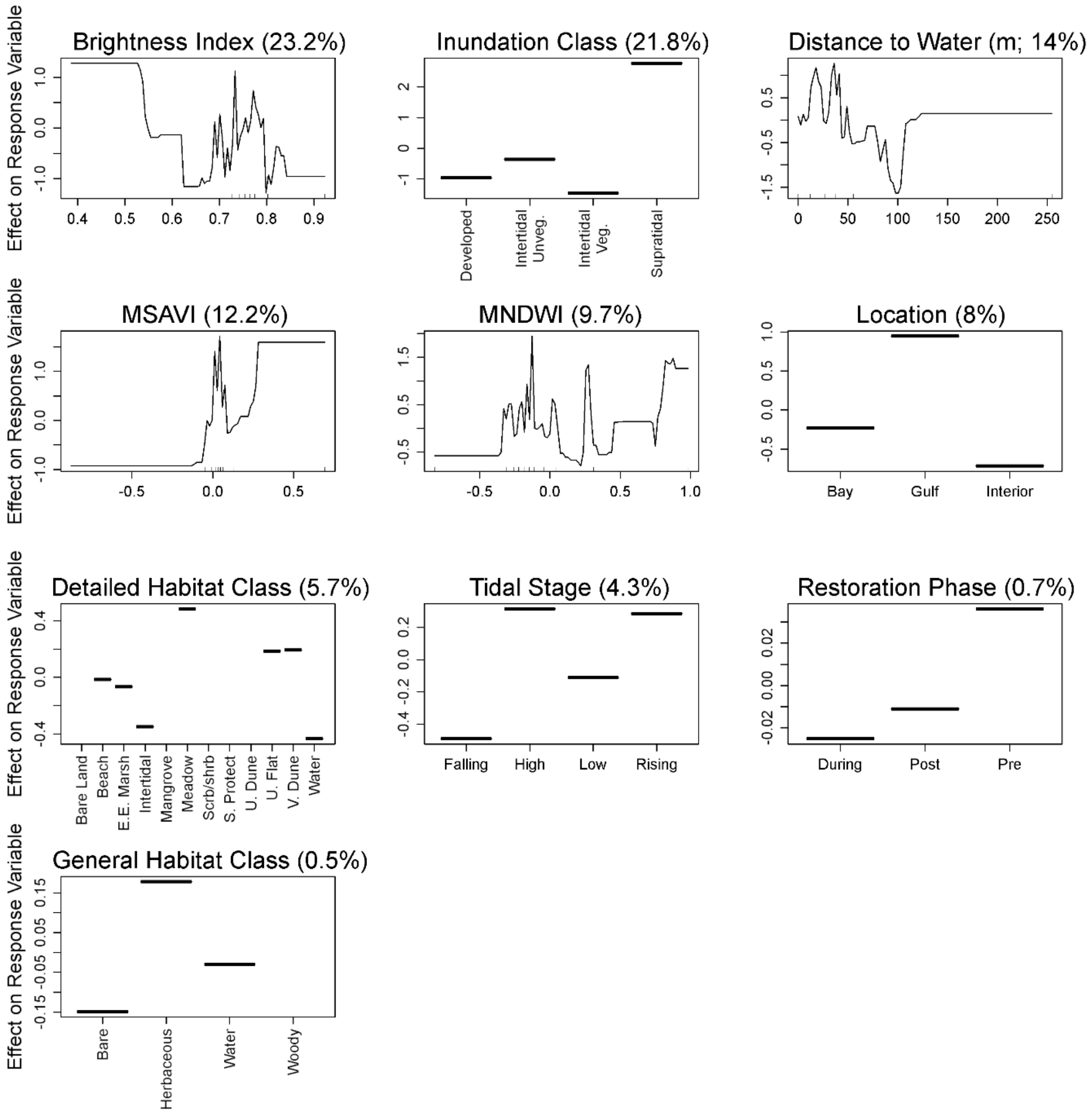
Brightness Index (23.2%)
Inundation Class (21.8%)
Distance to Water (m; 14%)
MSAVI (12.2%)
MNDWI (9.7%)
Location (8%)
Detailed Habitat Class (5.7%)
Tidal Stage (4.3%)
Restoration Phase (0.7%)
General Habitat Class (0.5%)
Effect on Response Variable
Developed
Intertidal Unveg.
Intertidal Veg.
Supratidal
Bay
Gulf
Interior
Bare Land
Beach
E.E. Marsh
Intertidal
Mangrove
Meadow
Scrb/shrb
S. Protect
U. Dune
U. Flat
V. Dune
Water
Falling
High
Low
Rising
During
Post
Pre
Bare
Herbaceous
Water
Woody

Figure S23. Partial dependence plots of variables predicting Red Knot abundance during the non-breeding seasons on Whiskey Island, Louisiana, USA from a boosted regression tree model. Y-axes are centered to have a zero mean over the data distribution. The relative influence (percent) of each predictor variable is shown in parentheses. Rug plots along the X-axis of each continuous variable plot illustrates the distribution of the data. Abbreviated labels on the X-axis of Detailed Habitat Class include Estuarine Emergent Marsh (E.E. Marsh), Scrub/Shrub (Scrb/Shrb), Shoreline Protection (S. Protect), Unvegetated Dune (U. Dune), and Vegetated Dune (V. Dune). Abbreviated labels on the X-axis of Inundation Class include Intertidal Unvegetated (Intertidal Unveg.) and Intertidal Vegetated (Intertidal Veg.). MNDWI indicates Modified Normalized Difference Water Index and MSAVI indicates Modified Soil Adjusted Vegetation Index.

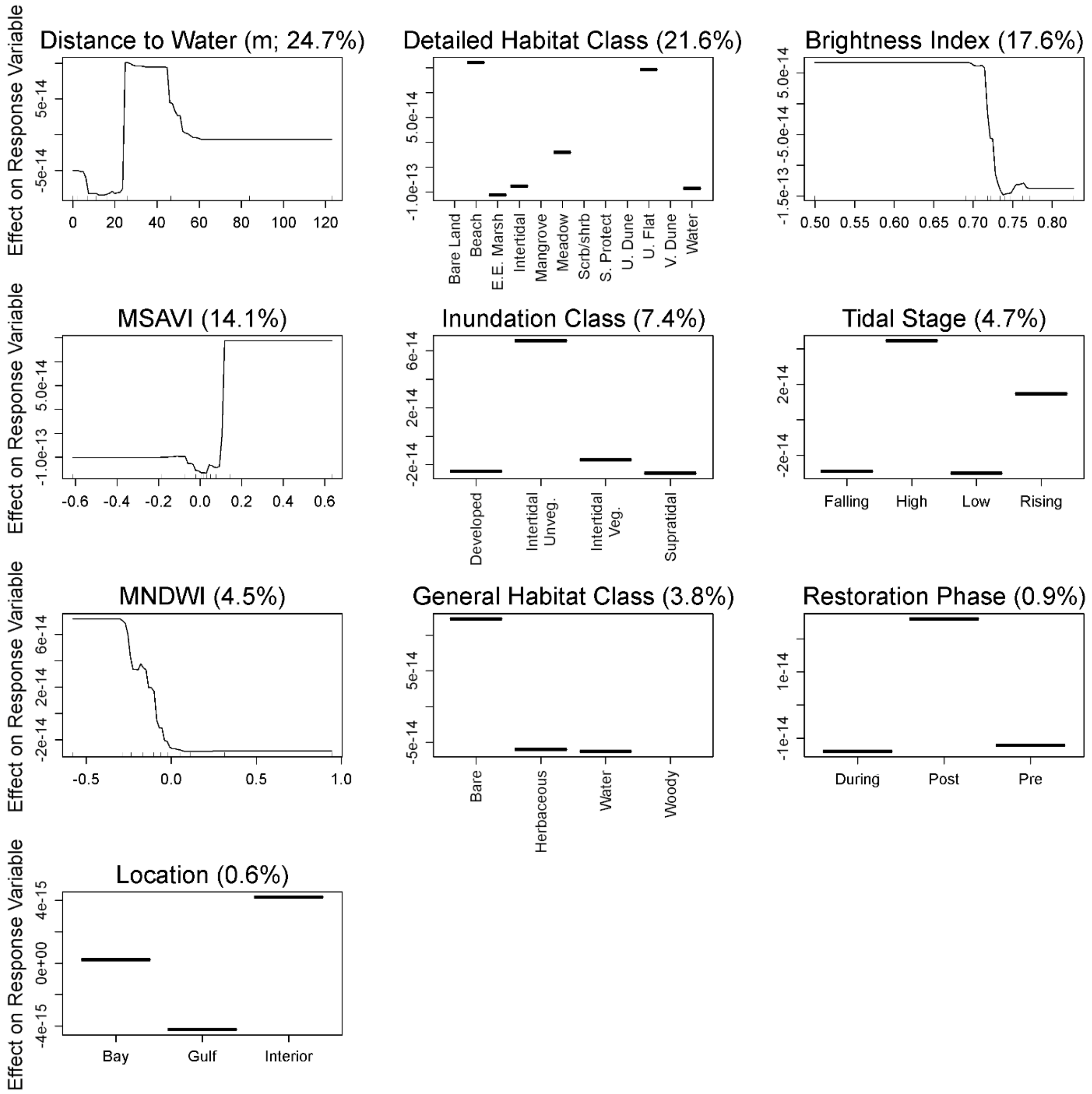
Distance to Water (m; 24.7%)
Detailed Habitat Class (21.6%)
Brightness Index (17.6%)
MSAVI (14.1%)
Inundation Class (7.4%)
Tidal Stage (4.7%)
MNDWI (4.5%)
General Habitat Class (3.8%)
Restoration Phase (0.9%)
Location (0.6%)
Effect on Response Variable
Bare Land
Beach
E.E. Marsh
Intertidal
Mangrove
Meadow
Scrb/shrb
S. Protect
U. Dune
U. Flat
V. Dune
Water
Developed
Intertidal Unveg.
Intertidal Veg.
Supratidal
Falling
High
Low
Rising
Bare
Herbaceous
Water
Woody
During
Post
Pre
Bay
Gulf
Interior

Figure S24. Partial dependence plots of variables predicting occurrences of Red Knot foraging behaviors during the non-breeding seasons on Whiskey Island, Louisiana, USA from a boosted regression tree model. Y-axes are centered to have a zero mean over the data distribution. The relative influence (percent) of each predictor variable is shown in parentheses. Rug plots along the X-axis of each continuous variable plot illustrates the distribution of the data. Abbreviated labels on the X-axis of Detailed Habitat Class include Estuarine Emergent Marsh (E.E. Marsh), Scrub/Shrub (Scrb/Shrb), Shoreline Protection (S. Protect), Unvegetated Dune (U. Dune), and Vegetated Dune (V. Dune). Abbreviated labels on the X-axis of Inundation Class include Intertidal Unvegetated (Intertidal Unveg.) and Intertidal Vegetated (Intertidal Veg.). MNDWI indicates Modified Normalized Difference Water Index and MSAVI indicates Modified Soil Adjusted Vegetation Index.

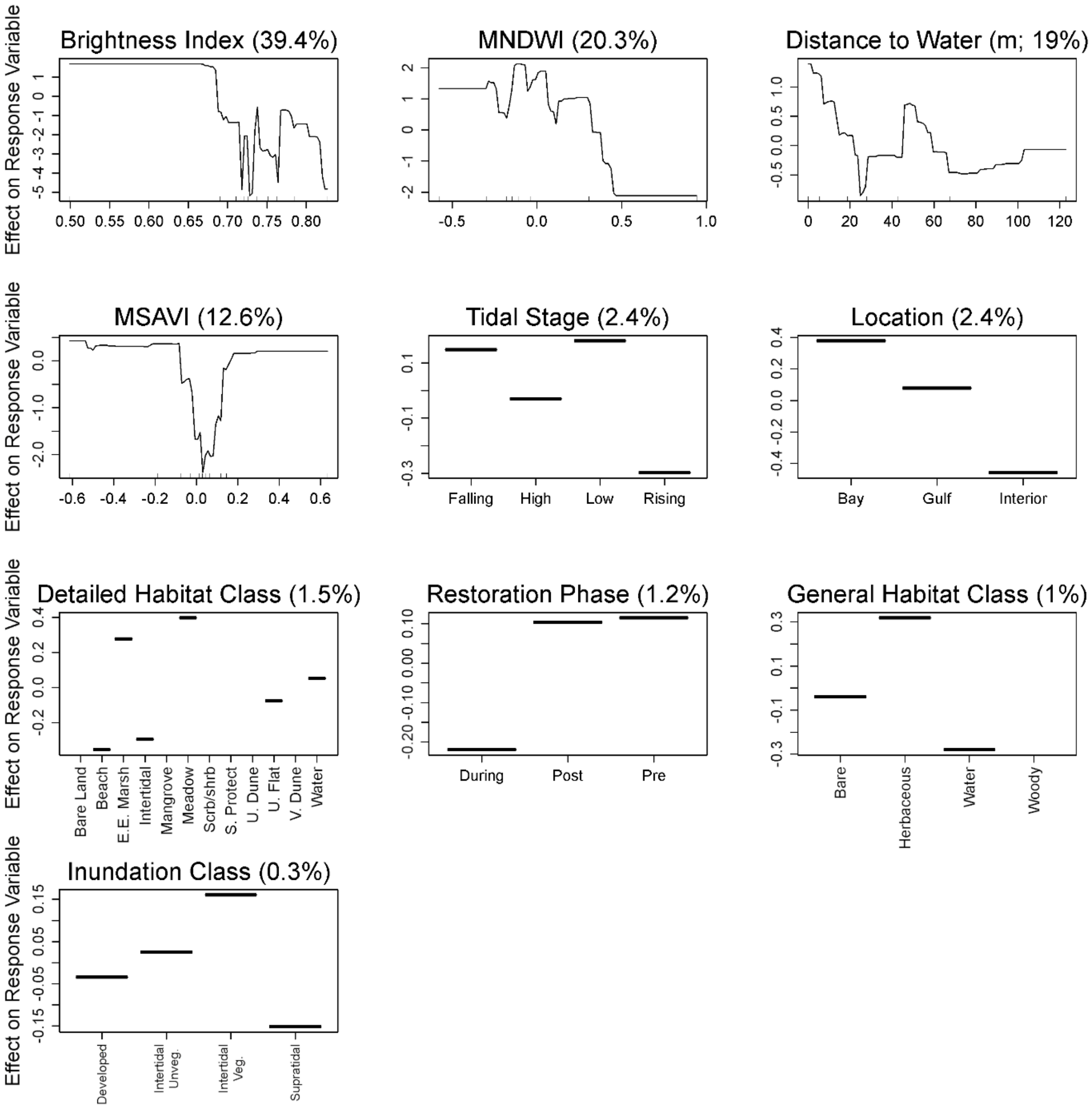
Effect on Response Variable
Brightness Index (39.4%)
MNDWI (20.3%)
Distance to Water (m; 19%)
MSAVI (12.6%)
Tidal Stage (2.4%)
Falling
High
Low
Rising
Location (2.4%)
Bay
Gulf
Interior
Detailed Habitat Class (1.5%)
Bare Land
Beach
E.E. Marsh
Intertidal
Mangrove
Meadow
Scrb/shrb
S. Protect
U. Dune
U. Flat
V. Dune
Water
Restoration Phase (1.2%)
During
Post
Pre
General Habitat Class (1%)
Bare
Herbaceous
Water
Woody
Inundation Class (0.3%)
Developed
Intertidal Unveg.
Intertidal Veg.
Supratidal

Figure S25. Partial dependence plots of variables predicting occurrences of Red Knot maintenance behaviors during the non-breeding seasons on Whiskey Island, Louisiana, USA from a boosted regression tree model. Y-axes are centered to have a zero mean over the data distribution. The relative influence (percent) of each predictor variable is shown in parentheses. Rug plots along the X-axis of each continuous variable plot illustrates the distribution of the data. Abbreviated labels on the X-axis of Detailed Habitat Class include Estuarine Emergent Marsh (E.E. Marsh), Scrub/Shrub (Scrb/Shrb), Shoreline Protection (S. Protect), Unvegetated Dune (U. Dune), and Vegetated Dune (V. Dune). Abbreviated labels on the X-axis of Inundation Class include Intertidal Unvegetated (Intertidal Unveg.) and Intertidal Vegetated (Intertidal Veg.). MNDWI indicates Modified Normalized Difference Water Index and MSAVI indicates Modified Soil Adjusted Vegetation Index.

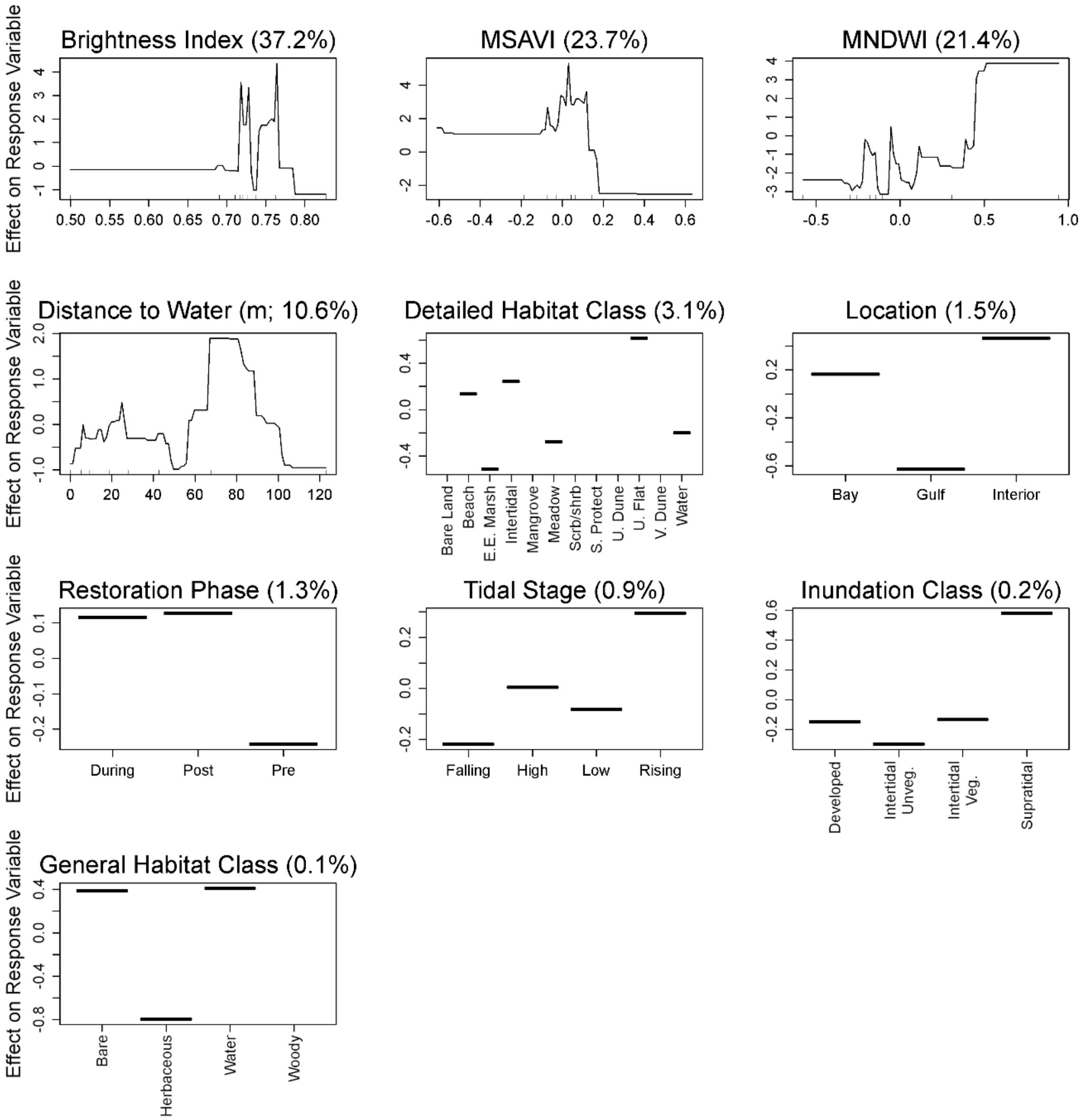

Brightness Index (37.2%)
MSAVI (23.7%)
MNDWI (21.4%)
Distance to Water (m; 10.6%)
Detailed Habitat Class (3.1%)
Location (1.5%)
Restoration Phase (1.3%)
Tidal Stage (0.9%)
Inundation Class (0.2%)
General Habitat Class (0.1%)
Effect on Response Variable
Bare Land
Beach
E.E. Marsh
Intertidal
Mangrove
Meadow
Scrb/shrb
S. Protect
U. Dune
U. Flat
V. Dune
Water
Bay
Gulf
Interior
During
Post
Pre
Falling
High
Low
Rising
Developed
Intertidal Unveg.
Intertidal Veg.
Supratidal
Bare
Herbaceous
Water
Woody

Figure S26. Partial dependence plots of variables predicting Snowy Plover abundance during the non-breeding seasons on Whiskey Island, Louisiana, USA from a boosted regression tree model. Y-axes are centered to have a zero mean over the data distribution. The relative influence (percent) of each predictor variable is shown in parentheses. Rug plots along the X-axis of each continuous variable plot illustrates the distribution of the data. Abbreviated labels on the X-axis of Detailed Habitat Class include Estuarine Emergent Marsh (E.E. Marsh), Scrub/Shrub (Scrb/Shrb), Shoreline Protection (S. Protect), Unvegetated Dune (U. Dune), and Vegetated Dune (V. Dune). Abbreviated labels on the X-axis of Inundation Class include Intertidal Unvegetated (Intertidal Unveg.) and Intertidal Vegetated (Intertidal Veg.). MNDWI indicates Modified Normalized Difference Water Index and MSAVI indicates Modified Soil Adjusted Vegetation Index.

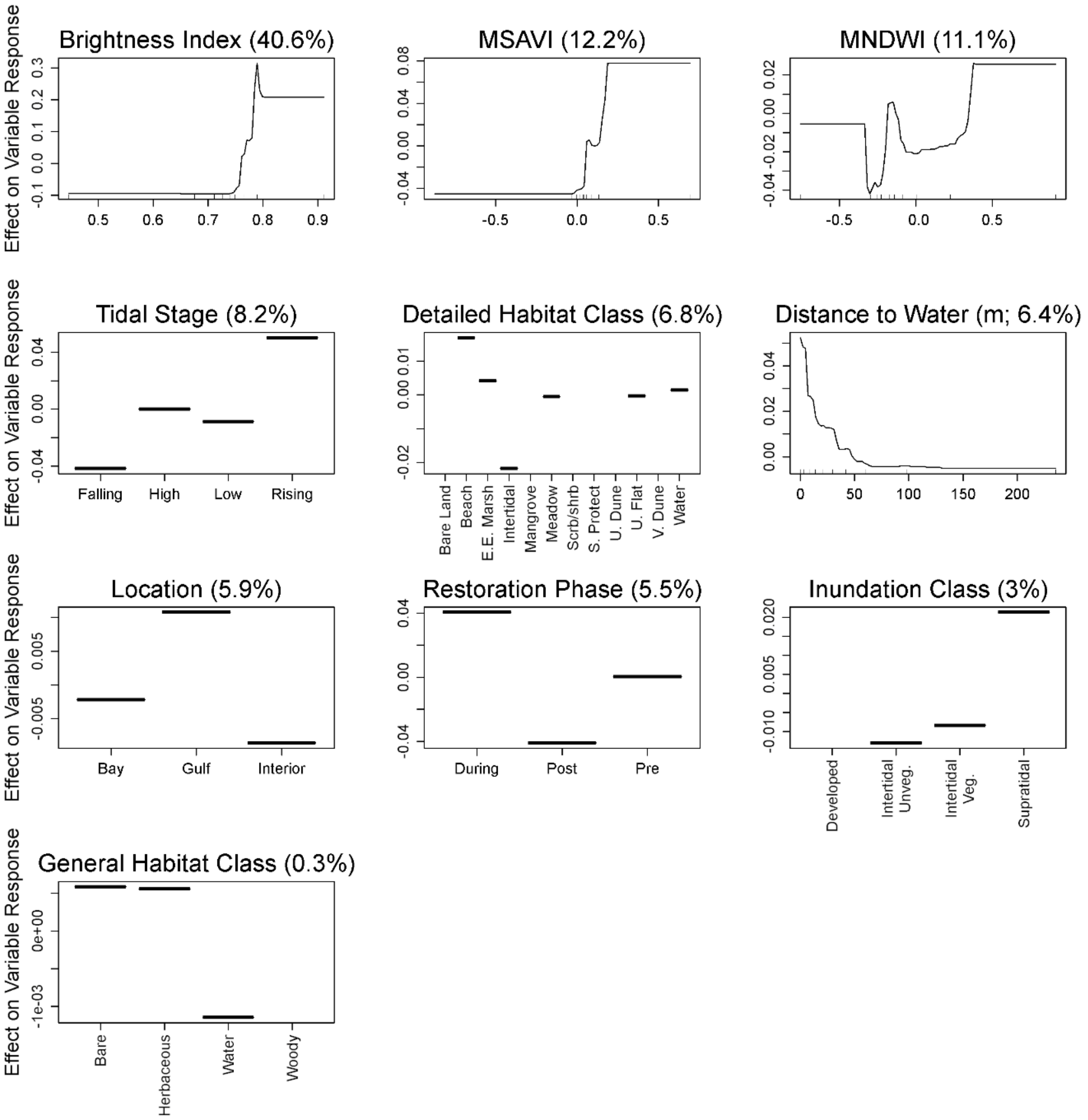
Brightness Index (40.6%)
MSAVI (12.2%)
MNDWI (11.1%)
Tidal Stage (8.2%)
Detailed Habitat Class (6.8%)
Distance to Water (m; 6.4%)
Location (5.9%)
Restoration Phase (5.5%)
Inundation Class (3%)
General Habitat Class (0.3%)
Effect on Variable Response
Falling
High
Low
Rising
Bare Land
Beach
E.E. Marsh
Intertidal
Mangrove
Meadow
Scrb/shrb
S. Protect
U. Dune
U. Flat
V. Dune
Water
Bay
Gulf
Interior
During
Post
Pre
Developed
Intertidal Unveg.
Intertidal Veg.
Supratidal
Bare
Herbaceous
Water
Woody

Figure S27. Partial dependence plots of variables predicting occurrences of Snowy Plover foraging behaviors during the non-breeding seasons on Whiskey Island, Louisiana, USA from a boosted regression tree model. Y-axes are centered to have a zero mean over the data distribution. The relative influence (percent) of each predictor variable is shown in parentheses. Rug plots along the X-axis of each continuous variable plot illustrates the distribution of the data. Abbreviated labels on the X-axis of Detailed Habitat Class include Estuarine Emergent Marsh (E.E. Marsh), Scrub/Shrub (Scrb/Shrb), Shoreline Protection (S. Protect), Unvegetated Dune (U. Dune), and Vegetated Dune (V. Dune). Abbreviated labels on the X-axis of Inundation Class include Intertidal Unvegetated (Intertidal Unveg.) and Intertidal Vegetated (Intertidal Veg.). MNDWI indicates Modified Normalized Difference Water Index and MSAVI indicates Modified Soil Adjusted Vegetation Index.

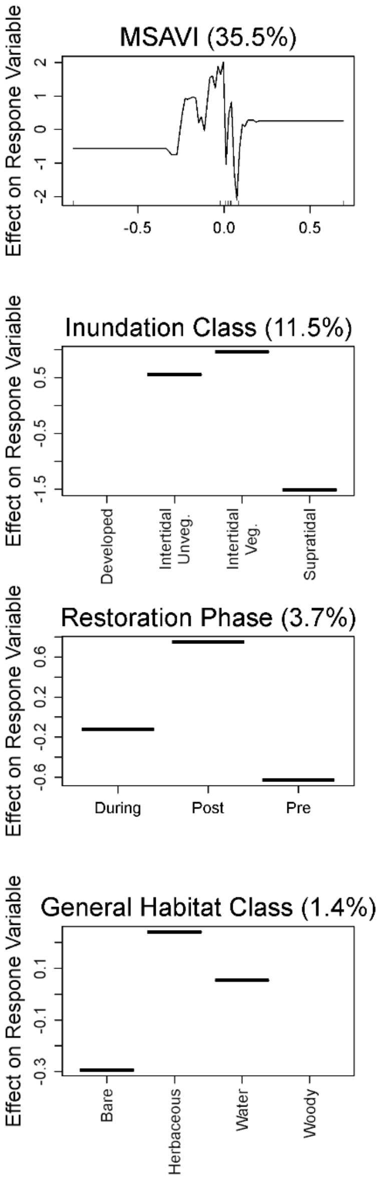

Figure S28. Partial dependence plots of variables predicting occurrences of Snowy Plover maintenance behaviors during the non-breeding seasons on Whiskey Island, Louisiana, USA from a boosted regression tree model. Y-axes are centered to have a zero mean over the data distribution. The relative influence (percent) of each predictor variable is shown in parentheses. Rug plots along the X-axis of each continuous variable plot illustrates the distribution of the data. Abbreviated labels on the X-axis of Detailed Habitat Class include Estuarine Emergent Marsh (E.E. Marsh), Scrub/Shrub (Scrb/Shrb), Shoreline Protection (S. Protect), Unvegetated Dune (U. Dune), and Vegetated Dune (V. Dune). Abbreviated labels on the X-axis of Inundation Class include Intertidal Unvegetated (Intertidal Unveg.) and Intertidal Vegetated (Intertidal Veg.). MNDWI indicates Modified Normalized Difference Water Index and MSAVI indicates Modified Soil Adjusted Vegetation Index.

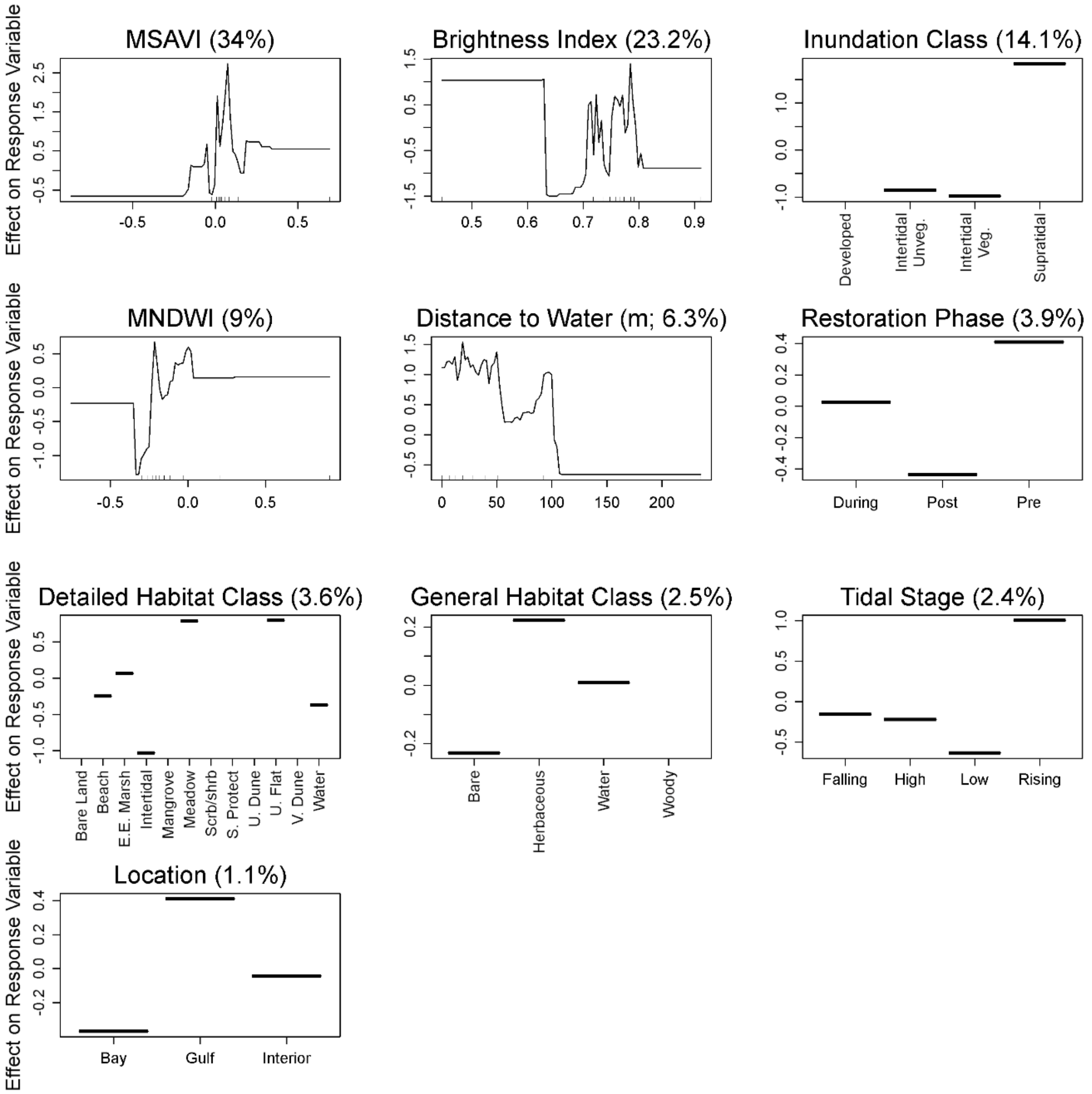

Figure S29. Partial dependence plots of variables predicting Wilson's Plover abundance during the breeding season on Whiskey Island, Louisiana, USA from a boosted regression tree model. Y-axes are centered to have a zero mean over the data distribution. The relative influence (percent) of each predictor variable is shown in parentheses. Rug plots along the X-axis of each continuous variable plot illustrates the distribution of the data. Abbreviated labels on the X-axis of Detailed Habitat Class include Estuarine Emergent Marsh (E.E. Marsh), Scrub/Shrub (Scrb/Shrb), Shoreline Protection (S. Protect), Unvegetated Dune (U. Dune), and Vegetated Dune (V. Dune). Abbreviated labels on the X-axis of Inundation Class include Intertidal Unvegetated (Intertidal Unveg.) and Intertidal Vegetated (Intertidal Veg.). MNDWI indicates Modified Normalized Difference Water Index and MSAVI indicates Modified Soil Adjusted Vegetation Index.

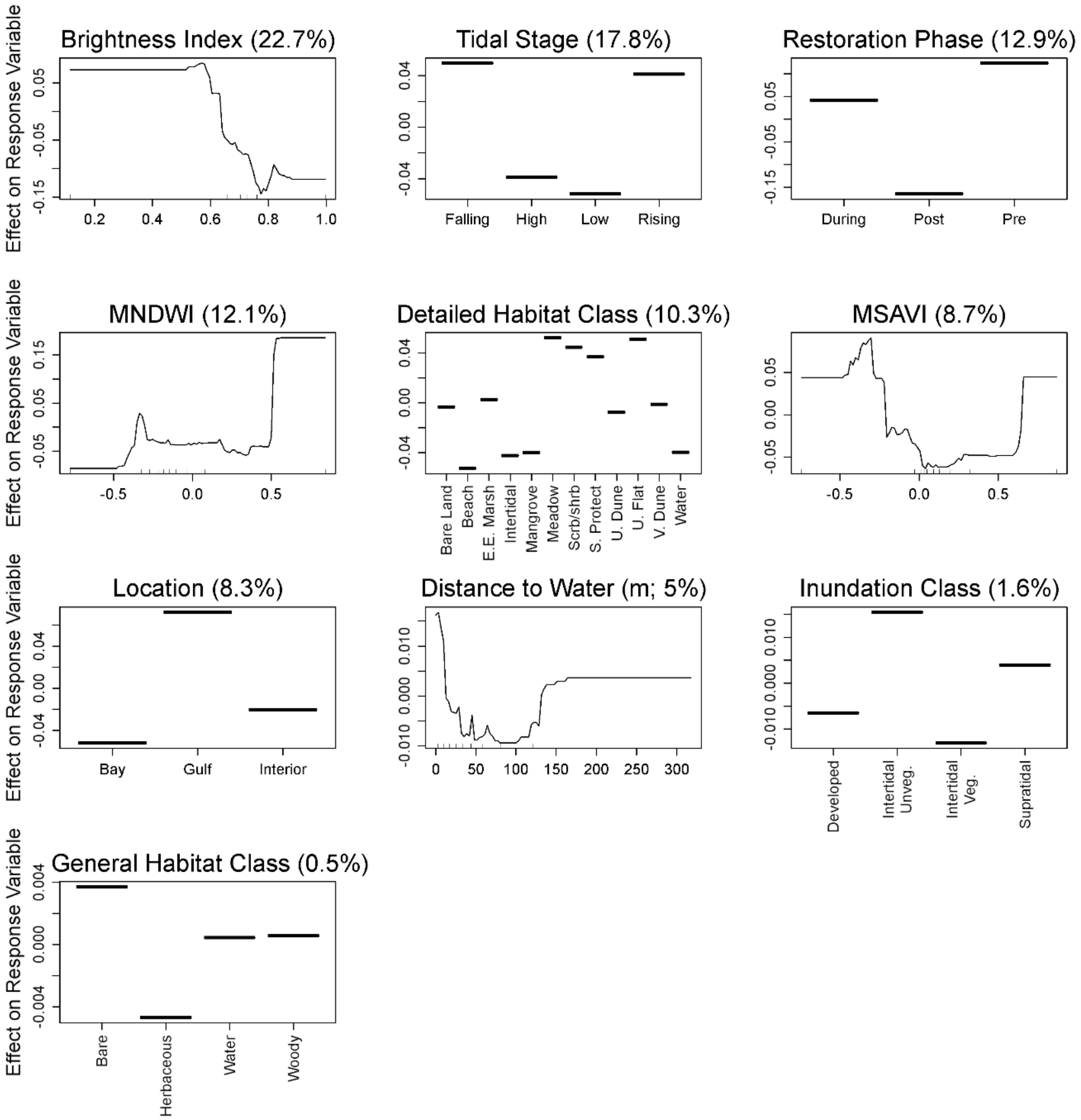

Brightness Index (22.7%)
Tidal Stage (17.8%)
Restoration Phase (12.9%)
MNDWI (12.1%)
Detailed Habitat Class (10.3%)
MSAVI (8.7%)
Location (8.3%)
Distance to Water (m; 5%)
Inundation Class (1.6%)
General Habitat Class (0.5%)
Effect on Response Variable
Falling
High
Low
Rising
During
Post
Pre
Bare Land
Beach
E.E. Marsh
Intertidal
Mangrove
Meadow
Scrb/shrb
S. Protect
U. Dune
U. Flat
V. Dune
Water
Bay
Gulf
Interior
Developed
Intertidal Unveg.
Intertidal Veg.
Supratidal
Bare
Herbaceous
Water
Woody

Figure S30. Partial dependence plots of variables predicting Wilson's Plover abundance during the non-breeding seasons on Whiskey Island, Louisiana, USA from a boosted regression tree model. Y-axes are centered to have a zero mean over the data distribution. The relative influence (percent) of each predictor variable is shown in parentheses. Rug plots along the X-axis of each continuous variable plot illustrates the distribution of the data. Abbreviated labels on the X-axis of Detailed Habitat Class include Estuarine Emergent Marsh (E.E. Marsh), Scrub/Shrub (Scrb/Shrb), Shoreline Protection (S. Protect), Unvegetated Dune (U. Dune), and Vegetated Dune (V. Dune). Abbreviated labels on the X-axis of Inundation Class include Intertidal Unvegetated (Intertidal Unveg.) and Intertidal Vegetated (Intertidal Veg.). MNDWI indicates Modified Normalized Difference Water Index and MSAVI indicates Modified Soil Adjusted Vegetation Index.

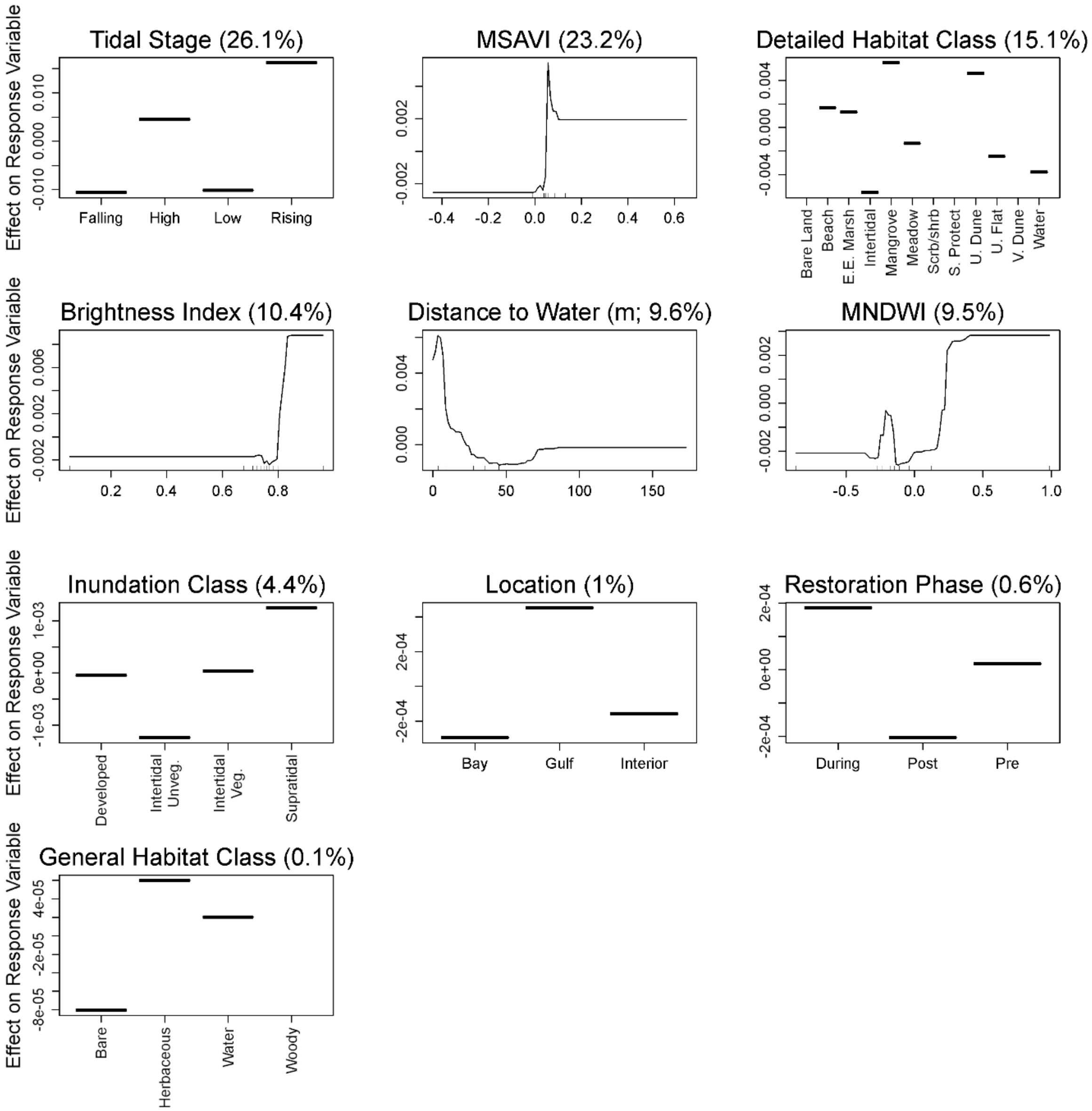
Tidal Stage (26.1%)
MSAVI (23.2%)
Detailed Habitat Class (15.1%)
Brightness Index (10.4%)
Distance to Water (m; 9.6%)
MNDWI (9.5%)
Inundation Class (4.4%)
Location (1%)
Restoration Phase (0.6%)
General Habitat Class (0.1%)
Effect on Response Variable
Falling
High
Low
Rising
Bare Land
Beach
E.E. Marsh
Intertidal
Mangrove
Meadow
Scrb/shrb
S. Protect
U. Dune
U. Flat
V. Dune
Water
Developed
Intertidal Unveg.
Intertidal Veg.
Supratidal
Bay
Gulf
Interior
During
Post
Pre
Bare
Herbaceous
Water
Woody

Figure S31. Partial dependence plots of variables predicting occurrences of Wilson's Plover breeding behaviors on Whiskey Island, Louisiana, USA from a boosted regression tree model. Y-axes are centered to have a zero mean over the data distribution. The relative influence (percent) of each predictor variable is shown in parentheses. Rug plots along the X-axis of each continuous variable plot illustrates the distribution of the data. Abbreviated labels on the X-axis of Detailed Habitat Class include Estuarine Emergent Marsh (E.E. Marsh), Scrub/Shrub (Scrb/Shrb), Shoreline Protection (S. Protect), Unvegetated Dune (U. Dune), and Vegetated Dune (V. Dune). Abbreviated labels on the X-axis of Inundation Class include Intertidal Unvegetated (Intertidal Unveg.) and Intertidal Vegetated (Intertidal Veg.). MNDWI indicates Modified Normalized Difference Water Index and MSAVI indicates Modified Soil Adjusted Vegetation Index.

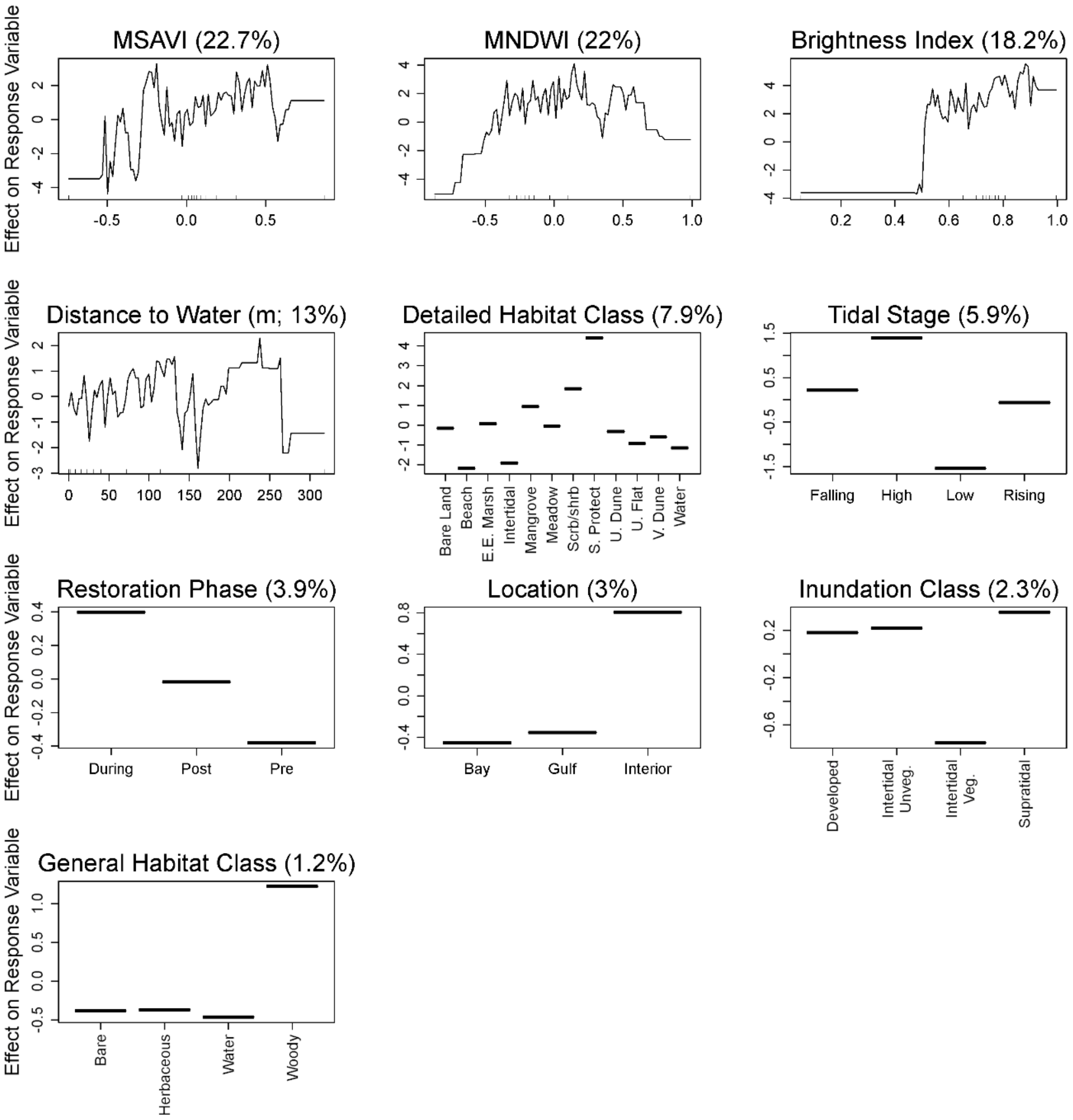

Effect on Response Variable
MSAVI (22.7%)
MNDWI (22%)
Brightness Index (18.2%)
Distance to Water (m; 13%)
Detailed Habitat Class (7.9%)
Bare Land
Beach
E.E. Marsh
Intertidal
Mangrove
Meadow
Scrb/shrb
S. Protect
U. Dune
U. Flat
V. Dune
Water
Tidal Stage (5.9%)
Falling
High
Low
Rising
Restoration Phase (3.9%)
During
Post
Pre
Location (3%)
Bay
Gulf
Interior
Inundation Class (2.3%)
Developed
Intertidal Unveg.
Intertidal Veg.
Supratidal
General Habitat Class (1.2%)
Bare
Herbaceous
Water
Woody

Figure S32. Partial dependence plots of variables predicting occurrences of Wilson's Plover foraging behaviors on Whiskey Island, Louisiana, USA from a boosted regression tree model. Y-axes are centered to have a zero mean over the data distribution. The relative influence (percent) of each predictor variable is shown in parentheses. Rug plots along the X-axis of each continuous variable plot illustrates the distribution of the data. Abbreviated labels on the X-axis of Detailed Habitat Class include Estuarine Emergent Marsh (E.E. Marsh), Scrub/Shrub (Scrb/Shrb), Shoreline Protection (S. Protect), Unvegetated Dune (U. Dune), and Vegetated Dune (V. Dune). Abbreviated labels on the X-axis of Inundation Class include Intertidal Unvegetated (Intertidal Unveg.) and Intertidal Vegetated (Intertidal Veg.). MNDWI indicates Modified Normalized Difference Water Index and MSAVI indicates Modified Soil Adjusted Vegetation Index.

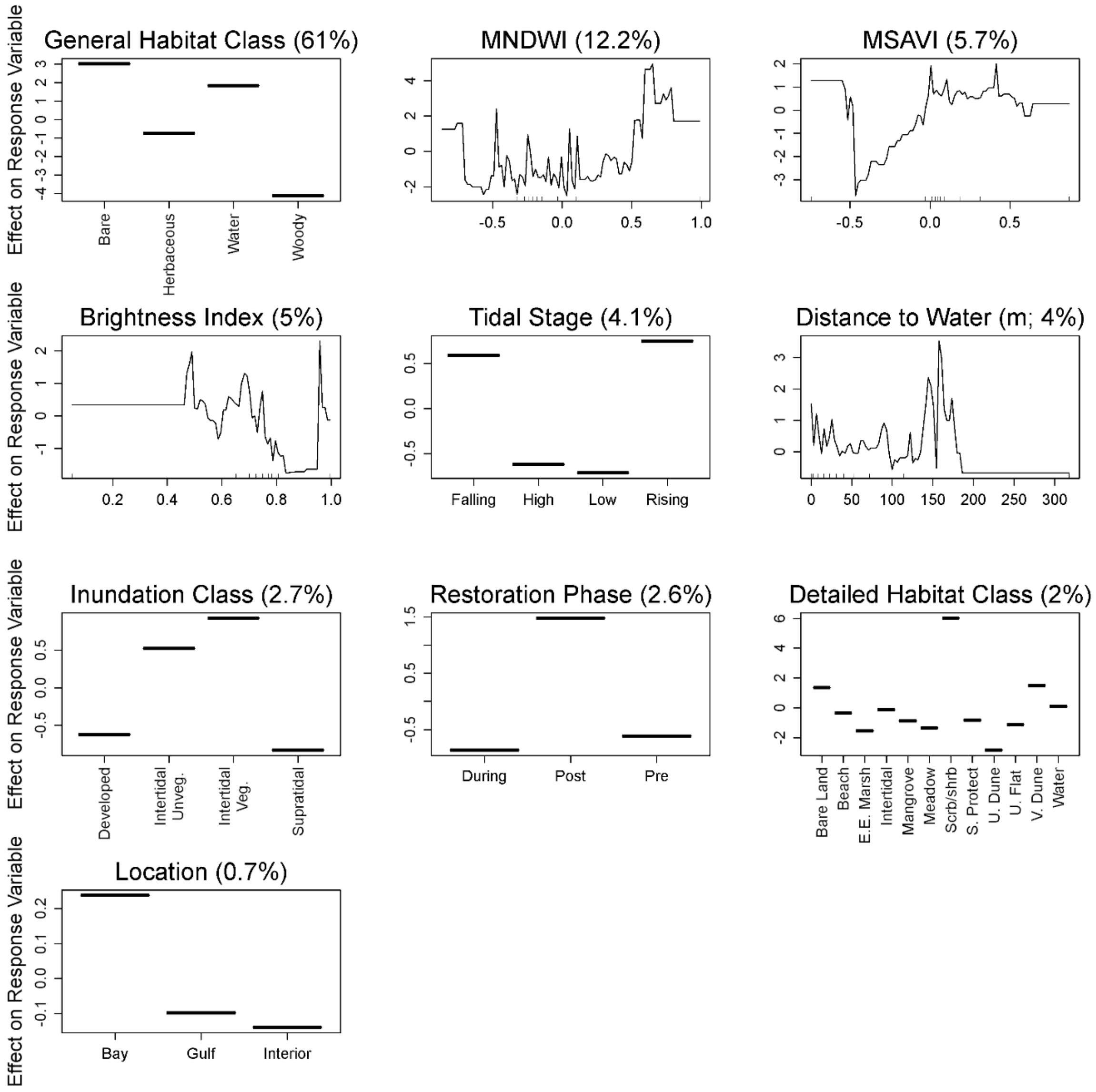
General Habitat Class (61%)
Bare
Herbaceous
Water
Woody
MNDWI (12.2%)
MSAVI (5.7%)
Brightness Index (5%)
Tidal Stage (4.1%)
Falling
High
Low
Rising
Distance to Water (m; 4%)
Inundation Class (2.7%)
Developed
Intertidal Unveg.
Intertidal Veg.
Supratidal
Restoration Phase (2.6%)
During
Post
Pre
Detailed Habitat Class (2%)
Bare Land
Beach
E.E. Marsh
Intertidal
Mangrove
Meadow
Scrb/shrb
S. Protect
U. Dune
U. Flat
V. Dune
Water
Location (0.7%)
Bay
Gulf
Interior
Effect on Response Variable

Figure S33. Partial dependence plots of variables predicting occurrences of Wilson's Plover maintenance behaviors on Whiskey Island, Louisiana, USA from a boosted regression tree model. Y-axes are centered to have a zero mean over the data distribution. The relative influence (percent) of each predictor variable is shown in parentheses. Rug plots along the X-axis of each continuous variable plot illustrates the distribution of the data. Abbreviated labels on the X-axis of Detailed Habitat Class include Estuarine Emergent Marsh (E.E. Marsh), Scrub/Shrub (Scrb/Shrb), Shoreline Protection (S. Protect), Unvegetated Dune (U. Dune), and Vegetated Dune (V. Dune). Abbreviated labels on the X-axis of Inundation Class include Intertidal Unvegetated (Intertidal Unveg.) and Intertidal Vegetated (Intertidal Veg.). MNDWI indicates Modified Normalized Difference Water Index and MSAVI indicates Modified Soil Adjusted Vegetation Index.

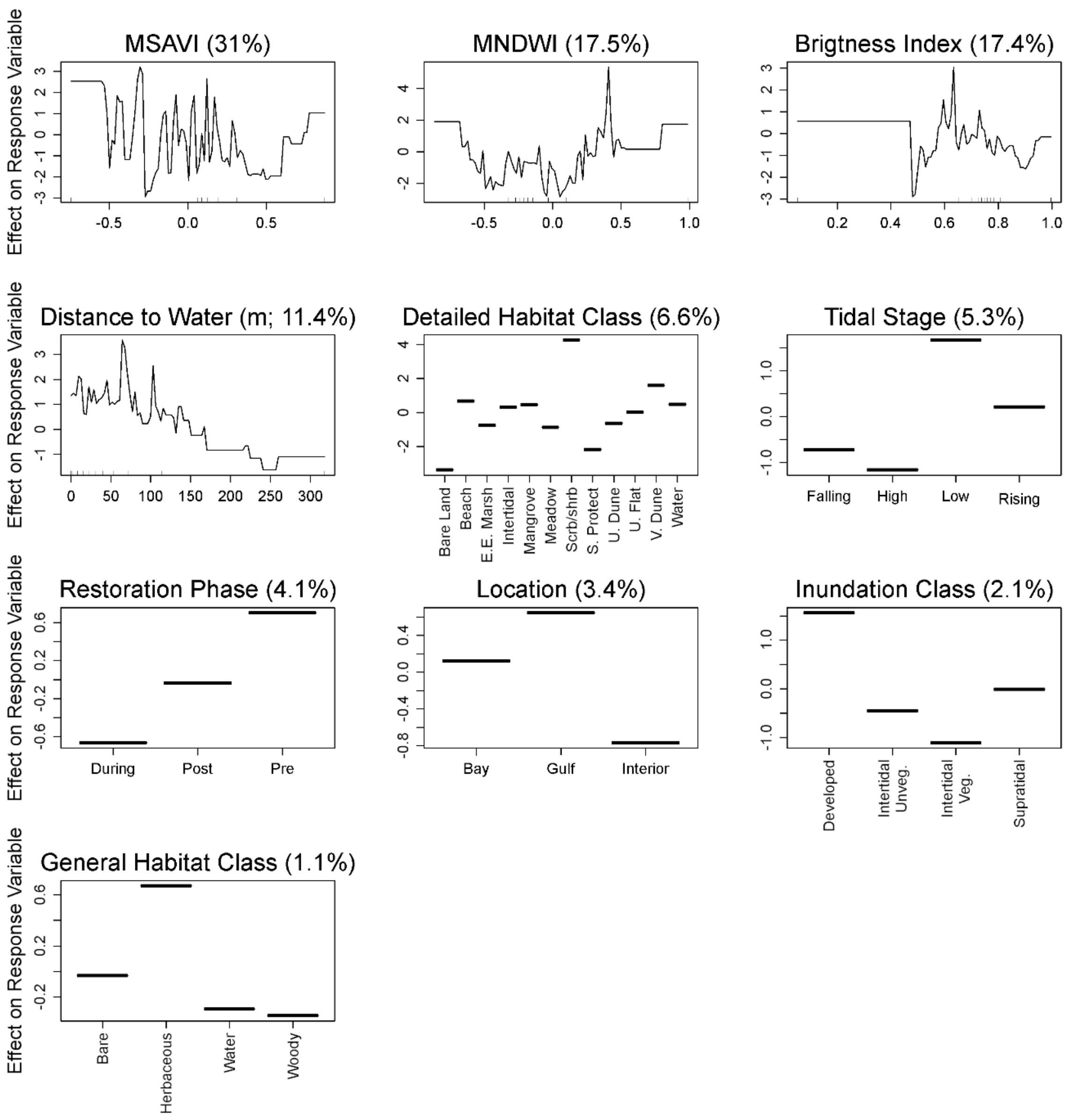

MSAVI (31%)
MNDWI (17.5%)
Brigtness Index (17.4%)
Distance to Water (m; 11.4%)
Detailed Habitat Class (6.6%)
Tidal Stage (5.3%)
Restoration Phase (4.1%)
Location (3.4%)
Inundation Class (2.1%)
General Habitat Class (1.1%)
Effect on Response Variable
Bare Land
Beach
E.E. Marsh
Intertidal
Mangrove
Meadow
Scrb/shrb
S. Protect
U. Dune
U. Flat
V. Dune
Water
Falling
High
Low
Rising
During
Post
Pre
Bay
Gulf
Interior
Developed
Intertidal Unveg.
Intertidal Veg.
Supratidal
Bare
Herbaceous
Water
Woody

Figure S34. Violin plot showing the distribution of spectral indices at Whiskey Island by sensor and index for the satellite-based maps that were assessed for accuracy (Table 6). a) Brightness index (BI) for Landsat; b) Modified normalized difference water index (MNDWI) for Landsat; c) Modified soil-adjusted vegetation index (MSAVI) for Landsat; d) BI for Sentinel-2; e) MNDWI for Sentinel-2; and f) MSAVI for Sentinel-2. On the x-axis, B = bare, HV = herbaceous vegetation, W = water, and WV = woody vegetation.

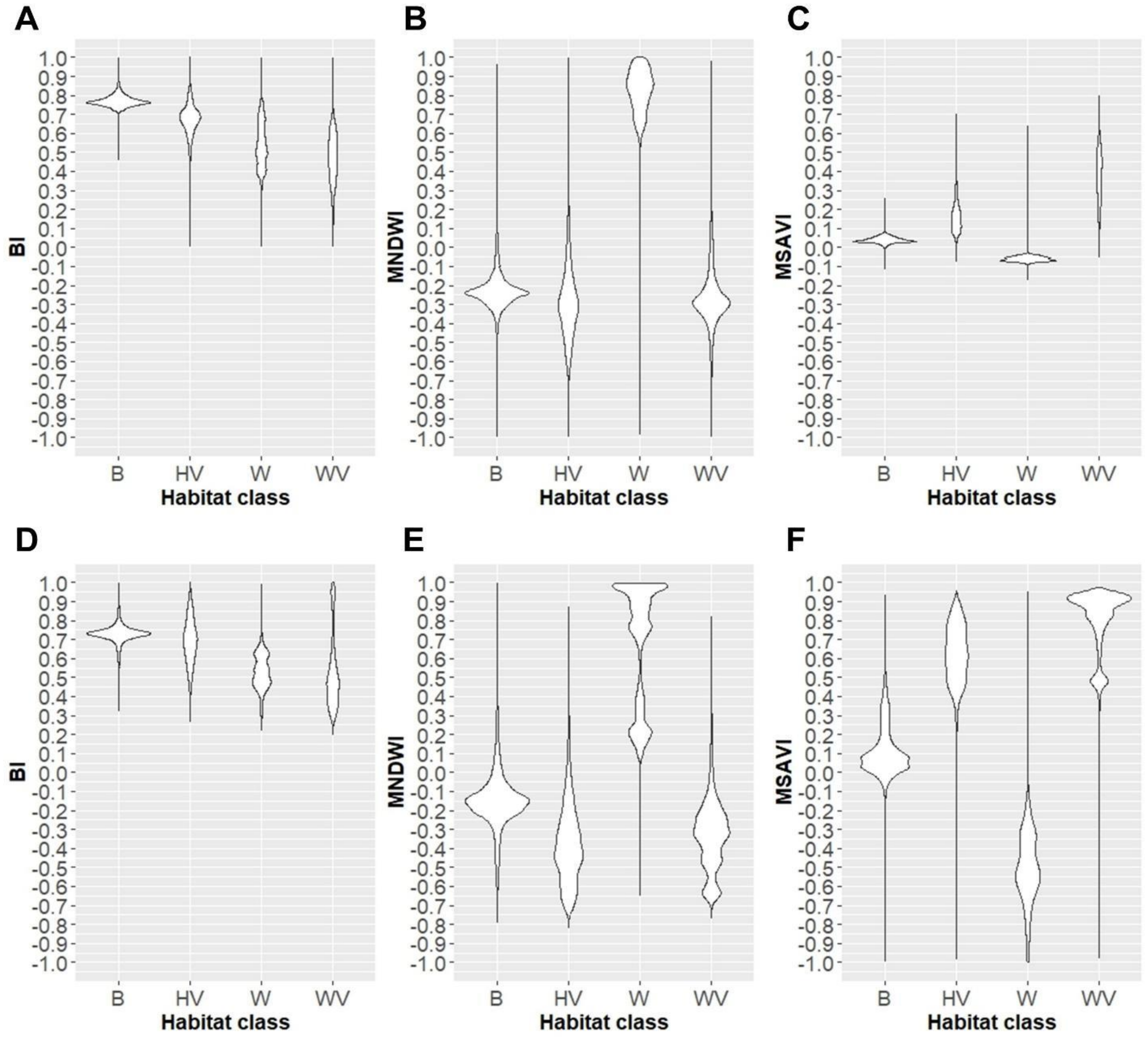

Figure S35. Violin plot showing the distribution of spectral indices at Caminada Headland by sensor and index for the satellite-based maps that were assessed for accuracy (Table 5). a) Brightness index (BI) for Landsat; b) Modified normalized difference water index (MNDWI) for Landsat; c) Modified soil-adjusted vegetation index (MSAVI) for Landsat; d) BI for Sentinel-2; e) MNDWI for Sentinel-2; and f) MSAVI for Sentinel-2. On the x-axis, B = bare, HV = herbaceous vegetation, W = water, and WV = woody vegetation.

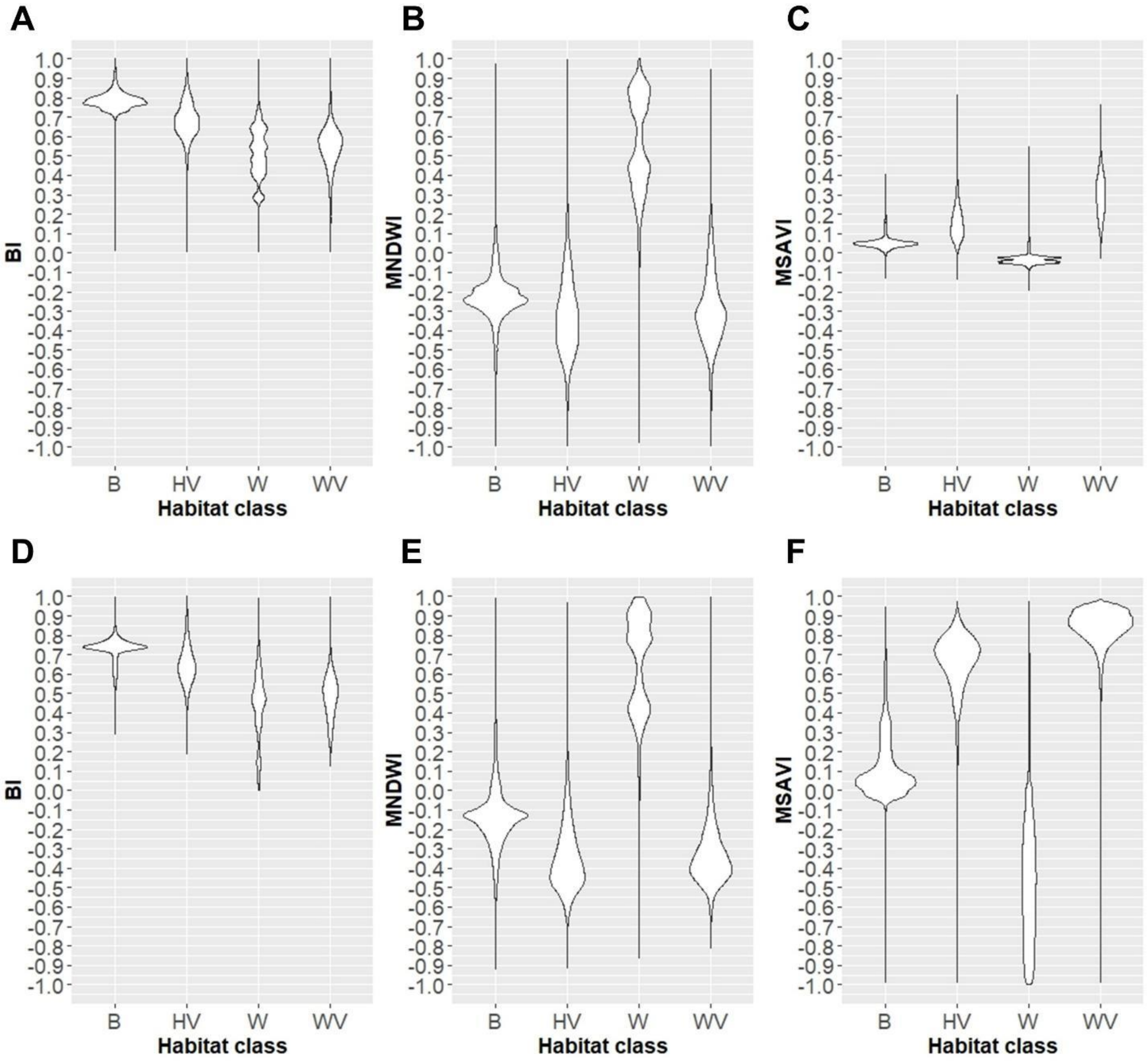
